# Nutrient signaling pathways regulate amyloid clearance and synaptic loss in Alzheimer’s disease

**DOI:** 10.1101/2020.11.13.381186

**Authors:** Mrityunjoy Mondal, Jitin Bali, Makis Tzioras, Rosa C. Paolicelli, Ali Jawaid, Martina Malnar, Kristina Dominko, Vinod Udayar, Saoussen Ben Halima, Krishna C. Vadodaria, Erica Manesso, Garima Thakur, Mickael Decressac, Constance Petit, Ravi Sudharshan, Silva Hecimovic, Mikael Simons, Judith Klumperman, Oliver Brüstle, Shawn Ferguson, Roger Nitsch, Paul E. Schulz, Tara L. Spires-Jones, Philipp Koch, Rudiyanto Gunawan, Lawrence Rajendran

## Abstract

Extra-cellular accumulation of Amyloid-β (Aβ) plaques is causatively associated with Alzheimer’s disease (AD). However, mechanisms that mediate the pre-pathological state of amyloid plaque formation remain elusive. Here, using paired RNAi and kinase inhibitor screens, we discovered that AKT-mediated insulin/nutrient signaling suppresses lysosomal clearance of Aβ and promotes amyloid formation. This mechanism is cell-autonomous and functions in multiple systems, including iPSC-derived human neurons and *in vivo.* Nutrient signaling regulates amyloid formation via distinct lysosomal functional mechanisms, while enhanced amino acid signaling promotes amyloid formation by transcriptionally suppressing lysosome biogenesis, and high intracellular cholesterol levels suppress lysosomal clearance of amyloid by increasing the number of non-functional lysosomes. The nutrient signaling pathway, present in both neurons and microglia, regulates lysosomal clearance of amyloid and microglia mediated synapse loss, both *in vitro* and *in vivo.* Clinically, older hyperlipidemic patients showed less synapse loss through microglia and performed better in cognitive tests. Thus, our results reveal a bi-partite cellular quality control system regulated by the insulin-nutrient signaling that in neurons regulates Aβ peptide clearance and in microglia regulates synaptic loss, both processes causally associated with AD. Our results also caution against reducing amyloid through such processes as this might also result in synapse loss.

## Introduction

Experimental models and data from Alzheimer’s disease (AD) patients demonstrates that Aβ deposition is preceded by pre-pathological mechanisms and understanding these would aid in early detection and intervention for AD (Jack et al., 2010; Roberts et al., 2017). While early-onset AD is primarily caused by autosomal dominant genetic mutations that increase the relative production of the toxic Aβ peptide (Aβ42) (Borchelt et al., 1996; Tanzi and Bertram, 2005; Thinakaran et al., 1996), the molecular and cellular basis of late-onset AD (LOAD), which comprises more than 98% AD patients, remains elusive. Defective Aβ clearance mechanisms perpetrated across aging are thought to contribute to amyloid accumulation (Roberts et al., 2017).

Sequential processing of the amyloid precursor protein (APP) by β- and γ-secretases produces Aβ peptides (De Strooper, 2010; Haass and Selkoe, 2007; Tanzi, 2005). Neurons generate Aβ peptides in endosomes (Grbovic et al., 2003; Rajendran et al., 2006; Udayar et al., 2013) and secrete them via exosomal and non-exosomal routes (Rajendran et al., 2006). Once released, Aβ peptides can be cleared via microglia, via the interstitial space to cerebrospinal fluid (CSF) bulk flow, efflux across the blood-brain barrier or degradation by enzymes such as insulin degrading enzyme (IDE) and Neprilysin (Howell et al., 1995; Iliff et al., 2012; Kress et al., 2014; Leissring et al., 2003; Selkoe, 2011; Weller et al., 2008). The current model posits that neurons produce and release Aβ, whose clearance is largely cell non-autonomous (through microglia, efflux out of the brain etc). This raises the question as to why neurons, which possess lysosomal machinery, do not clear the-majority of Aβ that they generate. We hypothesized that metabolic conditions may activate specific neuronal signaling pathways and inhibit such clearance mechanisms and may contribute to the pre-pathological state before Aβ accumulation occurs. In addition, we also studied the existence of such clearance mechanisms in microglia and how they would affect both amyloid levels and synapse loss and thus increase the risk for developing AD.

## Results

We previously showed that many of the LOAD risk genes identified by Genome Wide Association Studies (GWAS) do not affect the production of Aβ 42/40 peptides (Bali et al., 2012; Siegel et al., 2017). Hence we hypothesized that the signaling pathways that regulate Aβ clearance could predict late-onset Alzheimer’s Disease (LOAD) susceptibility. Cells grown on normal medium produce a large amounts of Aβ, which is a result of both production and clearance (Udayar et al., 2013). Aβ is generated in early endosomes (Ben Halima et al., 2016; Kinoshita et al., 2003; Koo and Squazzo, 1994; Rajendran et al., 2006; Small and Gandy, 2006). While much of the generated Aβ is released from cells, sporadic evidence suggests that Aβ can be degraded in lysosomes (Ben Halima et al., 2016; Buggia-Prevot et al., 2013; Glabe, 2001; Haass et al., 1992; Keilani et al., 2012; Nixon et al., 1992; Rajendran et al., 2006; Rajendran et al., 2008; Udayar et al., 2013; Xiao et al., 2015) but does not undergo this clearance under certain conditions, which could be one reason for the amyloid load seen in LOAD patients.

### Replication and validation of Aβ clearance by lysosomes

To explore Aβ clearance by lysosomes, we took two different approaches; a) to reproduce the findings that Aβ can indeed be degraded by lysosomes and b) understanding the cellular basis of inefficient lysosomal clearance under basal conditions and that may lead to amyloid formation in LOAD. First, we showed that inhibiting lysosomal acidification by chloroquine enhanced Aβ levels, confirmed that lysosomal pH and function are critical for Aβ clearance (SFigure 1A). Chloroquine treatment increased Aβ despite decreasing sAPPβ levels, consistent with the acidic environment required by β-secretase to process APP in the amyloidogenic pathway (Ben Halima and Rajendran, 2011; Rajendran et al., 2006). Secondly, we blocked endosome-lysosome fusion to prevent Aβ clearance through RNAi-mediated silencing Vamp7 and Syntaxin7, two SNARE proteins involved in endosome-lysosome fusion. This increased Aβ levels, while sAPPβ remained unaltered (SFigure 1B), indicating that Aβ degradation can occur in lysosomes and that it requires endosome-lysosome fusion.

We then tested if increasing lysosome biogenesis would clear Aβ (Parr et al., 2012). Transcription factor EB (TFEB) is a master regulator of lysosomal biogenesis (Settembre et al., 2011; Settembre et al., 2012), and it is predominantly found in the cytoplasmic compartment in its phosphorylated form. Once dephosphorylated, TFEB translocates to the nucleus and activates the expression of CLEAR (Coordinated Lysosomal Expression and Regulation) genes, which regulate autophagosome-lysosomal biogenesis. We overexpressed TFEB in HeLa SweAPP cells to increase lysosomal biogenesis and to investigate its effect on Aβ clearance. We found that overexpression of WT TFEB, and of a mutant form (TFEB-S211A/S142A) that results in constitutive localization in the nucleus significantly decreased Aβ levels (SFigure 1C, D). However, another mutant form that is lacking the nuclear localization signal (TFEB-ΔNLS), failed to significantly alter Aβ levels (SFigure 1C) (Roczniak-Ferguson et al., 2012; Settembre et al., 2012).

We next determined using a different TFEB family member, TFE3, also induces lysosome biogenesis via CLEAR gene transcription (Martina et al., 2014) and thus could modulate Aβ degradation. We found that TFE3 over expression in HeLa SweAPP cells reduced Aβ levels (SFigure 1E). Taken together, these replication data showed that the TFEB/TFE3 transcription factors that mediate lysosome biogenesis also regulate Aβ levels *in vitro.*

To validate the role of TFEB *in vivo,* we injected adeno-associated viruses (AAV) expressing TFEB or GFP into the cortex of WT mice expressing endogenous levels of APP. Brain samples were analysed 10 days later by immunostaining and Western blotting. TFEB or GFP expression was robustly induced in the injected regions (SFig.1F). Importantly, ectopic expression of TFEB significantly reduced both Aβ40 and Aβ42 levels (SFigure1G), consistent with our findings from cultured cells. The levels of LAMP1, a lysosomal membrane glycoprotein, and also of Cathepsin D, a lysosomal protease, increased in TFEB-injected regions, which suggests a role for TFEB in mediating lysosomal biogenesis through the expression of CLEAR genes (SFigure 1H) (Roczniak-Ferguson et al., 2012; Xiao et al., 2015). We further validated these findings *in vitro* showing that Cathepsin D, but not its heat-inactivated form, degrades Aβ (SFigure 1I). Thus, inducing lysosome biogenesis promoted the intracellular clearance of Aβ, both in cultured cells and *in vivo*. Taken together, these results validate the previously published work that lysosomes and the lysosomal proteases can, in principle, degrade endosomally-generated Aβ (Gouras et al., 2005; Xiao et al., 2015; Zhang and Zhao, 2015). However, under basal conditions, most of the endosomally generated Aβ seems to be secreted rather than degraded in the lysosomes.

### Insulin-AKT signaling pathway regulates intraneuronal clearance of Aβ

To understand why the bulk of Aβ is not intracellularly degraded under basal conditions, we sought to identify the signaling mechanisms that could suppress intracellular Aβ degradation and modulate its secretion. To this aim, we utilized our paired approach (RNAi and drug screen) to search for candidates that specifically affected Aβ levels without altering sAPPβ (Udayar et al., 2013). This approach focused only on the factors affecting Aβ levels via modulation of γ-secretase cleavage or via degradation. Since kinases play central functions to regulate the nuclear translocation of transcription factors involved in lysosomal biogenesis, we systematically silenced all of the 682 kinases in the human genome in HeLa-swAPP cells (Rajendran et al., 2008) and assayed the levels of secreted sAPPβ and Aβ from the conditioned media (Figure 1A). Silencing core components of the amyloidogenic machinery served as positive controls (APP, BACE1, PSEN1, PEN2) (Figure 1A). The primary RNAi screen identified key kinases involved in the insulin/IGF1/nutrient (IIN) signaling pathway, including AKT1, AKT2, PI3K, FRAP1 (mTOR), AURKAIP1 and EIF2AK1, that reduced Aβ (Figure 1A; SFigure 2) (Andersen et al., 2005; Rogaeva et al., 2007). Knockdown of AKT1 and AKT2 decreased both Aβ40 and Aβ42 (SFig.3A, B), without affecting sAPPβ levels (Figure 1A). A bioinformatics analysis of our screen also identified both the neurotrophin and insulin/IGF-1/nutrient signaling (IIN) pathways as top modulators (Figure 1C; SFigure 4A, B). Since the primary RNA screen was performed in a non-neuronal but in HeLa swAPP cell, we focused on the IIN-AKT signaling pathway with knocking down AKT2, the AKT isoform specifically involved in insulin signaling, yielding a greater reduction in Aβ40 (SFigure 3A, SFigure 5).

**Figure 1:**
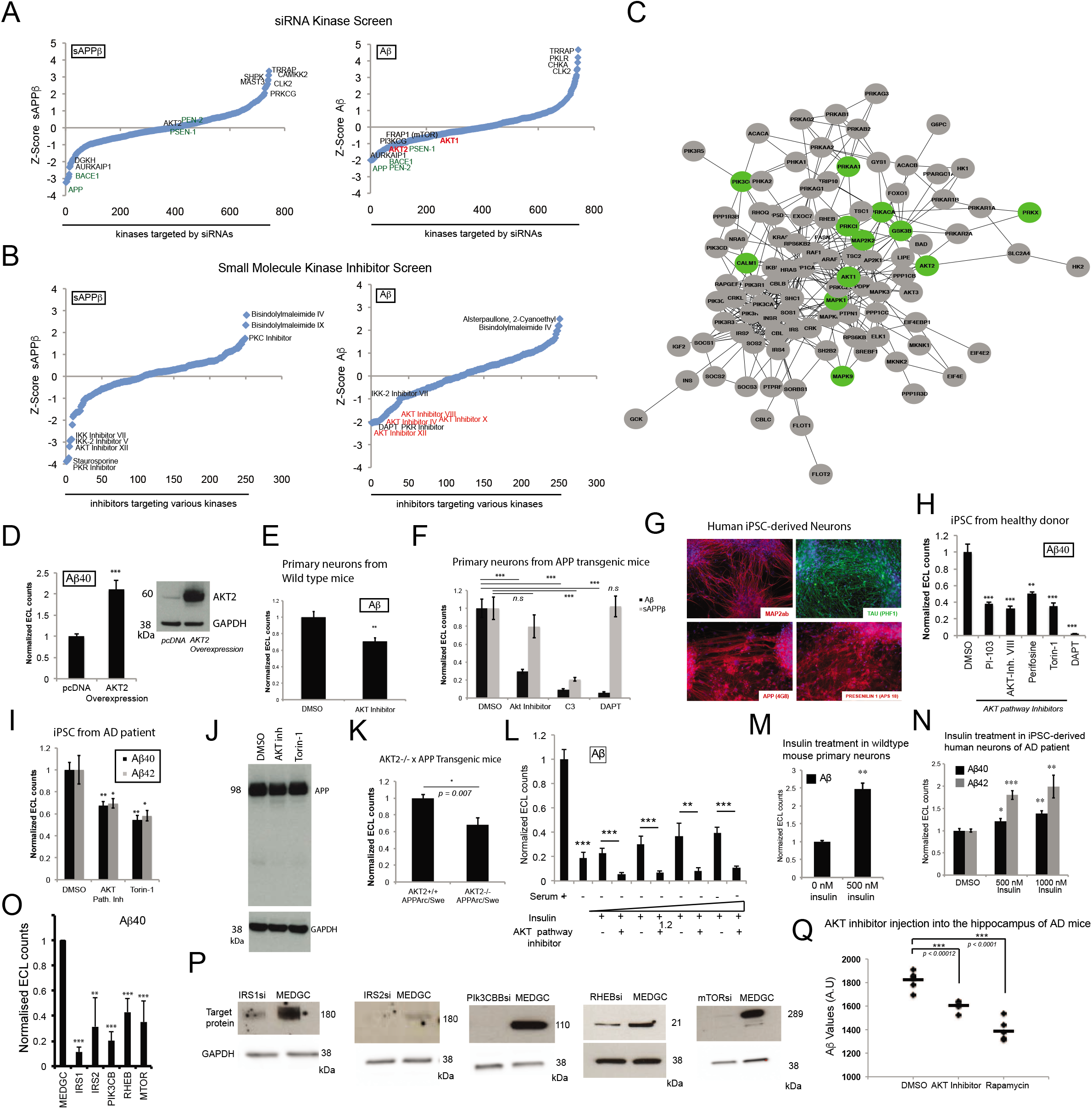
A paired RNAi and small molecule inhibitor screen of kinases using a multiplexing assay platform identify AKT as a positive regulator of Aβ levels. **A.** Graphical Z-scores (Y-axis) of the effects on sAPPβ and Aβ levels from the siRNA kinases screen. X-axis represents total number of kinases targeted by siRNA. Assay controls are indicated in green. **B**. Graphs showing Z-scores (Y-axis) of the effects on sAPPβ and Aβ levels from the kinase inhibitor/ drug screen. X-axis represents total number of inhibitors targeting various kinases. Z-Score for the screens is calculated using the formula: Z = (x – μ) / σ. Z= Z-score, x=Average value of individual sample point, μ= mean of the population, σ= Standard deviation of the population. **C.** Bioinformatics analysis of siRNA screen hits identifies various signalling networks. Insulin/nutrient signalling pathway is shown as one of the top clusters. **D.** HeLa-sweAPP cells transfected with a plasmid resulting in overexpression of AKT2 and assayed for Aβ levels. pcDNA was used as a negative control ***p<.0005. Transfection was confirmed by probing cells with AKT2 specific antibody with GAPDH being used as a loading control. **E.** Primary cortical neurons isolated from wild type mice treated with AKT inhibitor and assayed for Aβ levels. DMSO was used as a negative control, ***p<.005 (Error bars indicate SEM). **F.** Primary cortical neurons isolated from Arc/sweAPP Tg mice treated with AKT inhibitor and assayed for Aβ (black) and sAPPβ (grey) levels. DMSO was used as a negative control, DAPT and C3 treatment were used as positive controls ***p<.0005. **G, H.** Human iPSC-derived neurons treated with various AKT pathway inhibitors and analyzed for Aβ levels. DMSO was used as a negative control, DAPT treatment was used as positive control. **p<.005, ***<.0005. The accompanying immunofluorescence images demonstrate that iPSC-derived differentiated neuronal cultures mainly consist of neurons expressing MAP2ab and beta-III-tubulin (SFig.10). Expression of AD relevant proteins such as Tau, APP and Presenilin-1 were probed using indicated antibodies. **I, J.** Human iPSC-derived neurons from AD patients treated with AKT pathway inhibitors and analyzed for Aβ40 and Aβ42 levels (I) or probed with anti APP antibody (J). DMSO was used as a negative control. GAPDH was used as a loading control. *p<.05, **<.005. **K.** Primary neurons isolated from AKT2-/- x Arc/sweAPP Tg mice and AKT2+/+ x Arc/sweAPP Tg mice assayed for Aβ levels (black bars). *p<.05. **L.** HeLa-sweAPP cells serum deprived treated with insulin in the presence or absence of AKT inhibitor and assayed for Aβ levels*p<.05,** <.005,***<.0005. **M.** Primary cortical neurons isolated from wt mice treated with insulin and assayed for Aβ (black) levels,**p <.005. No insulin treatment was used as a negative control. **N.** Human iPSC-derived neurons from AD patients treated with insulin and analyzed for Aβ40 and Aβ42 levels. *p<.05,** <.005,***<.0005. **O, P.** Cells were assayed for Aβ levels upon knock down of various components of insulin signaling pathway (O). Error bars indicate S.D. Western blot analysis of insulin signaling components performed in Hela-sweAPP cells. GAPDH was used as loading control (P) **p<.005, ***p<.0005. **Q.** 3-4 month old transgenic Arc/swe APP Tg mice were stereotaxically injected with 2μl of DMSO (control), AKT inhibitor, or Rapamycin into each hippocampus and assayed for Aβ levels.

To ensure these results weren’t false positive hits, due to off-target effects, we used a complimentary approach: an unbiased screen for kinase chemical inhibitors to determine their effect on extracellular Aβ levels using 244 small molecule kinase inhibitors in the ECL-multiplexing platform. Consistent with the RNAi screen, this approach identified several inhibitors in the IIN-AKT signaling pathway that significantly decreased Aβ levels to a similar extent as the γ-secretase inhibitor DAPT (Figure 1B). The two-independent loss of function screens identified IIN-AKT signaling as a regulator of Aβ levels.

Since late-onset AD is an aging disease and there is evidence that aging modulates Aβ levels through alterations in either clearance or production, we investigated possible pathways that are enriched during aging (Mawuenyega et al., 2010). We thus analyzed human transcriptomics (RNA-seq) from the Genotype-Tissue Expression (GTEx) project and determined 2108 genes that showed differential expression (hereafter referred to as aging genes). To contextualize the aging genes to human phenotypes, we used gene ontology semantic similarity (GOSS) analysis (see supplementary material for details). Accordingly, we curated a compendium of human trait-gene associations, comprising 3358 traits and 7257 genes, from the Genome-Wide Association Study Central, Online Mendelian Inheritance in Man (OMIM) and Phenotype-Genotype Integrator (PheGenI) databases. The GOSS analysis returned a set of 51 human traits and 152 (unique) genes that are significantly similar to the ageing genes based on GO (Gene Ontology) biological processes (p < 0.05; see supplementary method). As illustrated in SFigure 4C, the Reactome pathway enrichment analysis of the aging-similar genes showed a prominent contribution from the IIN-AKT pathways among the 25 most enriched pathways (STable 1), the same pathway previously identified with using two-independent loss-of-function screens for Aβ regulation.

These independent methods ranging from loss of function screens in human cells to human tissue-specific transcriptome during aging implicated the dysregulation of IIN-AKT axis in the human aging process and possible involvement in Aβ homeostasis. Therefore, we focused on a) validating the role, b) the mechanism of IIN-AKT signaling in regulating Aβ levels and c) the relevance for AD.

As a part of the validation process, we modulated AKT levels to study how it affected Aβ. While knockdown or inhibition of AKT decreased Aβ levels, AKT2 overexpression led to a significant increase in Aβ levels (Figure 1D), ruling out any off-target effects from the loss-of-function studies. Furthermore, we also found that AKT knockdown significantly reduced not only secreted, but also intracellular Aβ levels (SFigure 6; SFigure 3A, B), indicating that AKT pathways could be involved in lysosomal clearance of Aβ prior to its release into the extracellular space. In addition, we found that the levels of APP or BACE1 (SFigure 7), amyloidogenic processing of APP (SFigure 8A, B for RNAi) and β-C-terminal fragment (CTF) (SFigure 8A-C) were unaltered by AKT knockdown. Since AKT inhibitors could thus be of therapeutic importance to reduce amyloid burden, we tested several AKT inhibitors currently in clinical trials for cancer therapeutics along with inhibitors of AKT pathway components identified in our screen. Remarkably, we found that all these inhibitors reduced Aβ levels (SFigure 9A, B). Importantly, AKT inhibition showed a concentration-dependent decrease in Aβ without affecting sAPPβ levels (SFigure 9C).

### IIN-AKT pathway regulates Aβ levels in neuronal cells, iPSC-derived human neurons and in mice

Next, we examined whether AKT modulation of Aβ levels is conserved across different cell types including human neurons. AKT inhibitors reduced levels of Aβ40 and Aβ42 in a human neuroblastoma cell line (SH-SY5Y) (SFigure 9D, E) and in primary neurons isolated either from wild-type or from Arc/SweAPP transgenic mice (Figure 1E, F). To determine if AKT inhibitors reduce Aβ levels in human neurons, we generated neuronal cultures from human induced pluripotent stem cells (iPSC) derived from two AD patients and control subjects (SFigure 10A, B; Figure 1G). AKT pathway inhibitors significantly decreased Aβ levels without influencing APP processing in human iPSC-derived neurons from AD and control patients (Figure 1H-J). These data demonstrate that Insulin-AKT signaling positively regulates Aβ levels in human neurons.

Finally, to study the *in vivo* significance, we crossed Arc/SweAPP transgenic mice, which have increased Aβ levels, with AKT2 knockout mice. Aβ levels in both primary neurons and brain extracts showed a significant reduction (Figure 1K; SFigure 11) consistent with *in vitro* findings, results which suggest that AKT signaling regulates Aβ levels both *in vitro* and *in vivo*.

### IIN-AKT pathway alters Aβ clearance without affecting Aβ production

After demonstrating that the AKT signaling pathway modulates Aβ levels, we investigated the mechanism through which this occurs. Given that APP and sAPPβ levels were not affected by AKT pathway inhibition (Fig.1A, B) we could rule out an effect on APP processing and β-secretase-mediated Aβ production. It was predicted that there are two ways through which AKT inhibition could alter Aβ levels – modulating γ-secretase cleavage or regulating Aβ clearance. We also excluded that AKT activity affects APP or BACE1 levels (SFigure 7), amyloidogenic processing of APP (SFigure 8A, B for RNAi), β-C-terminal fragment (CTF) (SFigure 8A, B) or influences γ-secretase activity (SFigure 8C). These results suggest that AKT inhibition acts independent of Aβ production.

Then we focused our attention on how AKT signaling regulated Aβ clearance. While extracellular Aβ clearance could occur via Aβ-degrading enzymes, we found no evidence that AKT inhibition influenced enzymatic degradation of Aβ (SFigure 12A) nor enhanced paracrine uptake and degradation (SFigure 12B). To rule out that the observed decrease in Aβ was due to enhanced protein secretion, we quantified the secretion of an unrelated protein, secreted alkaline phosphatase (SEAP), upon depletion of AKT1 or AKT2, and found no effects (SFigure 12C).

We found that treatment of cells with insulin, in the presence of nutrients, increased intracellular and secreted Aβ levels with no changes in APP, which was mediated by AKT (Figure 1L; SFigure 13B, C). Similarly, insulin treatment increased Aβ levels in primary mouse neurons (Figure 1M) and iPSC-derived human neurons (Figure 1N) without altering APP levels (SFigure 13A). Silencing the major signaling genes in insulin-AKT signaling pathway by RNAi (IRS1, IRS2, PI3K, mTOR and RHEB) reduced Aβ levels (Figure 1O, P). We also validated the role of insulin-AKT signaling on Aβ levels *in vivo* by stereotactically injecting an AKT inhibitor into the hippocampi of Arc/SweAPP Tg mice. We found significantly decreased Aβ levels compared to DMSO-treated controls (Figure 1Q). It has been shown that insulin delivery in the hippocampal led to increase in ISF Aβ levels (Stanley et al., 2016). These results indicate that insulin-AKT signaling positively stimulates Aβ accumulation but does not affect γ-secretase-mediated Aβ production, extracellular Aβ degradation or overall protein secretion. Thus, we hypothesized that the observed Aβ reduction following AKT inhibition could be elicited through enhanced intracellular Aβ clearance prior to release, possibly via lysosomes as the peptide is generated in endosomes (Edgar et al., 2015; Gouras et al., 2005; Morel et al., 2013; Nixon and Cataldo, 2006; Nixon et al., 2001; Rajendran et al., 2006; Takahashi et al., 2004; Tampellini et al., 2007; Udayar et al., 2013). Therefore, we speculated that the insulin-AKT signaling pathway alters intracellular lysosomal Aβ clearance, which has been previously linked but it was never experimentally demonstrated as to why it is not functional during basal conditions.

Since insulin and AKT affect lysosomal clearance of Aβ, we investigated the mechanism that links insulin-AKT and lysosomes. Given the role of TFEB to mediate Aβ clearance via lysosomes (SFigure 1D-G), we hypothesized that mTOR activation by insulin-AKT signaling regulates Aβ levels via TFEB phosphorylation and cytosolic retention to prevent lysosome biogenesis. TFEB phosphorylation is dependent on mTOR complex 1 (mTORC1), which is primarily localized to the lysosomal/late endosomal membrane (Puertollano et al., 2018; Roczniak-Ferguson et al., 2012). However, in the absence of insulin/nutrient signaling, mTOR is inhibited by TSC2 (Demetriades et al., 2014; Menon et al., 2014), an AKT substrate, that prevents its localization at the lysosomal membrane. Insulin-AKT signaling triggered TSC2 phosphorylation (Figure 2A), which displaces it from the lysosomal surface (Figure 2B). This displacement increases mTOR localization on the lysosomal membrane (SFigure 14A), and its activation, as confirmed by phosphorylation of known mTOR substrates S6K and 4EBP (Figure 2C).

**Figure 2:**
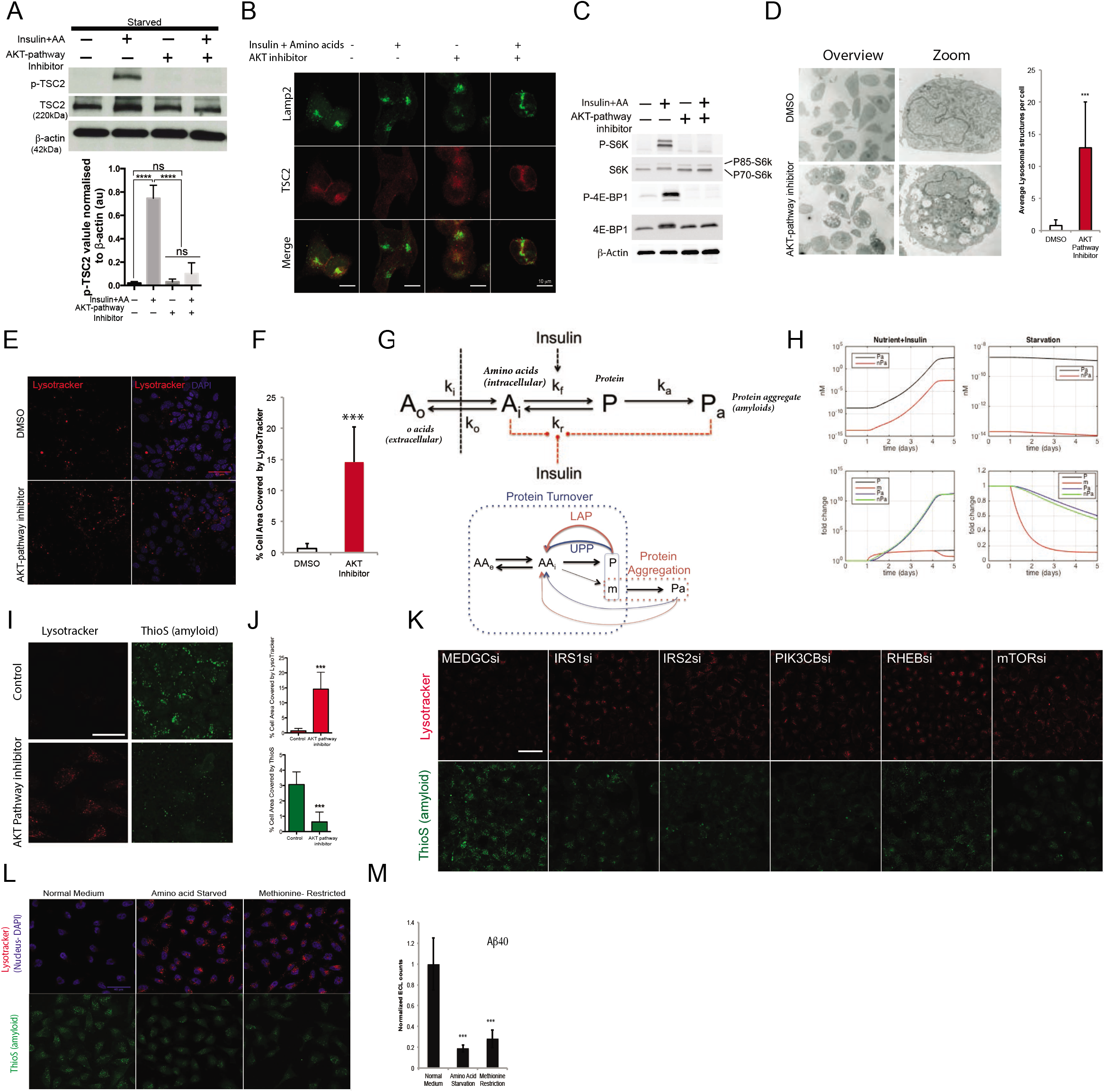
AKT inhibition decreases Aβ levels through mTOR-TFEB mediated induction of lysosomal/autophagosomal biogenesis. Increase of Lysosomal levels due to fasting, starvation or blocking IIN pathway leads to decrease of protein aggregates. **A, B, C.** HEK293 cells were serum starved for 10 h followed by amino acid starvation for 1h. Cells were then treated with AKT pathway inhibitor (10 μM) for 2.5 h. For insulin treatment, cells were pretreated (2h) with an AKT pathway inhibitor and then stimulated with insulin (1 μM) along with/without AKT pathway inhibitor for 30 min. DMSO treatment was used as negative control. p-TSC2 and total TSC2 amounts were determined by western blot with β-actin being used as a loading control. Bars indicate amount of p-TSC2 and total TSC2 normalised to β-actin (A). Cells were co-labeled for Lamp2 (green) and TSC2(red)(B). Scale bar is 10 μm. **C.** p-S6K, total S6K, p-4E-BP1 and 4E-BP1 amounts were determined by using specific antibodies. β-actin was used as a loading control. **D.** Electron microscopy images of cells treated with AKT inhibitor and DMSO (negative control). Quantification of vacuolar structures (lysosomal structures) after treatment with AKT pathway inhibitor. DMSO treatment was used as control. **E.** Primary neurons isolated from WT mice, treated with AKT pathway inhibitor and probed for lysotracker (red) and DAPI for nuclei. DMSO treatment was used as a negative control. Scale bar is 40 μm. **F.** The graph shows the percentage of cell area covered by Lysotracker positive puncta in cells treated with AKT pathway inhibitor (n = 38) vs DMSO control (n = 25). ***, P<0.0001. **G.** Mathematical model for protein aggregation. **H.** Simulated effects of increased nutrition and insulin (left column) and of starvation (right column) on the aggregation of WT Aβ. The treatment simulations started at the beginning of day 2. Increased nutrition (starvation) corresponds to three times higher (lower) extracellular amino acids. Insulin increases protein synthesis (by 5 times) and lowers the autophagosomal-lysosomal degradation pathway (by 80%) while upregulating ubiquitin-proteasomal degradation pathway (by 30%). The simulations were performed using protein aggregation kinetics of Aβ42 (Meisl *et al.,* 2014). **I.** Cells treated with AKT pathway inhibitor over night showed increase of lysotracker (red) positive structure and correlated with reduction of ThioS (green) positive structure. Scale bar is 40μm. **J.** Quantification of Lysotracker (red) and ThioS (green) signal of fig I. Error bars indicate S.D. ***p<.0005. **K.** Cell were stained with ThioS (green) or Lysotracker (red) upon knock down of various components of insulin signaling pathway. Scale bar is 40μm. **L, M.** Cells were subjected to amino acid and methionine starvation for 3 hr. After treatment cells were stained with lysotracker (red), ThioS (green) and DAPI (blue); Scale bar is 40μm. (L) or assayed for Aβ levels (M) Error bars indicate S.D. ***p<.0005.

Inhibition of insulin-AKT signaling significantly reduced TFEB phosphorylation (SFigure 14B), promoted nuclear localization of TFEB-GFP (SFigure 14C) and induced lysosome biogenesis, as confirmed by electron microscopy, confocal imaging and bright field imaging analysis (Figure 2D-F), with such increases possibly explaining previously described reduction in Aβ levels (SFigure 3A, B; SFigure 9A, B). Importantly, AKT inhibition induced lysosomal biogenesis and increased CLEAR gene expression in primary neurons (SFigure 15). Based on our data, we propose that insulin-AKT signaling positively regulates mTOR activation in a TSC2-dependent manner, which suppresses TFEB-mediated transcription of lysosomal biogenesis genes. This suppression of lysosome biogenesis results in decreased clearance of Aβ.

### Mathematical modelling the roles of insulin and nutrients on amyloid formation

Under active insulin/nutrient (amino acids) signaling conditions, cells induce protein synthesis and suppress degradative mechanisms, such as lysosomal and autophagic clearance with previous studies demonstrating that IIN signaling inhibits autophagy (Blommaart et al., 1995a; Blommaart et al., 1995b; Kim et al., 2011). Our results now show that this signaling also inhibits lysosomal biogenesis without significantly modulating proteasomal degradation, the other major cellular degradative pathway (Zhang et al., 2014) (SFigure 16A). We hypothesized then, that insulin-AKT signaling would inhibit both lysosomal and autophagosomal function and stimulate protein translation to increase protein load. The net intracellular protein concentration, where amyloid-prone proteins like Aβ likely aggregate, should reflect this increase (SFigure 16B). This implies that limiting insulin or nutrient intake should enhance lysosomal clearance of amyloid and that while early-onset AD mutations would increase the production of aggregation-prone amyloidogenic Aβ species, amyloid accumulation in late-onset AD could occur by modulating IIN-AKT signalling (hereafter termed IIN for Insulin-IGF1-Nutrient) or nutrient signalling.

While we understand how mutations in APP and Presenilins could enhance amyloid load in Alzheimer’s patients, we examined how IIN signaling affects amyloid formation as a putative mechanism active in late-onset AD. We developed a mathematical model to predict the effect of IIN signaling and nutrient overload or nutrient limitation (fasting) on amyloid levels. In this model, we incorporate protein turnover, amino acid transport across the cell membrane and Aβ monomer aggregation based on the mass balance of amino acids, proteins and protein aggregates (Figure 2G) (see Supplementary Material for more details). We estimated protein turnover kinetics by model fitting and adapted protein aggregation kinetics from a previous model of Aβ aggregation (Meisl et al., 2014) (Figure 2H). We assumed that Aβ monomer synthesis constitutes a fraction (10^-6^) of total protein synthesis, and degradation of protein aggregate is less efficient than proteins by a factor of 10. IIN signaling regulates the balance between protein production and degradation, with higher IIN signaling favouring protein synthesis.

Using this model, we investigated the effects of higher insulin and nutrient levels and caloric restriction (starvation) on the concentrations of protein (P), Aβ monomers (m), the number (nPa) and mass (Pa) concentration of protein aggregates. As shown in SFigure 16C, increased nutrition and insulin significantly increased protein concentration, including protein monomers, and increased protein aggregate number and size (average size ~ Pa/nPa). Caloric restriction caused the opposite, albeit weaker effects, with lowered protein concentration and aggregate formation being determined. We also studied mutations that increase the rates of aggregate formation and growth (Chiti et al., 2003). As shown in Figure 2H, mutant Aβ aggregated rapidly, reaching a plateau when monomer concentration limited new aggregate formation. This mutation did not affect the degradation kinetics of Aβ aggregates, so the effects of caloric restriction closely followed those of wild-type proteins. This model demonstrates that caloric restriction benefits both autosomal dominant and late-onset AD. Our mathematical model predicts that positive insulin/nutrient signaling could promote amyloid formation and that limiting this signaling could reduce amyloid load.

### Experimental evidence for the roles of insulin and nutrients (amino acids) on amyloid formation

Next, we experimentally tested the prediction that insulin/nutrient signaling induces amyloid formation by inhibiting lysosome biogenesis. We treated cells with or without an AKT inhibitor in the presence of insulin and amino acids and probed the aggregates using Thioflavin S (ThioS), a dye routinely used to detect amyloid through binding to its β-sheets, not just Aβ but all aggregates that are amyloid in nature. After optimizing our ThioS parameters, we demonstrated that ThioS detects intracellular amyloids (Figure 2I). Similar to our earlier results, cells treated with insulin contained lower levels of lysosomes but also showed higher amyloid levels (ThioS puncta). However, cells treated with an AKT pathway inhibitor had higher lysosome levels and lower amyloid levels (Figure 2I, J). We further evaluated the major constituents in IIN signaling mechanisms by RNAi (IRS1, IRS2, PI3K, mTOR and RHEB). Silencing of these genes also induced lysosome biogenesis and reduced Aβ levels and ThioS-positive amyloid puncta (Figure 1O, P; 2K). These results clearly demonstrate that positive IIN signaling decreases lysosome levels and promotes amyloid formation.

We then tested whether inhibition of insulin signaling and amino acid starvation increased lysosome levels, and whether this resulted in decreased amyloid load. We restricted cells of either all amino acids or methionine, an essential amino acid linked to longevity (Orentreich et al., 1993; Richie et al., 1994). We found that amino acid restriction increased lysosome levels and reduced amyloid load (Figure 2L). Restriction of even a single amino acid constituent, such as methionine, significantly increased lysosome levels and decreased Aβ and global amyloid content levels (Figure 2L, M). In addition, our results suggest methionine restriction has a distinct, beneficial effect in the clearance of Aβ and amyloid by enhancing lysosome levels. Caloric restriction, including amino acid restriction has been shown to regulate aging and increase lifespan in model organisms. Our results here imply an additional, important role for amino acids in amyloid formation in LOAD.

### Cholesterol regulates lysosomal clearance of amyloid

Lipids represent another major component in nutrition. Cholesterol, a major lipid in the diet, is a risk factor for late-onset AD (Burns and Duff, 2002; Kivipelto and Solomon, 2006). Cholesterol influences production of Aβ by increasing APP and BACE interaction in lipid rafts and removal of cholesterol decreased Aβ production (Ehehalt et al., 2003; Fassbender et al., 2001). Another study also showed that the enzyme Acyl-coenzyme A-cholesterol acyltransferase, which catalyzes the production of cholesteryl-ester levels regulates Aβ production (Di Paolo and Kim, 2011; Puglielli et al., 2001). Recent studies demonstrate that lysosomal cholesterol can increase protein synthesis via activating mTORC1, a master regulator of cell growth (Castellano et al., 2017). Because of our findings that Aβ peptides can also be cleared in the lysosomal compartment, we hypothesized that altered cholesterol levels would also modulate lysosomal clearance of amyloid, in addition to regulating Aβ production.

After loading cells with exogenous cholesterol, surprisingly, an increase, rather than a decrease, in LysoTracker levels was observed (Figure 3A). While higher amino acid levels suppressed lysosomes, higher levels of cholesterol increased the number of LysoTracker positive vesicles. Comparatively, decreasing cholesterol levels by methyl-β-cyclodextrin (MBC) reduced LysoTracker levels. Given this unexpected result, we used another cellular model to validate these findings. Using cells devoid of NPC1 or cells with silenced NPC1 genes, we observed increased intracellular cholesterol levels and again, a higher number of LysoTracker-positive structures following NPC1 protein depletion (Figure 3B). Based on our previous findings, we predicted that the increase in LysoTracker-positive lysosomes would result in more efficient clearance, thus leading to reduced Aβ levels. However, we found higher lysosomal structures accompanied with higher ThioS and intracellular Aβ levels in cholesterol treated cells (Figure 3C, D, I) suggesting that these are most likely defective in clearing amyloid. These results, as such, demonstrate that high cholesterol levels induce more non-functional lysosomes to elicit higher amyloid levels.

**Figure 3:**
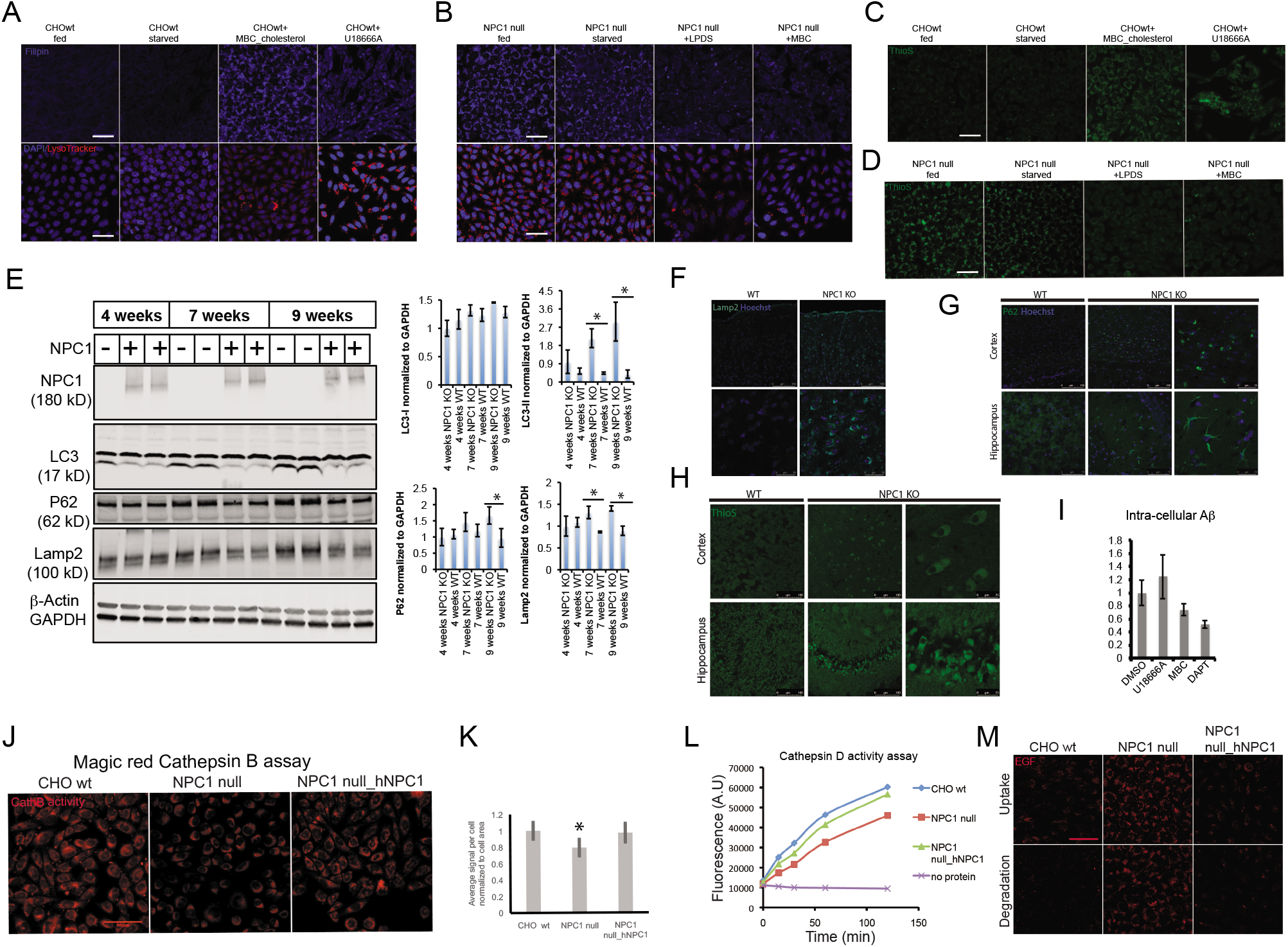
Higher cholesterol increases acidic organelles in cell. However, newly formed lysosomes are non-functional. **A, C.** CHO WT were subjected to overnight serum starvation and loading of extracellular cholesterol by treatment with MBC cholesterol and U18666A. After treatment cells were stained with filipin in (upper panel, blue) (A), lysotracker (lower planel, red) (A), and ThioS (green) (C). DAPI (blue) was used to stain the nucleus in figure A low planel. **B, D.** NPC1 ko cells were subjected to overnight serum starvation and removing of intracellular cholesterol by treatment with MBC and medium containing lipoprotein deficient serum (LPDS). After treatment cells were stained with filipin in (upper panel, blue) (B), lysotracker (lower panel, red) (B), and ThioS (green) (D). DAPI (blue) was used to stain the nucleus in figure B low planel. Scale bar is 40 μm. **E.** Western blot analysis of P62, LC3-I/LC3-II, Lamp2 in WT and NPC1 KO mice of different ages. GAPDH is used as control. Graph shows quantification of LC3-I, LC3-II P62 and P62 signals normalized to GAPDH. **F, G, H.** Lamp2 (green) staining of WT and NPC1 KO mice cortex (F), P62 staining (green) of WT and NPC1 KO mice cortex and hippocampus (G) and ThioS staining of WT and NPC1 KO mice cortex and hippocampus (H). **I.** Intracellular Aβ level was measured after loading or removal of cholesterol on HeLa swAPP cells using electrochemiluminescence assay. **J.** Cathepsin B activity was measured using cathepsin B magic red kit. Scale bar is 50 μm. **K.** Quantification of Fig J. **L.** Cathepsin D enzymatic assay was performed with cell lysates from WT cells, NPC1 knockout cells and NPC1 knockout cells stably expressed human NPC1 protein. **M.** Assessment of EGF uptake and degradation was performed for WT, NPC1 knockout cells and NPC1 knockout cells stably expressed human NPC1 protein. Scale bar is 40 μm.

Previously, we showed that inhibition of the IIN signaling pathway decreases ThioS positive amyloid by increasing lysosome abundance. Upon inhibition of IIN signaling in wild-type cells, we found an increase in LysoTracker positive structures and reduced ThioS positive puncta (SFigure 17A, B). However, in NPC1 null cells, IIN inhibition not only increased LysoTracker levels but also increased ThioS puncta indicating that these cells failed to clear the protein aggregates despite increased lysosome biogenesis. Expression of human NPC1 in the NPC1 null cells rescued this effect. As positive controls, we examined cholesterol levels and observed increased accumulation of cholesterol in NPC1 null cells that positively correlated with higher amyloid levels (SFigure 17C). This suggests that increased LysoTracker staining in the NPC1 null cells state reflected an increase in the abundance of non-functional lysosomes.

To determine the cause for increased non-functional lysosomes in the presence of higher intracellular cholesterol; we speculated that high intracellular cholesterol could impair lysosome function. Recent studies demonstrated that higher cholesterol levels can affect the fusion of autophagosome/late endosome with lysosome (Fader et al., 2009; Fraldi et al., 2010; Furuta et al., 2010). Since this fusion is crucial for trafficking of proteolytic enzymes, such as Cathepsins to lysosomes and are required for lysosomal proteolytic activity (Maruzs et al., 2015), we hypothesized that high cholesterol compromised lysosomal function by decreasing the sorting of lysosomal active enzymes. We thus tested whether high cholesterol would affect autophagosome-lysosome degradation (SFigure 18A) in NPC1 null cells, as demonstrated by accumulation of LC3-II. Indeed, we observed accumulation of LC3-II as well as defective degradation of p62, the cargo of autophagosomal-lysosomal degradation (SFigure 18A). These findings were then validated *in vivo* using mice have spontaneous mutation, which prevents NPC1 production (these mice hereafter termed as NPC1 KO mice). For more details, see at the supplementary method. Similar to cellular findings, we observed a significant age dependent accumulation of LC3-II and p62 in the NPC1 KO mice brain, whereas wild-type brains did not show any gross differences in p62 and LC3-II levels (Figure 3E). Results which indicate impaired autophagosome fusion with lysosomes in NPC1 KO mice. Consistent with these results, we observed significantly higher LAMP2 levels in NPC KO mice compared to wild-type mice, indicating higher lysosomal levels in NPC1 KO mice (Figure 3E). These findings were further validated by immunohistochemistry analysis of LAMP2 and p62 levels. We found significantly increased in LAMP2 and P62 in the cortex of NPC1 KO mice compared to WT control (Figure 3F, G).

As we observed impaired autophagy in NPC1 KO mice consistent with our *in vitro* findings, along with increased lysosomal markers, we decided to analyze amyloid level in NPC1 KO mice. We saw increased ThioS positive signal in the cortex and hippocampus of NPC1 KO mice compared to WT mice (Figure 3H), findings consistent with our *in vitro* studies (Figure 3A-D) thus suggesting defective autophagosomal and lysosomal clearance in NPC1 KO mice. Since we observed significant impaired autophagosome fusion with lysosomes, we examined whether high intracellular cholesterol in NPC1 knockdown cells resulted in defective cathepsin maturation and sorting to lysosomes. We found inadequate sorting of Cathepsin D to lysosome, as confirmed by higher pro-Cathepsin D levels (SFigure 18B) and lower Cathepsin B and Cathepsin D activity in NPC1 null cells (Figure 3J-L). In agreement with defective lysosomal proteolytic function, these cells exhibited increased total amyloid load and decreased degradation of the endosomal cargo, EGF despite of more uptake (Figure 3M). These results demonstrate that high intracellular cholesterol levels impair lysosomal proteolysis, which promotes amyloid formation.

### Nutrient limitation reduces Aβ and amyloid levels in a cellular model

Since over-nutrition and high insulin levels are linked to neurodegenerative diseases (Crane et al., 2013; Fewlass et al., 2004) and caloric restriction may be protective (Longo and Mattson, 2014; Mattson, 2003; Tajes et al., 2010), we examined how caloric restriction affects lysosome biogenesis and amyloid formation. We simulated fasting caloric restriction regimens in cell culture through adjusting serum and nutrient concentrations and compared against standard nutrition conditions, parameters which contrast cell culture protocols which provide constant insulin and nutrients. We found that cells cultured under periodic or chronic fasting conditions displayed increased lysosome biogenesis and reduced amyloid formation relative to those cultured using over-nutrition conditions (Figure 4A). Consistent with these data in cellular conditions, WT mice subjected to a 7 days periodic fasting regimen displayed reduced Aβ levels in cortex (Figure 4B). These results highlight the importance of nutrient limitation to reduce amyloid levels.

### In aged AD animal models, time restricted fasting leads to reduced Aβ and amyloid levels, but also to synapse loss

Through a series of experiments, we showed that insulin/amino acid signaling promotes amyloid formation by transcriptionally suppressing lysosome biogenesis and that high intracellular cholesterol levels suppress lysosomal clearance of amyloid by increasing the number of non-functional lysosomes. In addition, nutrient limitation enhanced amyloid clearance in our cellular model. Therefore, we then tested the effect of fasting *in vivo* using APP transgenic mouse model. Time restricted feeding in 12 months old APP transgenic mice led to increased lysosomal markers, indicating enhanced lysosomal levels (Figure 4F). Similarly, to our *in vitro* results, we observed that fasting led to reductions in amyloid levels (Figure 4C-E). However, we found a surprising effect upon fasting regimens as time restricted feeding also led to the reduction of synaptic markers in cortex and dendritic spine in motor/somatosensory cortex (Figure 4G-I).

### Nutrient limitation enhances phagocytic activity of microglia

Heretofore, we demonstrated that nutrient limitation reduces amyloid load, but may exacerbate synaptic loss, which would be an undesired outcome for AD management. Moreover, it challenges the amyloid-centric model of AD. We previously showed that increased lysosomal levels after depletion of TDP-43 protein in microglia enhanced amyloid clearance and increased synaptic pruning in an AD mouse (Paolicelli et al., 2017). Microglia, being the main phagocytic cells in the brain, play a crucial role in Aβ clearance.

To investigate how IIN-signaling could affect amyloid clearance and synapse loss through microglia and to complement our in vivo observations, we used the BV2 mouse microglia cell line and treated the cells with the AKT-pathway inhibitor or Torin-1, a general mTOR inhibitor, whilst also serum starving the cells. After treatment BV2 cells were incubated with conditioned medium from HeLa swAPP cells containing human Aβ for 12h and an Aβ uptake assay was performed as described in SFigure 19 by measuring the residual Aβ in the supernatant. Upon inhibition of IIN and under starvation conditions, we observed a marked increase in Aβ uptake (Figure 4J) measured through both the ECL and fluorescent Aβ uptake imaging assays (Figure 4K).

We then tested whether inhibition of IIN signaling pathway would alter synaptosome uptake in the cells. To this end, BV2 microglia cells were treated with the Torin-1 or were serum starved, followed by incubation with medium containing synaptosomes isolated from CamKIIcre Tg/+;Rosa26-floxedStopTdTomato mice. Interestingly, we observed increased synaptosome uptake upon inhibition of insulin signaling pathway and under starvation conditions (Figure 4L), demonstrating that inhibition of the IIN signaling pathway not only increases Aβ uptake but also increases synaptosomes uptake in microglia. These in vitro data complement the data from our in vivo experiments with nutrient deprivation in older animals.

### Age dependent effect of insulin and nutrient signaling on AD patients

Here we describe a positive role for IIN signaling in the regulation of amyloid formation, thereby increasing the risk for developing AD. Although a paradoxical role for insulin is observed in late-stage AD patients (Cohen and Dillin, 2008; Steculorum et al., 2014), our results suggest that insulin-induced impaired clearance of amyloid can contribute to synaptic dysfunction in the early stages of late-onset AD. In the late-stages in aging, insulin resistance could contribute to higher catabolic activity in both neurons and microglia that produce synaptic/neuronal atrophy, which further contributes to cognitive impairment (Craft and Watson, 2004; Steculorum et al., 2014). Diabetes, hyperinsulinemia and obesity are linked to a higher incidence of AD (Candeias et al., 2012; Duarte et al., 2012; Moreira, 2012), but it is unresolved whether mid-life or late-life symptoms most contribute to AD development. The first symptoms of late-onset AD manifest after almost 30 years. Excessive insulin signaling during these early stages could promote increased amyloid levels by impairing intracellular clearance to induce synaptic dysfunction (Jack et al., 2010; Villemagne et al., 2013). In the later stages, higher levels of insulin/nutrients could actually slow synaptic deterioration and promote a protective role in synapse strength.

To test whether the hypothesis, that nutrient levels could affect microglia mediated synapse loss in human with AD clinical pathology, we examined cohorts with hyperlipidemia for this study as cohorts with dysregulated amino acid levels were unavailable. Type 2 diabetes (T2D), which was previously linked to increased AD onset risk, was used as a control. Hyperlipidemia is a condition with higher than normal levels of blood cholesterol and triglycerides, and thus is a surrogate marker for higher levels of lipids in the body. Firstly, we observed higher synaptic particles ingested inside microglia from AD patients compared to age-matched control in post-mortem superior temporal lobe, as confirmed by higher co-localization of CD68 and Synapsin1 (Figure 5A, B). We further sub-categorized the cohort based on patients with hyperlipidemia and T2D. We observed reduced internal synaptic particles in microglia in hyperlipidemia conditions compared to those with hypolipidemia, which indicates hyperlipidemia has a protective effect against microglia mediated synapse loss (Figure 5A-C, SFigure 20A, B). To test whether T2D and hyperlipidemia have age dependent effect on AD progression, we compared disease progression in AD patients with T2D, hyperlipidemia and a BMI >25 to AD patients without T2D, hyperlipidemia or a BMI <25, respectively. To account for developing these conditions in mid-versus late-life, we divided the AD cohort into three age groups based on the onset of cognitive symptoms: before 65 years, between 65 to 75 years and after 75 years. The presence of either T2D or hyperlipidemia significantly increased disease progression measured by yearly decline in mini mental state exam (MMSE) scores in the patients whose cognitive symptoms began before 65 years of age. Similarly, patients who had an onset of cognitive symptoms before 65 years and had a BMI >25 showed a trend (p<0.1) towards faster cognitive deterioration. These results are consistent with previous reports on how mid-life diabetes, obesity and hyperlipidemia contribute to higher risk for AD and dementia (Kivipelto et al., 2005; Meyer et al., 2000; Profenno et al., 2010; Xu et al., 2010). These effects were reversed, however, in hyperlipidemic conditions in the older AD patients. Indeed, hyperlipidemia was significantly associated with a slower decline of MMSE in AD patients with an onset of cognitive symptoms after 75 years of age. In T2D patients, or with a BMI>25, older than aged 65, we observed non-significant associations with progression of cognitive impairment (Figure 5D). Together, these results indicate that although higher insulin and nutrient signaling increases the risk of AD in humans by increasing amyloid load, it could be protective for the age group >75 by mitigating microglia mediated synapse loss.

**Figure 4.**
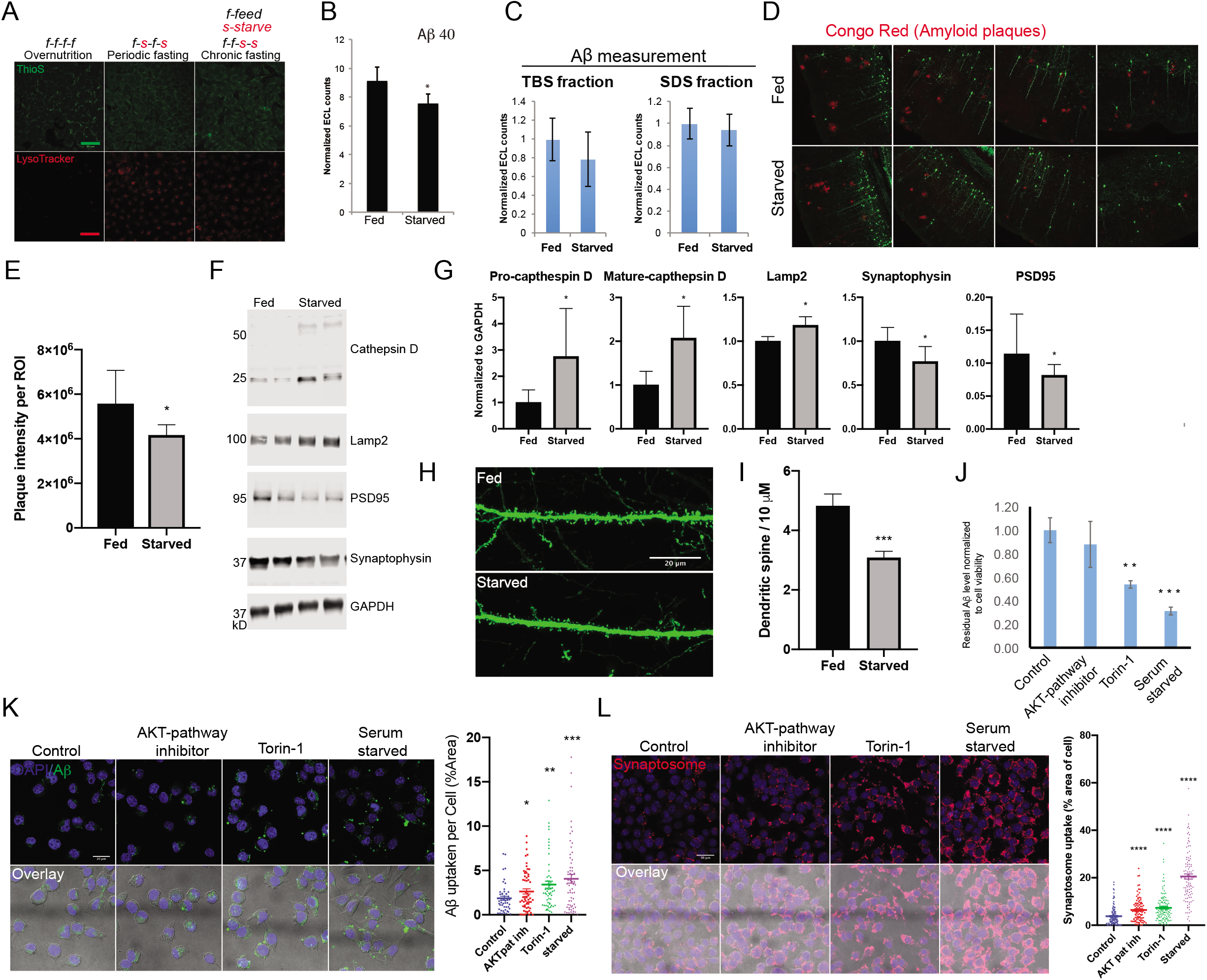
Inhibition of Insulin/amino acid signalling pathway or starvation decrease amyloid level and increases synaptosomes uptake in BV2 microglia cells. **A.** Cells subjected to different starvation paradigms. For overnutrition diet medium was replaced with fresh full medium every 24 h. For periodic fasting medium was replaced with fresh medium, alternating between full medium and serum free medium every 24 h. For chronic fasting medium was replaced with fresh medium, alternating between between full medium and serum free medium every 48 h for four days. After treatment cells were stained with lysotracker (red) and ThioS (green); Scale bar is 50 μm. **B.** Wild type mice were subjected to periodic fasting for 7 days, and their brain homogenates were assayed for Aβ levels. Error bars indicate S.D. *p<0.05. **C.** 12 months old APP Arc, Sw mice were subjected to periodic starvation for a period of 40 days. Following starvation, Aβ was measured from both TBS and SDS fraction of cortex-homogenates. Error bar is SD. **D.** Imaging of brain sections for APP transgenic mice either fed or starved, amyloid plaques (red) stained using Congo Red. **E.** Quantification of (D), **F.** Westernblot analysis showing the effects feeding and starved conditions have on APP transgenic mice lysosomal markers. Blot was stained using lysosomal makers cathepsin D and LAMP2, whilst synaptic markers PSD95 and synaptophysin were also used. GAPDH was used as a loading control. **G.** Quantification of the effects fed/starved conditions have on lysosomal and synaptic markers in APP transgenic mice. **H.** Confocal imaging of cortical dendritic spines from APP transgenic mice from either fed or starved conditions. Green = Thy1 GFP neurons. **I.** Quantification of (H), effects fed/starved conditions have on spine density. Error bar is SD. **J.** Residual Aβ levels upon treatment with AKT-pathway inhibitor, Torin-1 and serum starvation, Error bars indicate S.D. **p<0.005, ***p<0.0005. **K.** Alexa-647 Aβ uptake in BV2 microglia upon treatment with AKT-pathway inhibitor, Torin-1 and serum starvation. Polt is quantification of Fig. 4K. Error bars indicate S.E.M *p<0.05, **p<0.005, ***p<0.0005. **L.** Synaptosomes (red) uptake in BV2 microglia upon treatment with AKT-pathway inhibitor, Torin-1 and serum starvation. DAPI (blue). Plot is quantification of the Fig. 6L.

## Discussion

In this study, we reveal a bi-partite cellular quality control system regulated by the insulin-nutrient signaling that regulates Aβ peptide clearance through the lysosomal pathway in neurons while also regulating microglia-dependent synapse loss. These findings are important as both processes are causally associated with AD. We also demonstrate that neuronal lysosomes can clear Aβ peptides, which is regulated by IIN signaling. This previously uncharacterized clearance mechanism provides additional neuronal Aβ clearance through non-cell autonomous mechanisms via microglia (Chung et al., 1999), interstitial fluid to CSF bulk flow, efflux through blood brain barrier (Bell et al., 2009; Shibata et al., 2000) and enzymatic degradation (Iwata et al., 2000; Qiu et al., 1998). Exploiting this new intracellular clearance mechanism will provide an additional therapeutic target in AD, particularly for late-onset AD patients who do not express mutations linked to APP processing and Aβ production.

Unlike familial AD, hyperinsulinemia and diabetes are additional risk factors for late onset AD (Candeias et al., 2012; Duarte et al., 2012; Moreira, 2012), which may develop due to associated impairments with Aβ clearance (Candeias et al., 2012; Duarte et al., 2012; Moreira, 2012). We also demonstrate that high insulin/nutrient levels, in particular amino acid levels, activate AKT signaling, which inhibits lysosome biogenesis and lysosomal clearance of Aβ and amyloid proteins (summary in SFigure 21). This reduced intracellular clearance of Aβ facilitates an increase in amyloid load and risk for developing AD.

In addition to insulin and amino acids, we also demonstrate a possible mechanism through which higher cellular cholesterol alters lysosomal clearance of amyloid. While amino acids influence lysosomal clearance through lysosome biogenesis involving transcription, high cholesterol levels impair the proteolytic activity of lysosomes. Excessive cholesterol accumulation in late-endosomes and lysosomes can impair SNARE function (Fraldi et al., 2010). SNARE function is critical to fuse autophagosomes to lysosomes (Fader et al., 2009; Furuta et al., 2010). This impairment demonstrates why higher cholesterol levels are a risk factor for impaired autophagy. Autophagosomes clear cytosolic protein aggregates for lysosome degradation (Barbero-Camps et al., 2018). Impaired autophagosome-lysosome fusion will promote the accumulation of cytosolic protein aggregates, similar to previous studies that established a link between impaired autophagy and Parkinson’s disease (PD) (Cuervo et al., 2004). Our study, then, provides a possible explanation for why high dietary cholesterol intake increases the risk for PD (de Lau et al., 2006; Hu et al., 2008; Powers et al., 2009).

Several epidemiological studies suggest high cholesterol levels are a risk factor for AD (Burns and Duff, 2002; Kivipelto and Solomon, 2006). A high cholesterol environment alters APP processing (Burns et al., 2003; Runz et al., 2002; Simons et al., 1998; Wahrle et al., 2002), leading to increased production of the Aβ peptide (Simons et al., 1998). While cholesterol increases the production of the Aβ peptide, we also show that cholesterol compromises Aβ clearance through lysosomal dysfunction, a result supported by evidence that cholesterol-reducing agents (statins) can protect against AD (Fassbender et al., 2001). A recent report lends further validation to our study showing that high brain cholesterol enhanced autophagosome formation but disrupted its fusion with endosomal-lysosomal vesicles(Barbero-Camps et al., 2018). Furthermore, our results have important implications for nutrient and brain cholesterol levels and the risk of elevated production of amyloid and developing Alzheimer’s disease. Our results, then, indicate that lifestyle changes are essential for healthy aging and reducing disease risk factors.

Our results also further suggest that excessive insulin signaling from over-nutrition could contribute to the development of late-onset AD. Overlaying global maps for per capita food intake with AD incidence is strikingly correlated. AD correlates with obesity, diabetes and elevated blood sugar (Crane et al., 2013; Ott et al., 1999). If excessive IIN signaling does contribute to late-onset AD initiation, then lifestyle changes with reduced food intake, caloric restriction or periodic fasting could be preventative (Brandhorst et al., 2015; Longo and Mattson, 2014; Mattson, 2003, 2004; Mattson et al., 2014). For example, intermittent fasting and energy consumption through exercise could reduce INS signaling and protect against AD. Although a paradoxical role for insulin signaling exists in late-stage AD patients (Cohen and Dillin, 2008; Steculorum et al., 2014), we postulate that insulin-induced impairments in amyloid clearance contribute to amyloid accumulation in early, preclinical stages of late-onset AD. Once amyloid oligomers form, our results imply they could bind to insulin receptors to confer insulin resistance at later stages (De Felice, 2013). An alternative explanation would be that our bodies enter a catabolism-dominant phase during the later stages of aging, due to anorexia and frailty, wherein a positive IIN signaling could contribute to less atrophy due to inhibition of autophagy and lysosomal proteolysis.

An interesting, unanticipated result from our study is the finding that insulin and nutrient signaling regulates Aβ levels by negatively modulating lysosomal clearance in neurons, but in microglia, the same pathway controls Aβ phagocytosis and synapse loss. Time restricted feeding in older AD mice resulted in synapse loss despite an increased in Aβ clearance. This is similar to our previously published observations in microglial TDP-43 depleted mice, which demonstrated enhanced synapse loss despite effective amyloid clearance (Paolicelli et al., 2017). Because of these results, a key question arises: Is fasting or caloric restriction protective against AD even if it were to enhance synapse loss in the older patients? If so, is there a time window when fasting might be beneficial and a phase where it might not. Our clinical data indicate that T2D and hyperlipidemia increase the risk of dementia in the age group below 70 (Figure 5B-D). However, hyperlipidemia could be protective above 75 years of age against cognitive loss as higher nutrient levels could suppress microglial phagocytosis of synapses (Jawaid et al., 2018).

**Figure 5.**
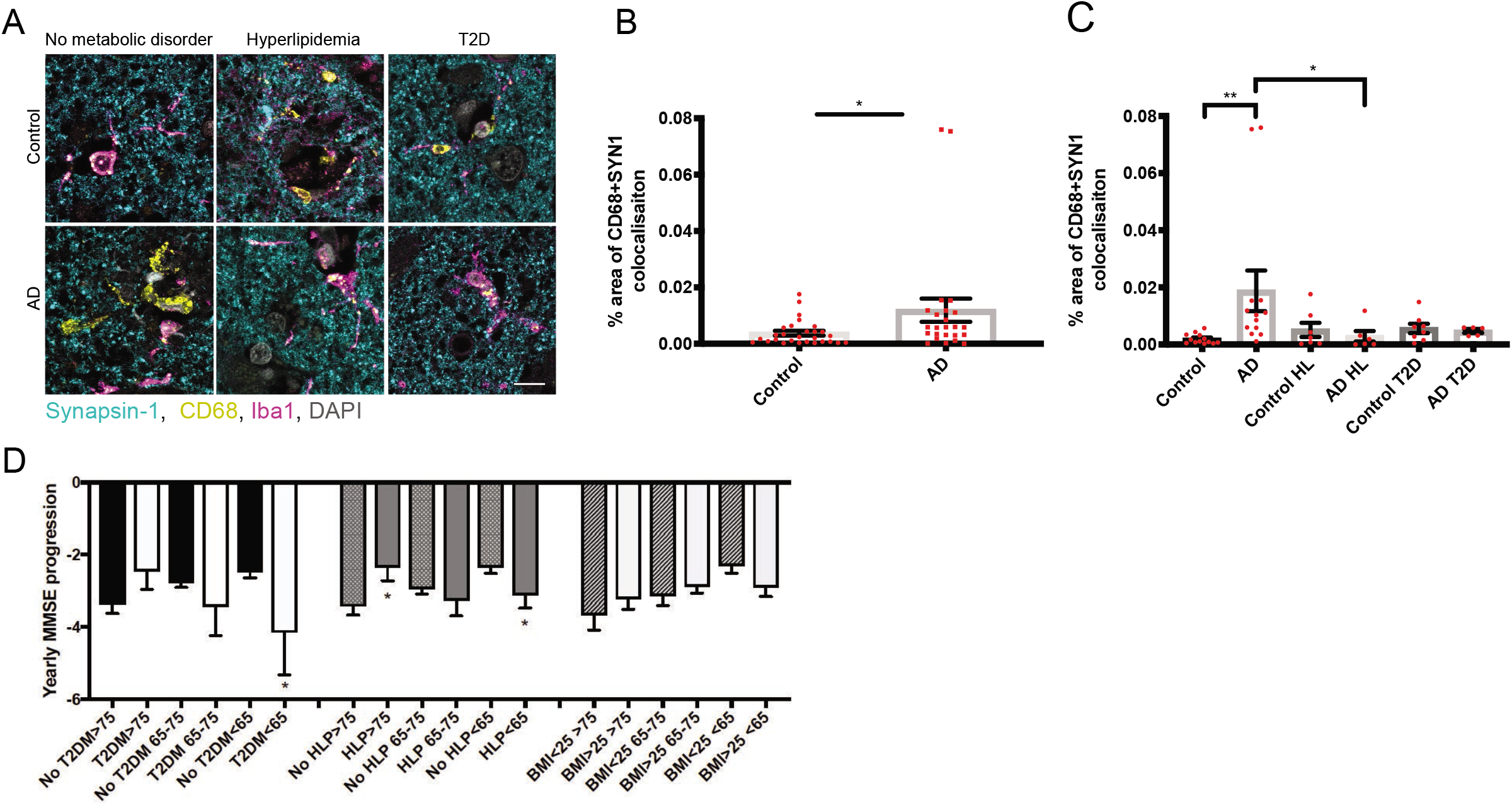
Effects hyperlipidemia, T2D, BMI and age in combinations have on AD pathological status. **A.** Post-mortem tissue from superior temporal lobe of AD and age matched control are stained for microglial markers Iba1 (magenta) and CD68 (yellow), and pre-synaptic marker Synapsin-1 (cyan) and DAPI for nuclei (grey). Representative images are shown. Scale bar 10 microns. **B**. Post-mortem comparison of Synapsin-1 inside CD68-positive cells from age-matched control cases (n=28) and Alzheimer’s disease (AD) (n=24) cases showed increased levels of Synapsin-1 inside CD68 in AD (Mann-Whitney U test, p=0.0274). **C.** Control and AD cases are stratified by lack or presence of metabolic disorders, namely hyperlipidemia (HL) and type 2 diabetes (T2D). AD cases with no metabolic disorder (AD) showed significantly greater Synapsin-1/CD68 colocalisation compared to control with no metabolic disorder (Control) (p=0.022) and AD with hyperlipidaemia (AD HL) (p=0.0196) (Kruskal-Wallis test). AD cases with T2D showed no difference to controls with T2D. **D.** Disease progression analysis of AD patients with T2D, hyperlipidemia and a BMI>25 against AD patients without T2DM, hyperlipidemia or a BMI<25. Measured using yearly progression scores taken from MMSE.

It is tempting to postulate a biphasic role for insulin and nutrient signaling during the development of AD pathology-the first controlling amyloid clearance and the later controlling synaptic loss, which contribute to dementia. In mid-life, where anabolic and catabolic processes needed to be kept in equilibrium, excess insulin and nutrient signaling on the one-hand could promote Aβ load in neurons by suppressing neuronal lysosomal clearance, and also microglial Aβ uptake and clearance. Together it leads to amyloid accumulation and this increases the risk of dementia. However, at the age above 75, due to anorexia of aging, catabolic activity increases, with enhanced lysosomal enzymes levels and activity (Verdugo and Ray, 1997). Increased lysosomal activity due to suppression of IIN signaling in microglia leads to higher synaptic pruning which could contribute to a progressive deterioration in cognitive function. This indicates that high insulin and increased nutrient signaling above age 75 could suppress microglia mediated synapse loss. From these results, we would hypothesize that two things. First, that amyloid reducing therapies should target the mid-life, amyloid phase prior to when synapse loss occurs and synapse stabilizing therapies should be developed to target the synaptic phase. Second, that depending on the metabolic state of the individual, fasting or caloric restriction in the midlife phase and nutrient-rich intake in the old age phase to offset the catabolic dominance in this phase would be protective against AD.

## Supporting information

Human cases details

## Abbreviations

Aβ: Amyloid-β;
AD: Alzheimer’s Disease;
APP: amyloid precursor protein;
BACE1: β-amyloid cleaving enzyme 1;
RNAi: RNA interference;
PEN2: Presenilin enhancer 2

## Author contributions

LR, MM designed research, MM, JB, RP, MT, TS-J, VU, AJ, KCV, SBH, EM, RS, GT, MD, CP, MM, KD, MS, JK, OB, PK performed research, MM, JB, RP, VU, AJ, KCV, SBH, EM, RS, GT, MD, CP, MM, KD, SH, MS, JK, OB, SF, PK, RN, CH, PS, RG and LR analyzed the data. MT performed the immunohistochemistry analysis of human brain tissue for synapse engulfment assay. M.S provided reagents. LR, MM and JB wrote the paper and all the authors participated in the editing of the paper.

## Acknowledgements

We thank G. Yu for the HeLa-sweAPP cells, M. Neumann and the ADDF (Alzheimer Drug Discovery Foundation) for the help with Kinase siRNA library. We thank Dr. S. Ferguson for the generous support with TFEB-GFP constructs and cell lines. We thank Dr. Tara Spires-Jones for kindly providing immortalized astrocytes expressing different isoforms of ApoE. We thank Dr. D. Ory for providing us NPC1 null cells and NPC1 null cells stably expressed human NPC1 cell-line. We thank A. Aguzzi, W. Annaert, R. Sawarkar and K. Simons for critical input in the study and Z. Goodger, S. Hoey and W. Zacchariae for the critical reading of the manuscript. We thank K. Korn, M. Stoeter and M. Bickle for the help and advice with the screen and V. Surendranath for the help with bioinformatics analysis on the screen hits. We thank J. Oorschot for the help with electron microscopy. We thank Andrea Valeri for the help with animal experiement. We thank S. Hoey, B. Siegenthaler, G. Siegel, J. Issa, B. Felmy, R. Bolognini, G. Minakaki and F. Kivrak-Pfiffner for the technical help. We thank Ivo Carre, Shiden Solomon, Nirmal Kumar Sampathkumar and Jacqueline C Mitchell for help with editing the manuscript. We thank Life Science Editors for editing assistance. L.R acknowledges the Professorship Grant and financial support from the Velux Foundation. Swiss National Science Foundation grant, the Novartis Foundation grant, Bangerter Stiftung, Baugarten Stiftung and the Synapsis Foundation are acknowledged. MM acknowledge Forschungskredit Candoc, University of Zürich for funding. LR and VU acknowledge the funding support from the European Neuroscience Campus of the Erasmus Mundus Program. L.R. and S.H. acknowledge the funding support from the SCOPES programme of the Swiss National Science Foundation. M.S. acknowledges NSF funding from DGE-1143954. S.M.F. acknowledges funding support from the NIH (AG062210 and GM105718). Some samples and pathological diagnoses were provided by the MRC London Neurodegenerative Diseases Brain Bank and the Manchester Brain Bank from Brains for Dementia Research, jointly funded from ARUK and AS via ABBUK Ltd.

This work was funded by the UK Dementia Research Institute which receives its funding from DRI Ltd, funded by the UK Medical Research Council, Alzheimer’s Society, and Alzheimer’s Research UK, and the European Research Council (ERC) under the European Union’s Horizon 2020 research and innovation programme (Grant agreement No. 681181). We would like to thank the South West Dementia Brain Bank (SWDBB) for providing brain tissue for this study. The SWDBB is part of the Brains for Dementia Research programme, jointly funded by Alzheimer’s Research UK and Alzheimer’s Society and is supported by BRACE (Bristol Research into Alzheimer’s and Care of the Elderly) and the Medical Research Council. Some of the control participants in the human study were from the Lothian Birth Cohort 1936, thus we wish to thank the cohort and research team supported by Age UK (Disconnected Mind project) in The University of Edinburgh Centre for Cognitive Ageing and Cognitive Epidemiology, funded by the Biotechnology and Biological Sciences Research Council (BBSRC) and Medical Research Council (MRC) ((MR/K026992/1).

## References

Andersen, O.M., Reiche, J., Schmidt, V., Gotthardt, M., Spoelgen, R., Behlke, J., von Arnim, C.A., Breiderhoff, T., Jansen, P., Wu, X., et al. (2005). Neuronal sorting protein-related receptor sorLA/LR11 regulates processing of the amyloid precursor protein. Proc Natl Acad Sci U S A 102, 13461–13466.

Bali, J., Gheinani, A.H., Zurbriggen, S., and Rajendran, L. (2012). Role of genes linked to sporadic Alzheimer’s disease risk in the production of beta-amyloid peptides. Proc Natl Acad Sci U S A 109, 15307–15311.

Barbero-Camps, E., Roca-Agujetas, V., Bartolessis, I., de Dios, C., Fernandez-Checa, J.C., Mari, M., Morales, A., Hartmann, T., and Colell, A. (2018). Cholesterol impairs autophagy-mediated clearance of amyloid beta while promoting its secretion. Autophagy 14, 1129–1154.

Bell, R.D., Deane, R., Chow, N., Long, X., Sagare, A., Singh, I., Streb, J.W., Guo, H., Rubio, A., Van Nostrand, W., et al. (2009). SRF and myocardin regulate LRP-mediated amyloid-beta clearance in brain vascular cells. Nat Cell Biol 11, 143–153.

Ben Halima, S., Mishra, S., Raja, K.M., Willem, M., Baici, A., Simons, K., Brustle, O., Koch, P., Haass, C., Caflisch, A., et al. (2016). Specific Inhibition of beta-Secretase Processing of the Alzheimer Disease Amyloid Precursor Protein. Cell Rep 14, 2127–2141.

Ben Halima, S., and Rajendran, L. (2011). Membrane anchored and lipid raft targeted beta-secretase inhibitors for Alzheimer’s disease therapy. J Alzheimers Dis 24 Suppl 2, 143–152.

Blommaart, E.F., Luiken, J.J., Blommaart, P.J., van Woerkom, G.M., and Meijer, A.J. (1995a). Phosphorylation of ribosomal protein S6 is inhibitory for autophagy in isolated rat hepatocytes. J Biol Chem 270, 2320–2326.

Blommaart, P.J., Charles, R., Meijer, A.J., and Lamers, W.H. (1995b). Changes in hepatic nitrogen balance in plasma concentrations of amino acids and hormones and in cell volume after overnight fasting in perinatal and adult rat. Pediatr Res 38, 1018–1025.

Borchelt, D.R., Thinakaran, G., Eckman, C.B., Lee, M.K., Davenport, F., Ratovitsky, T., Prada, C.M., Kim, G., Seekins, S., Yager, D., et al. (1996). Familial Alzheimer’s disease-linked presenilin 1 variants elevate Abeta1-42/1-40 ratio in vitro and in vivo. Neuron 17, 1005–1013.

Brandhorst, S., Choi, I.Y., Wei, M., Cheng, C.W., Sedrakyan, S., Navarrete, G., Dubeau, L., Yap, L.P., Park, R., Vinciguerra, M., et al. (2015). A Periodic Diet that Mimics Fasting Promotes Multi-System Regeneration, Enhanced Cognitive Performance, and Healthspan. Cell Metab 22, 86–99.

Buggia-Prevot, V., Fernandez, C.G., Udayar, V., Vetrivel, K.S., Elie, A., Roseman, J., Sasse, V.A., Lefkow, M., Meckler, X., Bhattacharyya, S., et al. (2013). A function for EHD family proteins in unidirectional retrograde dendritic transport of BACE1 and Alzheimer’s disease Abeta production. Cell Rep 5, 1552–1563.

Burns, M., and Duff, K. (2002). Cholesterol in Alzheimer’s disease and tauopathy. Ann N Y Acad Sci 977, 367–375.

Burns, M., Gaynor, K., Olm, V., Mercken, M., LaFrancois, J., Wang, L., Mathews, P.M., Noble, W., Matsuoka, Y., and Duff, K. (2003). Presenilin redistribution associated with aberrant cholesterol transport enhances beta-amyloid production in vivo. J Neurosci 23, 5645–5649.

Candeias, E., Duarte, A.I., Carvalho, C., Correia, S.C., Cardoso, S., Santos, R.X., Placido, A.I., Perry, G., and Moreira, P.I. (2012). The impairment of insulin signaling in Alzheimer’s disease. IUBMB life 64, 951–957.

Castellano, B.M., Thelen, A.M., Moldavski, O., Feltes, M., van der Welle, R.E., Mydock-McGrane, L., Jiang, X., van Eijkeren, R.J., Davis, O.B., Louie, S.M., et al. (2017). Lysosomal cholesterol activates mTORC1 via an SLC38A9-Niemann-Pick C1 signaling complex. Science 355, 1306–1311.

Chiti, F., Stefani, M., Taddei, N., Ramponi, G., and Dobson, C.M. (2003). Rationalization of the effects of mutations on peptide and protein aggregation rates. Nature 424, 805-808.

Chung, H., Brazil, M.I., Soe, T.T., and Maxfield, F.R. (1999). Uptake, degradation, and release of fibrillar and soluble forms of Alzheimer’s amyloid beta-peptide by microglial cells. J Biol Chem 274, 32301–32308.

Cohen, E., and Dillin, A. (2008). The insulin paradox: aging, proteotoxicity and neurodegeneration. Nat Rev Neurosci 9, 759–767.

Craft, S., and Watson, G.S. (2004). Insulin and neurodegenerative disease: shared and specific mechanisms. Lancet Neurol 3, 169–178.

Crane, P.K., Walker, R., Hubbard, R.A., Li, G., Nathan, D.M., Zheng, H., Haneuse, S., Craft, S., Montine, T.J., Kahn, S.E., et al. (2013). Glucose levels and risk of dementia. N Engl J Med 369, 540–548.

Cuervo, A.M., Stefanis, L., Fredenburg, R., Lansbury, P.T., and Sulzer, D. (2004). Impaired degradation of mutant alpha-synuclein by chaperone-mediated autophagy. Science 305, 1292–1295.

De Felice, F.G. (2013). Alzheimer’s disease and insulin resistance: translating basic science into clinical applications. J Clin Invest 123, 531–539.

de Lau, L.M., Koudstaal, P.J., Hofman, A., and Breteler, M.M. (2006). Serum cholesterol levels and the risk of Parkinson’s disease. Am J Epidemiol 164, 998–1002.

De Strooper, B. (2010). Proteases and proteolysis in Alzheimer disease: a multifactorial view on the disease process. Physiol Rev 90, 465–494.

Demetriades, C., Doumpas, N., and Teleman, A.A. (2014). Regulation of TORC1 in response to amino acid starvation via lysosomal recruitment of TSC2. Cell 156, 786-799.

Di Paolo, G., and Kim, T.W. (2011). Linking lipids to Alzheimer’s disease: cholesterol and beyond. Nat Rev Neurosci 12, 284–296.

Duarte, A.I., Moreira, P.I., and Oliveira, C.R. (2012). Insulin in central nervous system: more than just a peripheral hormone. Journal of aging research 2012, 384017.

Edgar, J.R., Willen, K., Gouras, G.K., and Futter, C.E. (2015). ESCRTs regulate amyloid precursor protein sorting in multivesicular bodies and intracellular amyloid-beta accumulation. J Cell Sci 128, 2520–2528.

Ehehalt, R., Keller, P., Haass, C., Thiele, C., and Simons, K. (2003). Amyloidogenic processing of the Alzheimer beta-amyloid precursor protein depends on lipid rafts. J Cell Biol 160, 113–123.

Fader, C.M., Sanchez, D.G., Mestre, M.B., and Colombo, M.I. (2009). TI-VAMP/VAMP7 and VAMP3/cellubrevin: two v-SNARE proteins involved in specific steps of the autophagy/multivesicular body pathways. Biochim Biophys Acta 1793, 1901–1916.

Fassbender, K., Simons, M., Bergmann, C., Stroick, M., Lutjohann, D., Keller, P., Runz, H., Kuhl, S., Bertsch, T., von Bergmann, K., et al. (2001). Simvastatin strongly reduces levels of Alzheimer’s disease beta-amyloid peptides Abeta 42 and Abeta 40 in vitro and in vivo. Proc Natl Acad Sci U S A 98, 5856–5861.

Fewlass, D.C., Noboa, K., Pi-Sunyer, F.X., Johnston, J.M., Yan, S.D., and Tezapsidis, N. (2004). Obesity-related leptin regulates Alzheimer’s Abeta. FASEB J 18, 1870–1878.

Fraldi, A., Annunziata, F., Lombardi, A., Kaiser, H.J., Medina, D.L., Spampanato, C., Fedele, A.O., Polishchuk, R., Sorrentino, N.C., Simons, K., et al. (2010). Lysosomal fusion and SNARE function are impaired by cholesterol accumulation in lysosomal storage disorders. EMBO J 29, 3607–3620.

Furuta, N., Fujita, N., Noda, T., Yoshimori, T., and Amano, A. (2010). Combinational soluble N-ethylmaleimide-sensitive factor attachment protein receptor proteins VAMP8 and Vti1b mediate fusion of antimicrobial and canonical autophagosomes with lysosomes. Mol Biol Cell 21, 1001–1010.

Glabe, C. (2001). Intracellular mechanisms of amyloid accumulation and pathogenesis in Alzheimer’s disease. J Mol Neurosci 17, 137–145.

Gouras, G.K., Almeida, C.G., and Takahashi, R.H. (2005). Intraneuronal Abeta accumulation and origin of plaques in Alzheimer’s disease. Neurobiol Aging 26, 1235–1244.

Grbovic, O.M., Mathews, P.M., Jiang, Y., Schmidt, S.D., Dinakar, R., Summers-Terio, N.B., Ceresa, B.P., Nixon, R.A., and Cataldo, A.M. (2003). Rab5-stimulated up-regulation of the endocytic pathway increases intracellular beta-cleaved amyloid precursor protein carboxyl-terminal fragment levels and Abeta production. J Biol Chem 278, 31261–31268.

Haass, C., Koo, E.H., Mellon, A., Hung, A.Y., and Selkoe, D.J. (1992). Targeting of cellsurface beta-amyloid precursor protein to lysosomes: alternative processing into amyloid-bearing fragments. Nature 357, 500–503.

Haass, C., and Selkoe, D.J. (2007). Soluble protein oligomers in neurodegeneration: lessons from the Alzheimer’s amyloid beta-peptide. Nat Rev Mol Cell Biol 8, 101–112.

Howell, S., Nalbantoglu, J., and Crine, P. (1995). Neutral endopeptidase can hydrolyze beta-amyloid(1-40) but shows no effect on beta-amyloid precursor protein metabolism. Peptides 16, 647–652.

Hu, G., Antikainen, R., Jousilahti, P., Kivipelto, M., and Tuomilehto, J. (2008). Total cholesterol and the risk of Parkinson disease. Neurology 70, 1972–1979.

Iliff, J.J., Wang, M., Liao, Y., Plogg, B.A., Peng, W., Gundersen, G.A., Benveniste, H., Vates, G.E., Deane, R., Goldman, S.A., et al. (2012). A paravascular pathway facilitates CSF flow through the brain parenchyma and the clearance of interstitial solutes, including amyloid beta. Sci Transl Med 4, 147ra111.

Iwata, N., Tsubuki, S., Takaki, Y., Watanabe, K., Sekiguchi, M., Hosoki, E., Kawashima-Morishima, M., Lee, H.J., Hama, E., Sekine-Aizawa, Y., et al. (2000). Identification of the major Abeta1-42-degrading catabolic pathway in brain parenchyma: suppression leads to biochemical and pathological deposition. Nat Med 6, 143–150.

Jack, C.R., Jr., Knopman, D.S., Jagust, W.J., Shaw, L.M., Aisen, P.S., Weiner, M.W., Petersen, R.C., and Trojanowski, J.Q. (2010). Hypothetical model of dynamic biomarkers of the Alzheimer’s pathological cascade. Lancet Neurol 9, 119–128.

Jawaid, A., Khan, R., Polymenidou, M., and Schulz, P.E. (2018). Disease-modifying effects of metabolic perturbations in ALS/FTLD. Mol Neurodegener 13, 63.

Keilani, S., Lun, Y., Stevens, A.C., Williams, H.N., Sjoberg, E.R., Khanna, R., Valenzano, K.J., Checler, F., Buxbaum, J.D., Yanagisawa, K., et al. (2012). Lysosomal dysfunction in a mouse model of Sandhoff disease leads to accumulation of ganglioside-bound amyloid-beta peptide. J Neurosci 32, 5223–5236.

Kim, J., Kundu, M., Viollet, B., and Guan, K.L. (2011). AMPK and mTOR regulate autophagy through direct phosphorylation of Ulk1. Nat Cell Biol 13, 132–141.

Kinoshita, A., Fukumoto, H., Shah, T., Whelan, C.M., Irizarry, M.C., and Hyman, B.T. (2003). Demonstration by FRET of BACE interaction with the amyloid precursor protein at the cell surface and in early endosomes. J Cell Sci 116, 3339–3346.

Kivipelto, M., Ngandu, T., Fratiglioni, L., Viitanen, M., Kareholt, I., Winblad, B., Helkala, E.L., Tuomilehto, J., Soininen, H., and Nissinen, A. (2005). Obesity and vascular risk factors at midlife and the risk of dementia and Alzheimer disease. Arch Neurol 62, 1556–1560.

Kivipelto, M., and Solomon, A. (2006). Cholesterol as a risk factor for Alzheimer’s disease – epidemiological evidence. Acta Neurol Scand Suppl 185, 50–57.

Koo, E.H., and Squazzo, S.L. (1994). Evidence that production and release of amyloid beta-protein involves the endocytic pathway. J Biol Chem 269, 17386–17389.

Kress, B.T., Iliff, J.J., Xia, M., Wang, M., Wei, H.S., Zeppenfeld, D., Xie, L., Kang, H., Xu, Q., Liew, J.A., et al. (2014). Impairment of paravascular clearance pathways in the aging brain. Ann Neurol 76, 845–861.

Leissring, M.A., Farris, W., Chang, A.Y., Walsh, D.M., Wu, X., Sun, X., Frosch, M.P., and Selkoe, D.J. (2003). Enhanced proteolysis of beta-amyloid in APP transgenic mice prevents plaque formation, secondary pathology, and premature death. Neuron 40, 1087–1093.

Longo, V.D., and Mattson, M.P. (2014). Fasting: molecular mechanisms and clinical applications. Cell Metab 19, 181–192.

Martina, J.A., Diab, H.I., Li, H., and Puertollano, R. (2014). Novel roles for the MiTF/TFE family of transcription factors in organelle biogenesis, nutrient sensing, and energy homeostasis. Cell Mol Life Sci 71, 2483–2497.

Maruzs, T., Lorincz, P., Szatmari, Z., Szeplaki, S., Sandor, Z., Lakatos, Z., Puska, G., Juhasz, G., and Sass, M. (2015). Retromer Ensures the Degradation of Autophagic Cargo by Maintaining Lysosome Function in Drosophila. Traffic 16, 1088–1107.

Mattson, M.P. (2003). Will caloric restriction and folate protect against AD and PD? Neurology 60, 690–695.

Mattson, M.P. (2004). Pathways towards and away from Alzheimer’s disease. Nature 430, 631–639.

Mattson, M.P., Allison, D.B., Fontana, L., Harvie, M., Longo, V.D., Malaisse, W.J., Mosley, M., Notterpek, L., Ravussin, E., Scheer, F.A., et al. (2014). Meal frequency and timing in health and disease. Proc Natl Acad Sci U S A 111, 16647–16653.

Mawuenyega, K.G., Sigurdson, W., Ovod, V., Munsell, L., Kasten, T., Morris, J.C., Yarasheski, K.E., and Bateman, R.J. (2010). Decreased clearance of CNS beta-amyloid in Alzheimer’s disease. Science 330, 1774.

Meisl, G., Yang, X., Hellstrand, E., Frohm, B., Kirkegaard, J.B., Cohen, S.I., Dobson, C.M., Linse, S., and Knowles, T.P. (2014). Differences in nucleation behavior underlie the contrasting aggregation kinetics of the Abeta40 and Abeta42 peptides. Proc Natl Acad Sci U S A 111, 9384–9389.

Menon, S., Dibble, C.C., Talbott, G., Hoxhaj, G., Valvezan, A.J., Takahashi, H., Cantley, L.C., and Manning, B.D. (2014). Spatial control of the TSC complex integrates insulin and nutrient regulation of mTORC1 at the lysosome. Cell 156, 771–785.

Meyer, J.S., Rauch, G.M., Rauch, R.A., Haque, A., and Crawford, K. (2000). Cardiovascular and other risk factors for Alzheimer’s disease and vascular dementia. Ann N Y Acad Sci 903, 411–423.

Moreira, P.I. (2012). Alzheimer’s disease and diabetes: an integrative view of the role of mitochondria, oxidative stress, and insulin. J Alzheimers Dis 30 Suppl 2, S199–215.

Morel, E., Chamoun, Z., Lasiecka, Z.M., Chan, R.B., Williamson, R.L., Vetanovetz, C., Dall’Armi, C., Simoes, S., Point Du Jour, K.S., McCabe, B.D., et al. (2013). Phosphatidylinositol-3-phosphate regulates sorting and processing of amyloid precursor protein through the endosomal system. Nat Commun 4, 2250.

Nixon, R.A., and Cataldo, A.M. (2006). Lysosomal system pathways: genes to neurodegeneration in Alzheimer’s disease. J Alzheimers Dis 9, 277–289.

Nixon, R.A., Cataldo, A.M., Paskevich, P.A., Hamilton, D.J., Wheelock, T.R., and Kanaley-Andrews, L. (1992). The lysosomal system in neurons. Involvement at multiple stages of Alzheimer’s disease pathogenesis. Ann N Y Acad Sci 674, 65–88.

Nixon, R.A., Mathews, P.M., and Cataldo, A.M. (2001). The neuronal endosomal-lysosomal system in Alzheimer’s disease. J Alzheimers Dis 3, 97–107.

Orentreich, N., Matias, J.R., DeFelice, A., and Zimmerman, J.A. (1993). Low methionine ingestion by rats extends life span. The Journal of nutrition 123, 269–274.

Ott, A., Stolk, R.P., van Harskamp, F., Pols, H.A., Hofman, A., and Breteler, M.M. (1999). Diabetes mellitus and the risk of dementia: The Rotterdam Study. Neurology 53, 1937–1942.

Paolicelli, R.C., Jawaid, A., Henstridge, C.M., Valeri, A., Merlini, M., Robinson, J.L., Lee, E.B., Rose, J., Appel, S., Lee, V.M., et al. (2017). TDP-43 Depletion in Microglia Promotes Amyloid Clearance but Also Induces Synapse Loss. Neuron 95, 297–308 e296.

Parr, C., Carzaniga, R., Gentleman, S.M., Van Leuven, F., Walter, J., and Sastre, M. (2012). Glycogen synthase kinase 3 inhibition promotes lysosomal biogenesis and autophagic degradation of the amyloid-beta precursor protein. Mol Cell Biol 32, 4410–4418.

Powers, K.M., Smith-Weller, T., Franklin, G.M., Longstreth, W.T., Jr., Swanson, P.D., and Checkoway, H. (2009). Dietary fats, cholesterol and iron as risk factors for Parkinson’s disease. Parkinsonism Relat Disord 15, 47–52.

Profenno, L.A., Porsteinsson, A.P., and Faraone, S.V. (2010). Meta-analysis of Alzheimer’s disease risk with obesity, diabetes, and related disorders. Biol Psychiatry 67, 505–512.

Puertollano, R., Ferguson, S.M., Brugarolas, J., and Ballabio, A. (2018). The complex relationship between TFEB transcription factor phosphorylation and subcellular localization. EMBO J 37.

Puglielli, L., Konopka, G., Pack-Chung, E., Ingano, L.A., Berezovska, O., Hyman, B.T., Chang, T.Y., Tanzi, R.E., and Kovacs, D.M. (2001). Acyl-coenzyme A: cholesterol acyltransferase modulates the generation of the amyloid beta-peptide. Nat Cell Biol 3, 905–912.

Qiu, W.Q., Walsh, D.M., Ye, Z., Vekrellis, K., Zhang, J., Podlisny, M.B., Rosner, M.R., Safavi, A., Hersh, L.B., and Selkoe, D.J. (1998). Insulin-degrading enzyme regulates extracellular levels of amyloid beta-protein by degradation. J Biol Chem 273, 32730–32738.

Rajendran, L., Honsho, M., Zahn, T.R., Keller, P., Geiger, K.D., Verkade, P., and Simons, K. (2006). Alzheimer’s disease beta-amyloid peptides are released in association with exosomes. Proc Natl Acad Sci U S A 103, 11172–11177.

Rajendran, L., Schneider, A., Schlechtingen, G., Weidlich, S., Ries, J., Braxmeier, T., Schwille, P., Schulz, J.B., Schroeder, C., Simons, M., et al. (2008). Efficient inhibition of the Alzheimer’s disease beta-secretase by membrane targeting. Science 320, 520–523.

Richie, J.P., Jr., Leutzinger, Y., Parthasarathy, S., Malloy, V., Orentreich, N., and Zimmerman, J.A. (1994). Methionine restriction increases blood glutathione and longevity in F344 rats. Faseb J 8, 1302–1307.

Roberts, B.R., Lind, M., Wagen, A.Z., Rembach, A., Frugier, T., Li, Q.X., Ryan, T.M., McLean, C.A., Doecke, J.D., Rowe, C.C., et al. (2017). Biochemically-defined pools of amyloid-beta in sporadic Alzheimer’s disease: correlation with amyloid PET. Brain.

Roczniak-Ferguson, A., Petit, C.S., Froehlich, F., Qian, S., Ky, J., Angarola, B., Walther, T.C., and Ferguson, S.M. (2012). The transcription factor TFEB links mTORC1 signaling to transcriptional control of lysosome homeostasis. Sci Signal 5, ra42.

Rogaeva, E., Meng, Y., Lee, J.H., Gu, Y., Kawarai, T., Zou, F., Katayama, T., Baldwin, C.T., Cheng, R., Hasegawa, H., et al. (2007). The neuronal sortilin-related receptor SORL1 is genetically associated with Alzheimer disease. Nat Genet 39, 168–177.

Runz, H., Rietdorf, J., Tomic, I., de Bernard, M., Beyreuther, K., Pepperkok, R., and Hartmann, T. (2002). Inhibition of intracellular cholesterol transport alters presenilin localization and amyloid precursor protein processing in neuronal cells. J Neurosci 22, 1679–1689.

Selkoe, D.J. (2011). Alzheimer’s disease. Cold Spring Harbor perspectives in biology 3.

Settembre, C., Di Malta, C., Polito, V.A., Garcia Arencibia, M., Vetrini, F., Erdin, S., Erdin, S.U., Huynh, T., Medina, D., Colella, P., et al. (2011). TFEB links autophagy to lysosomal biogenesis. Science 332, 1429–1433.

Settembre, C., Zoncu, R., Medina, D.L., Vetrini, F., Erdin, S., Erdin, S., Huynh, T., Ferron, M., Karsenty, G., Vellard, M.C., et al. (2012). A lysosome-to-nucleus signalling mechanism senses and regulates the lysosome via mTOR and TFEB. Embo J 31, 1095–1108.

Shibata, M., Yamada, S., Kumar, S.R., Calero, M., Bading, J., Frangione, B., Holtzman, D.M., Miller, C.A., Strickland, D.K., Ghiso, J., et al. (2000). Clearance of Alzheimer’s amyloid-ss(1-40) peptide from brain by LDL receptor-related protein-1 at the bloodbrain barrier. J Clin Invest 106, 1489–1499.

Siegel, G., Gerber, H., Koch, P., Bruestle, O., Fraering, P.C., and Rajendran, L. (2017). The Alzheimer’s Disease gamma-Secretase Generates Higher 42:40 Ratios for beta-Amyloid Than for p3 Peptides. Cell Rep 19, 1967–1976.

Simons, M., Keller, P., De Strooper, B., Beyreuther, K., Dotti, C.G., and Simons, K. (1998). Cholesterol depletion inhibits the generation of beta-amyloid in hippocampal neurons. Proc Natl Acad Sci U S A 95, 6460–6464.

Small, S.A., and Gandy, S. (2006). Sorting through the cell biology of Alzheimer’s disease: intracellular pathways to pathogenesis. Neuron 52, 15–31.

Stanley, M., Macauley, S.L., Caesar, E.E., Koscal, L.J., Moritz, W., Robinson, G.O., Roh, J., Keyser, J., Jiang, H., and Holtzman, D.M. (2016). The Effects of Peripheral and Central High Insulin on Brain Insulin Signaling and Amyloid-beta in Young and Old APP/PS1 Mice. J Neurosci 36, 11704–11715.

Steculorum, S.M., Solas, M., and Bruning, J.C. (2014). The paradox of neuronal insulin action and resistance in the development of aging-associated diseases. Alzheimers Dement 10, S3–11.

Tajes, M., Gutierrez-Cuesta, J., Folch, J., Ortuno-Sahagun, D., Verdaguer, E., Jimenez, A., Junyent, F., Lau, A., Camins, A., and Pallas, M. (2010). Neuroprotective role of intermittent fasting in senescence-accelerated mice P8 (SAMP8). Exp Gerontol 45, 702–710.

Takahashi, R.H., Almeida, C.G., Kearney, P.F., Yu, F., Lin, M.T., Milner, T.A., and Gouras, G.K. (2004). Oligomerization of Alzheimer’s beta-amyloid within processes and synapses of cultured neurons and brain. J Neurosci 24, 3592–3599.

Tampellini, D., Magrane, J., Takahashi, R.H., Li, F., Lin, M.T., Almeida, C.G., and Gouras, G.K. (2007). Internalized antibodies to the Abeta domain of APP reduce neuronal Abeta and protect against synaptic alterations. J Biol Chem 282, 18895–18906.

Tanzi, R.E. (2005). The synaptic Abeta hypothesis of Alzheimer disease. Nat Neurosci 8, 977–979.

Tanzi, R.E., and Bertram, L. (2005). Twenty years of the Alzheimer’s disease amyloid hypothesis: a genetic perspective. Cell 120, 545–555.

Thinakaran, G., Teplow, D.B., Siman, R., Greenberg, B., and Sisodia, S.S. (1996). Metabolism of the “Swedish” amyloid precursor protein variant in neuro2a (N2a) cells. Evidence that cleavage at the “beta-secretase” site occurs in the golgi apparatus. J Biol Chem 271, 9390–9397.

Udayar, V., Buggia-Prevot, V., Guerreiro, R.L., Siegel, G., Rambabu, N., Soohoo, A.L., Ponnusamy, M., Siegenthaler, B., Bali, J., Aesg, et al. (2013). A paired RNAi and RabGAP overexpression screen identifies Rab11 as a regulator of beta-amyloid production. Cell Rep 5, 1536–1551.

Verdugo, M.E., and Ray, J. (1997). Age-related increase in activity of specific lysosomal enzymes in the human retinal pigment epithelium. Exp Eye Res 65, 231–240.

Villemagne, V.L., Burnham, S., Bourgeat, P., Brown, B., Ellis, K.A., Salvado, O., Szoeke, C., Macaulay, S.L., Martins, R., Maruff, P., et al. (2013). Amyloid beta deposition, neurodegeneration, and cognitive decline in sporadic Alzheimer’s disease: a prospective cohort study. Lancet Neurol 12, 357–367.

Wahrle, S., Das, P., Nyborg, A.C., McLendon, C., Shoji, M., Kawarabayashi, T., Younkin, L.H., Younkin, S.G., and Golde, T.E. (2002). Cholesterol-dependent gamma-secretase activity in buoyant cholesterol-rich membrane microdomains. Neurobiol Dis 9, 11–23.

Weller, R.O., Subash, M., Preston, S.D., Mazanti, I., and Carare, R.O. (2008). Perivascular drainage of amyloid-beta peptides from the brain and its failure in cerebral amyloid angiopathy and Alzheimer’s disease. Brain Pathol 18, 253–266.

Xiao, Q., Yan, P., Ma, X., Liu, H., Perez, R., Zhu, A., Gonzales, E., Tripoli, D.L., Czerniewski, L., Ballabio, A., et al. (2015). Neuronal-Targeted TFEB Accelerates Lysosomal Degradation of APP, Reducing Abeta Generation and Amyloid Plaque Pathogenesis. J Neurosci 35, 12137–12151.

Xu, W., Caracciolo, B., Wang, H.X., Winblad, B., Backman, L., Qiu, C., and Fratiglioni, L. (2010). Accelerated progression from mild cognitive impairment to dementia in people with diabetes. Diabetes 59, 2928–2935.

Zhang, Y., Nicholatos, J., Dreier, J.R., Ricoult, S.J., Widenmaier, S.B., Hotamisligil, G.S., Kwiatkowski, D.J., and Manning, B.D. (2014). Coordinated regulation of protein synthesis and degradation by mTORC1. Nature 513, 440–443.

Zhang, Y.D., and Zhao, J.J. (2015). TFEB Participates in the Abeta-Induced Pathogenesis of Alzheimer’s Disease by Regulating the Autophagy-Lysosome Pathway. DNA Cell Biol 34, 661–668.

