## Supplementary Figures for "Nutrient signaling pathways regulate amyloid clearance and synaptic loss in Alzheimer’s disease"

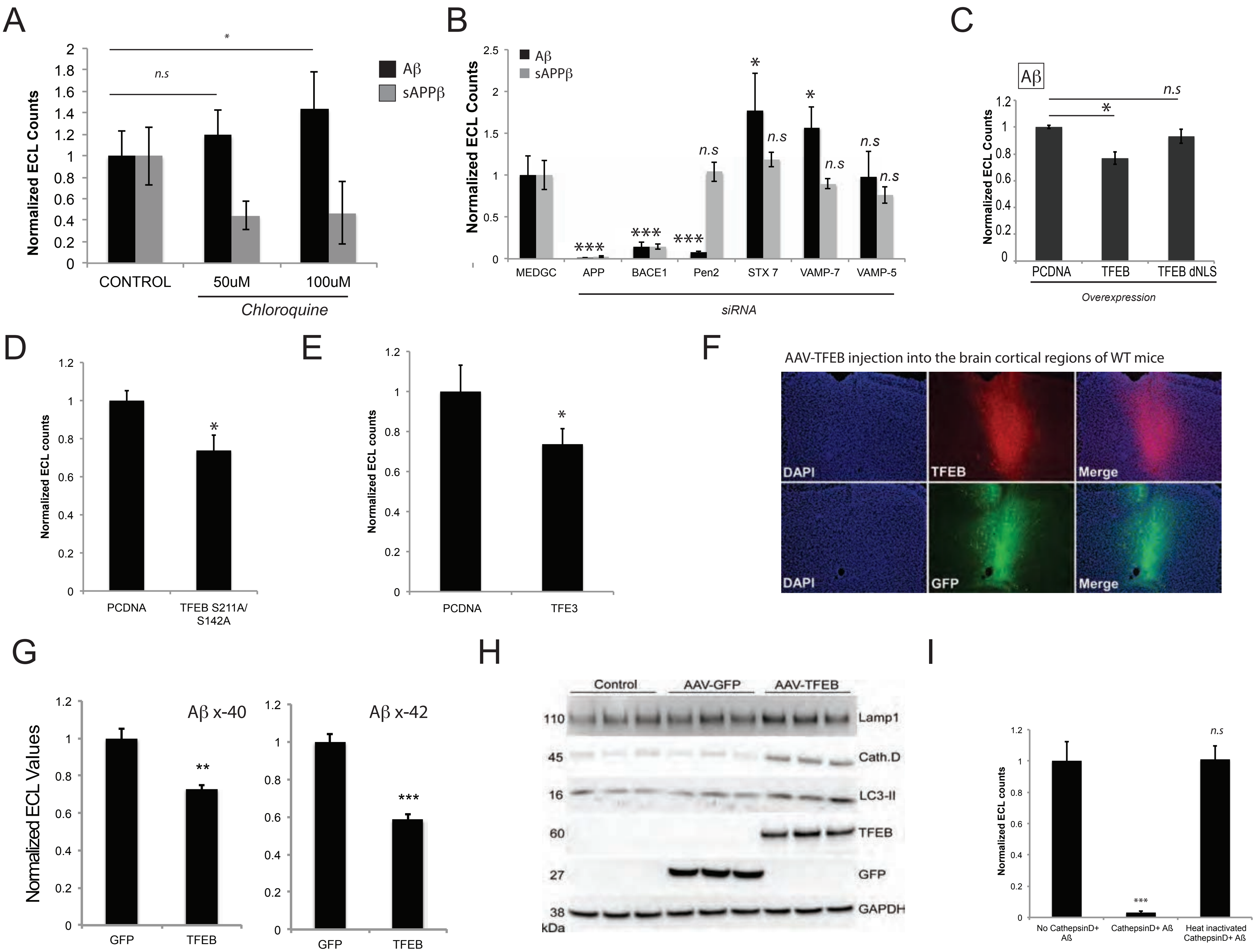


Mondal et al, SFig.1

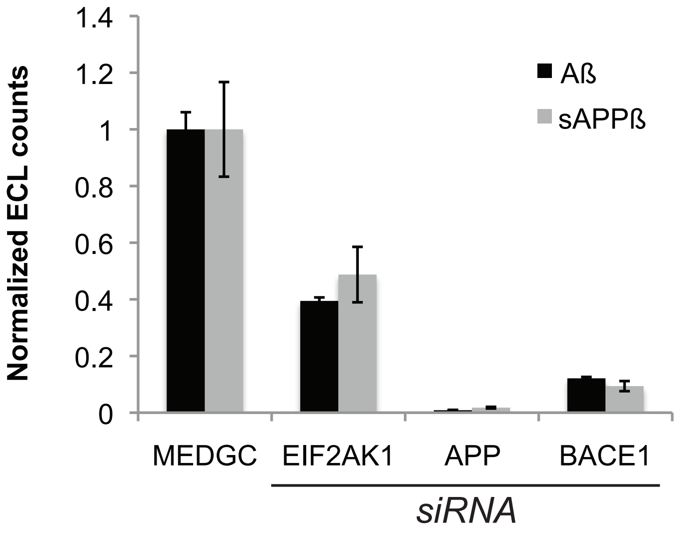


Mondal et al., SFig.2

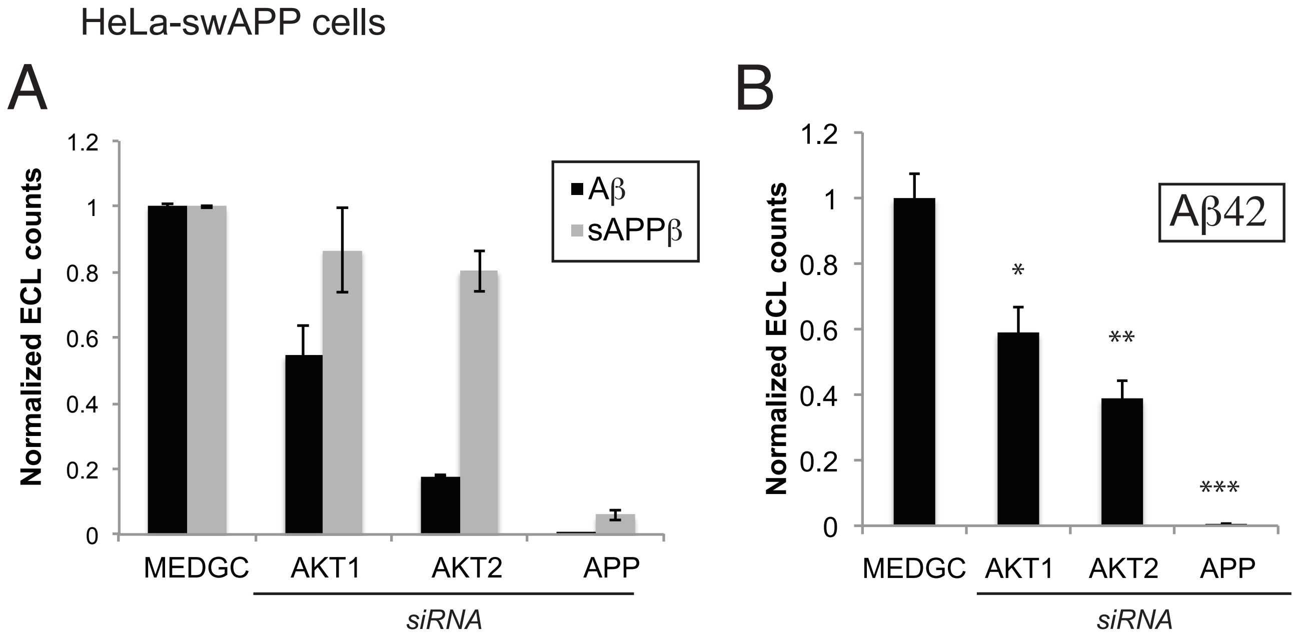


Mondal et al, SFig.3

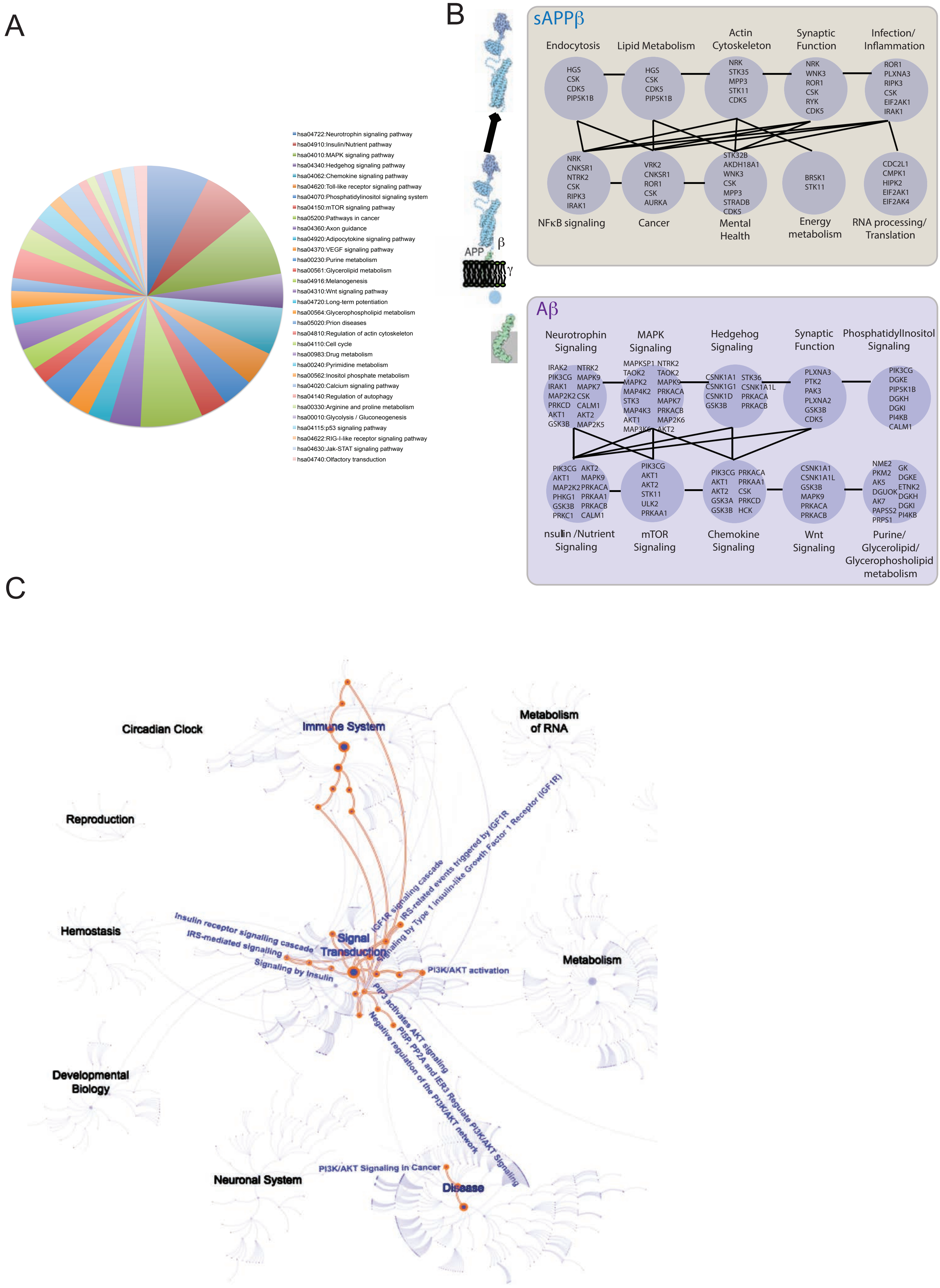


Mondal et al., SFig.4

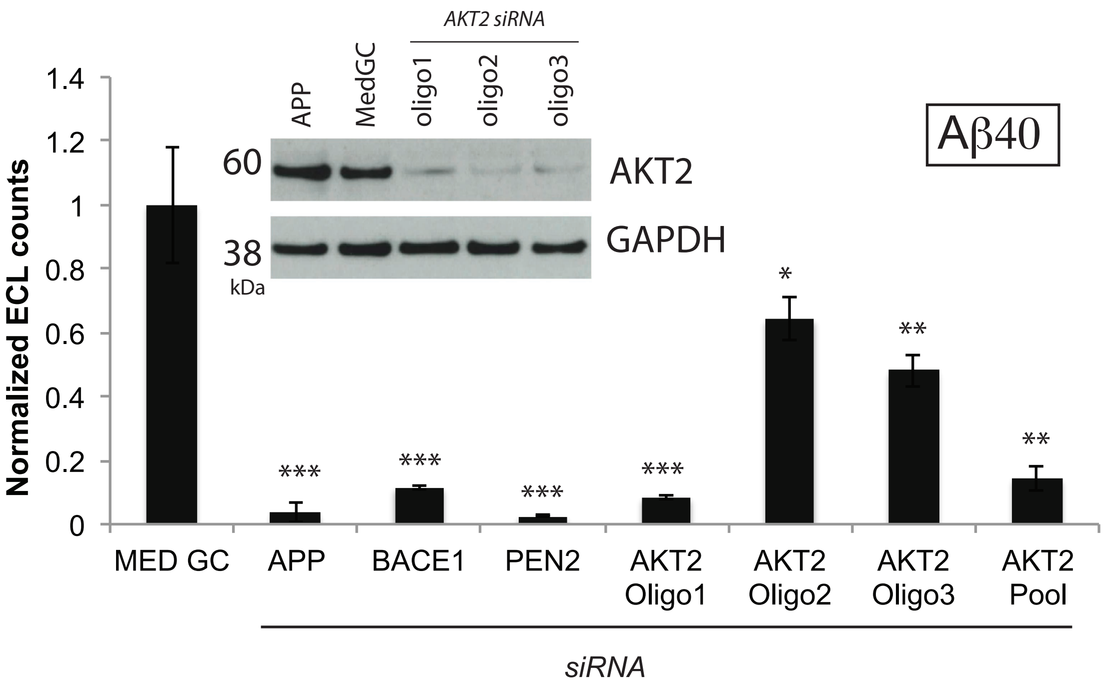


Mondal et al., SFig.5

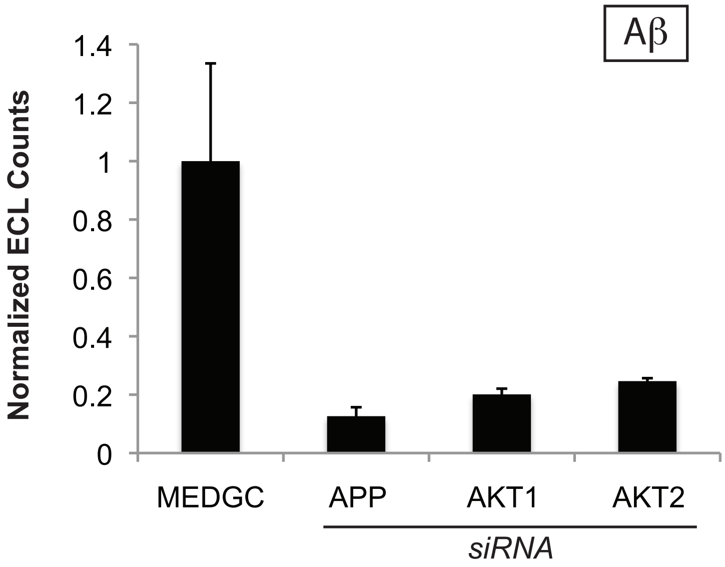


Mondal et al, SFig.6

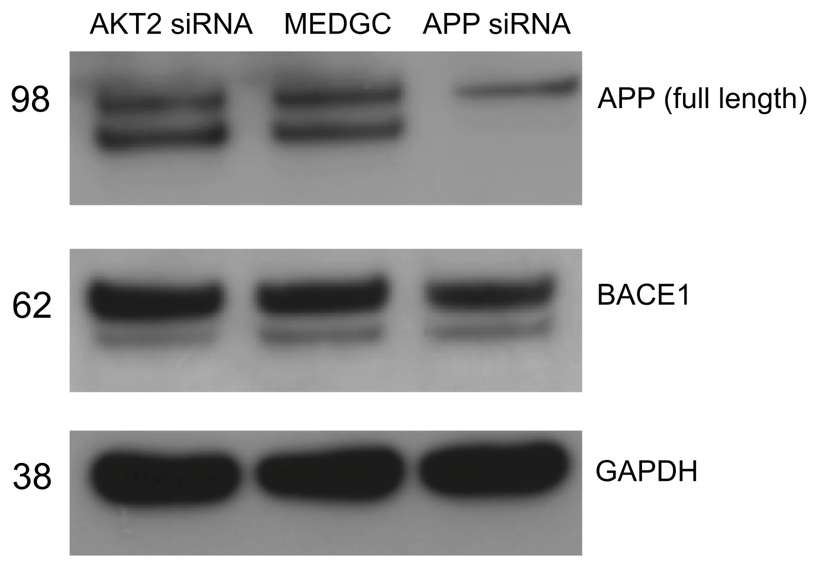


Mondal et al., SFig. 7

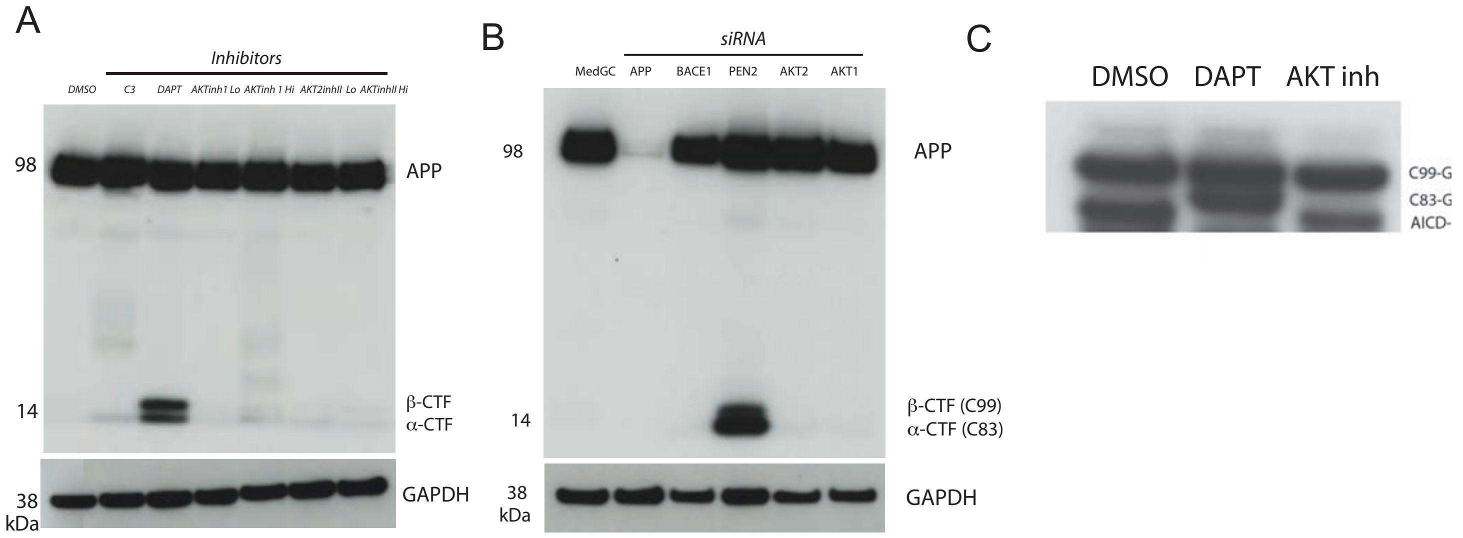


Mondal et al, SFig.8

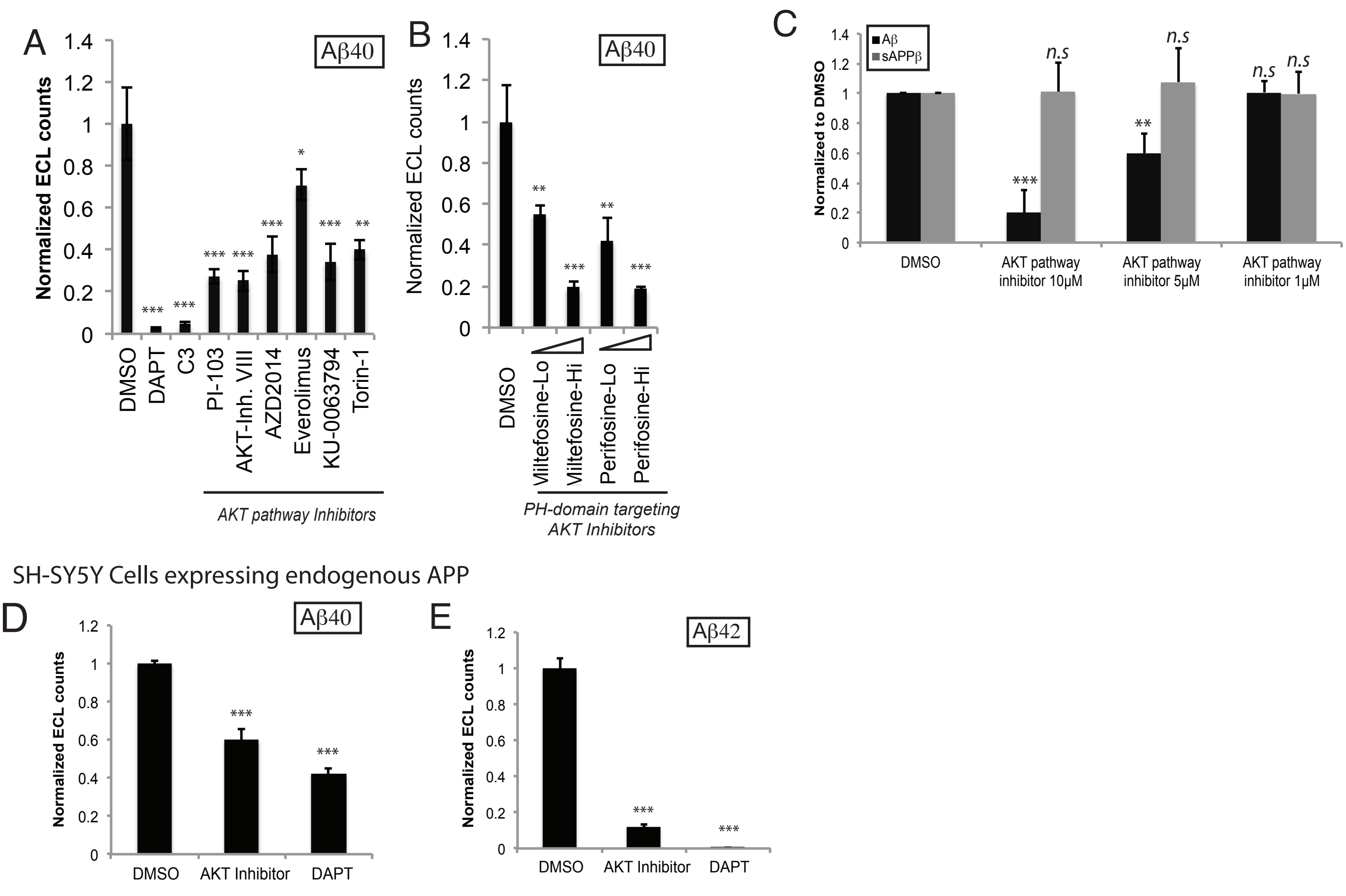


Mondal et al, SFig.9

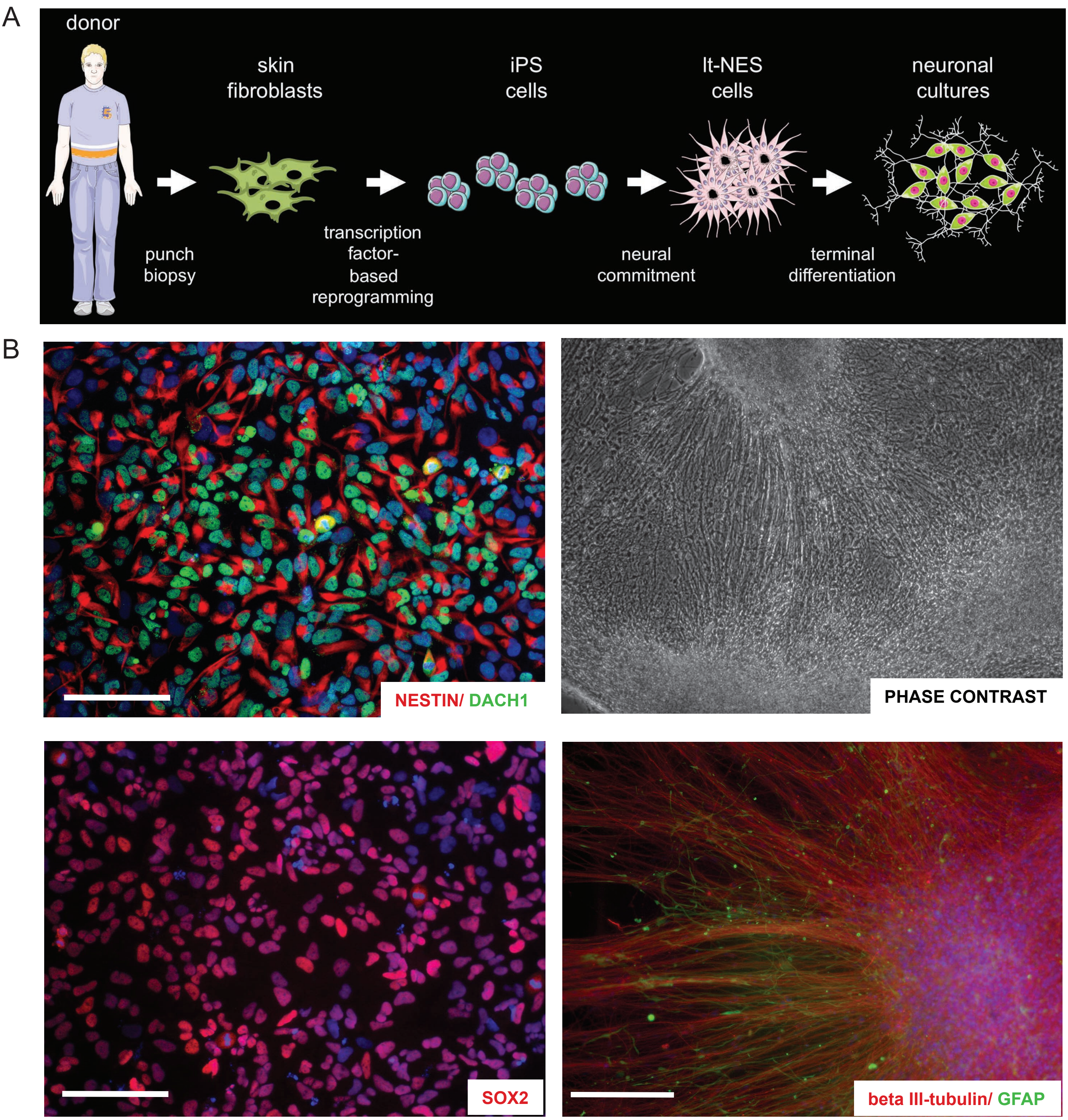


Mondal et al, SFig.10

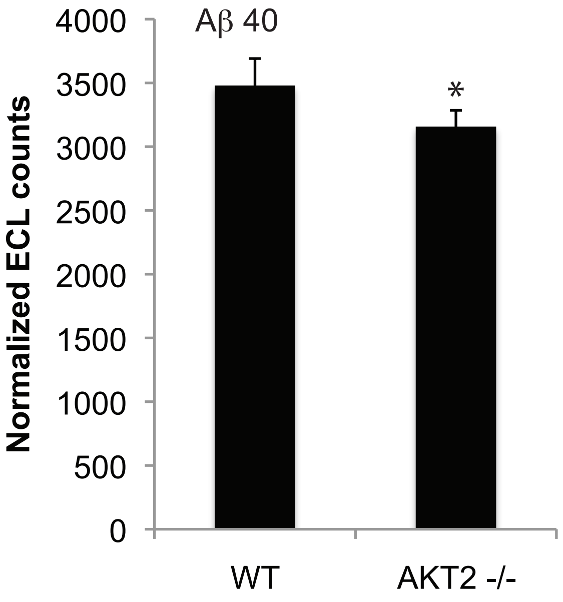


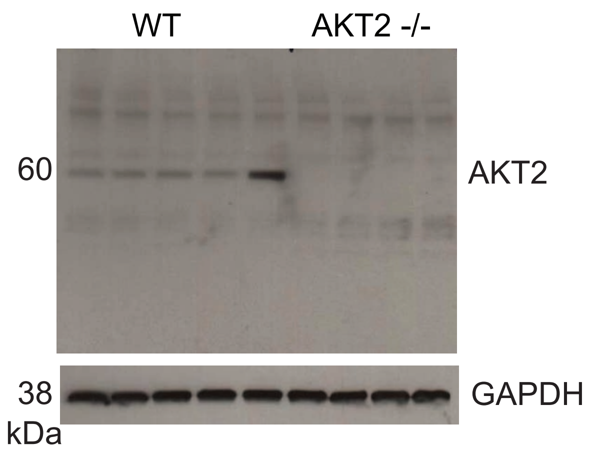


Mondal et al, SFig.11

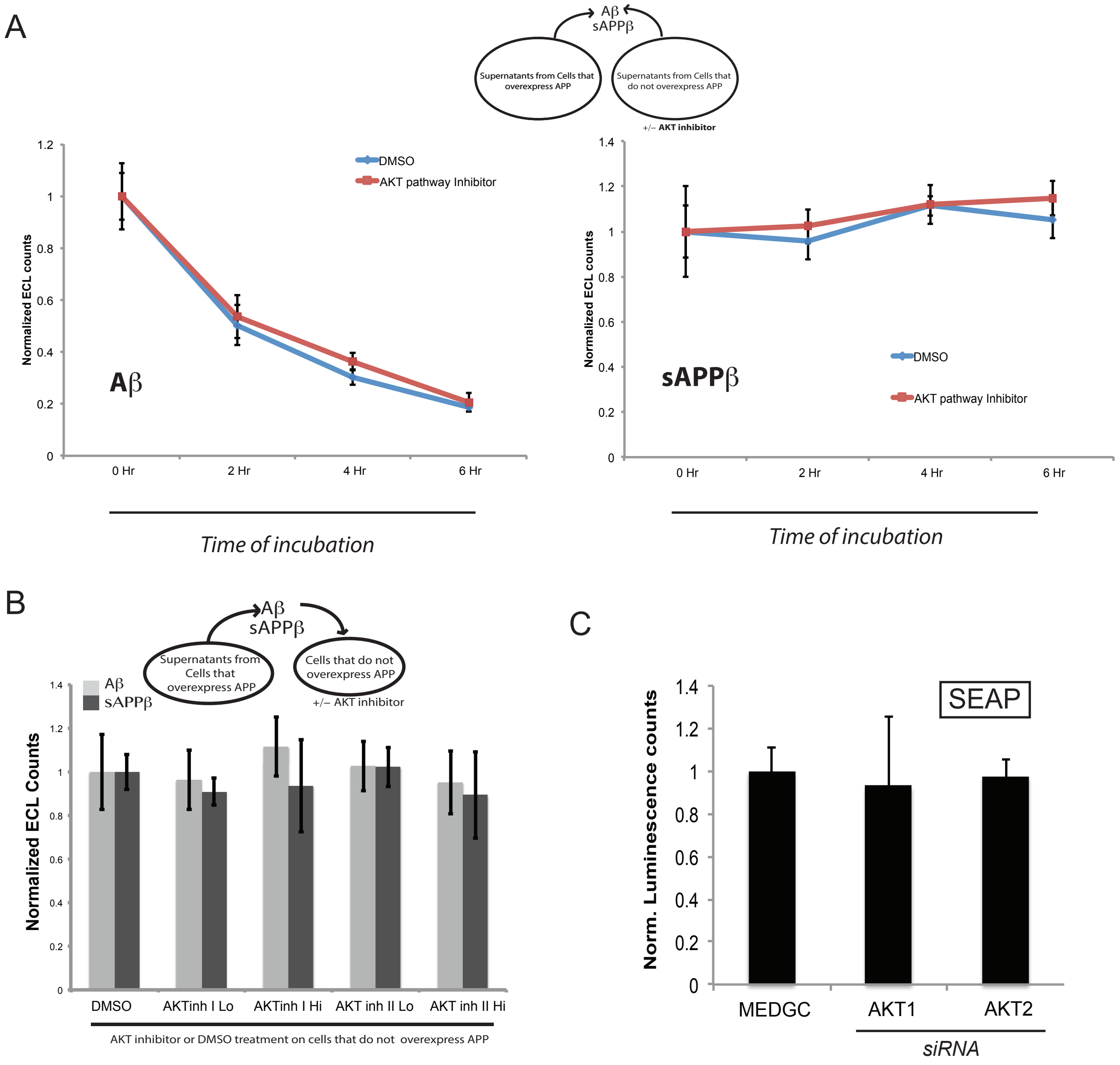


Mondal et al., SFig.12

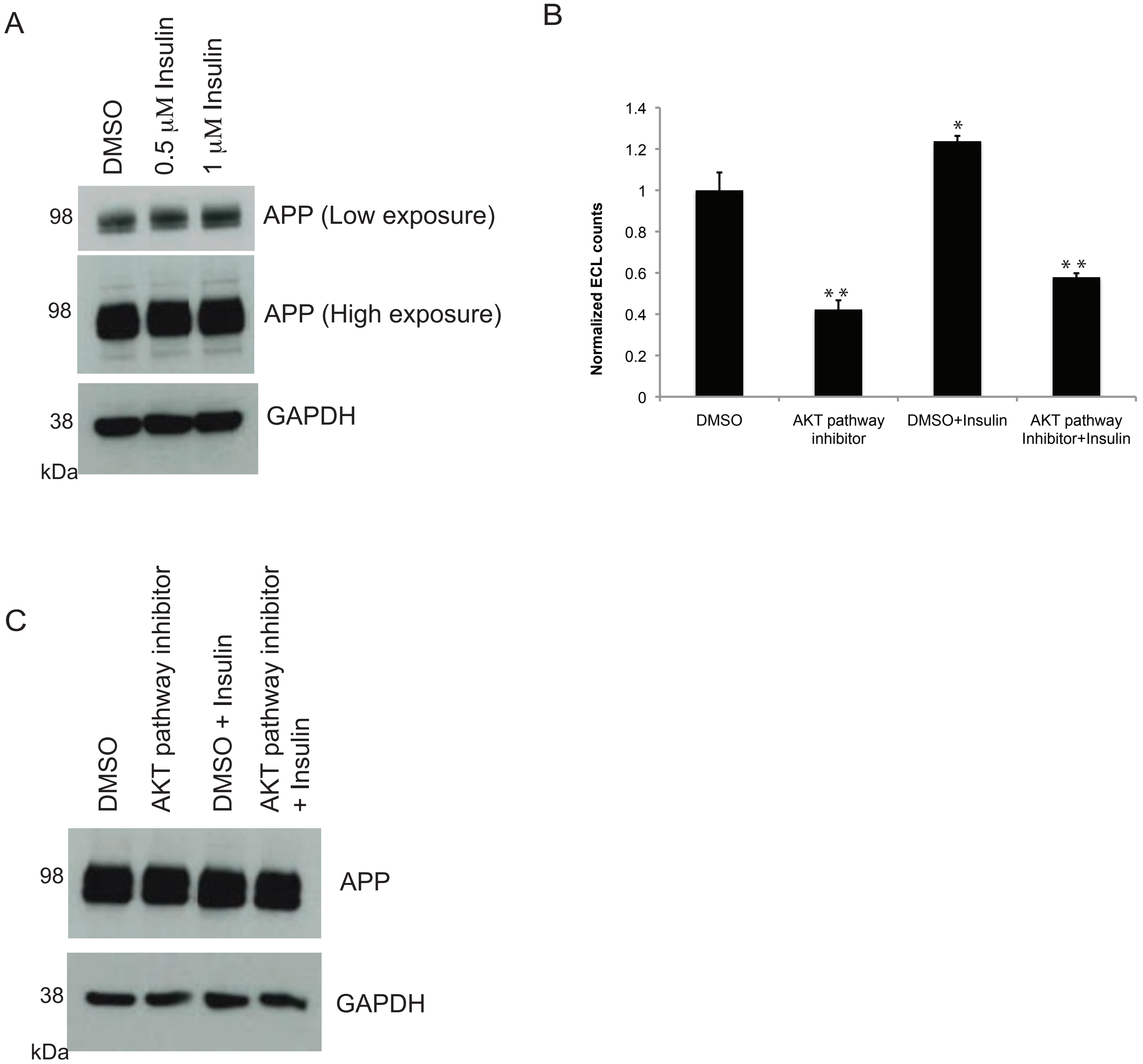


Mondal et al, SFig. 13

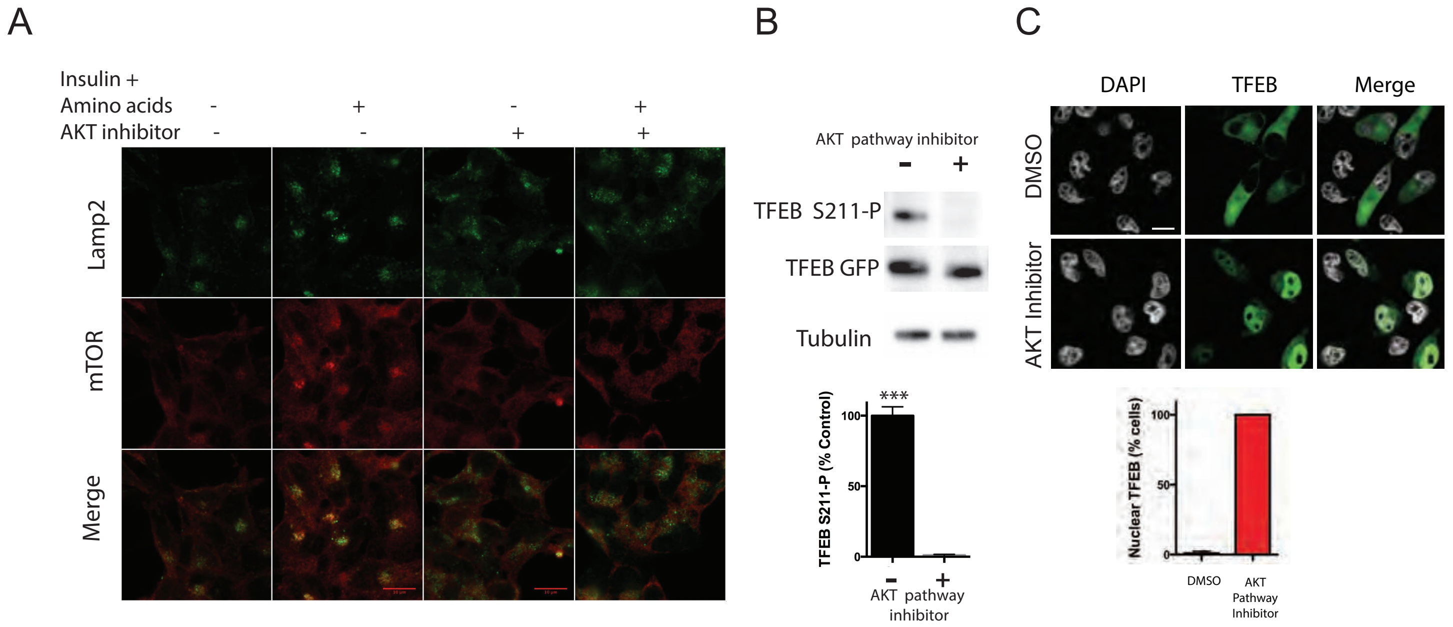


Mondal et al. SFig.14

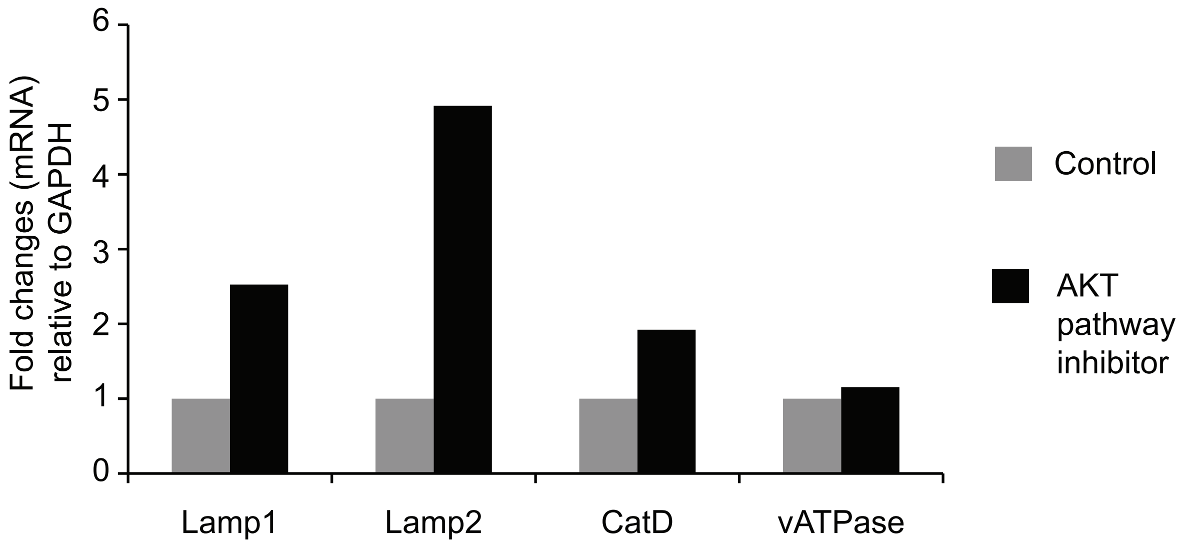


Mondal et al, SFig.15

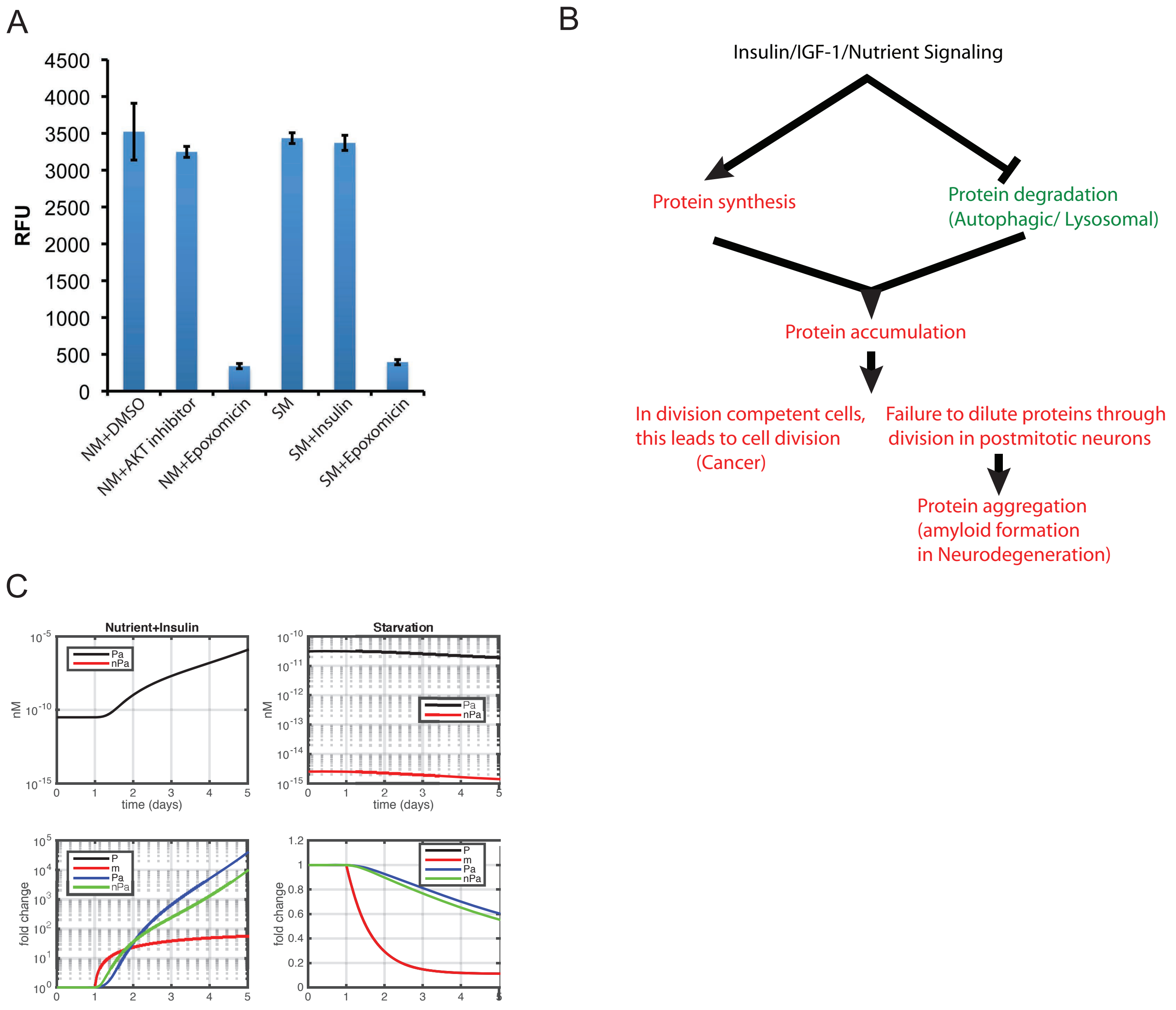


Mondal et al, SFig.16

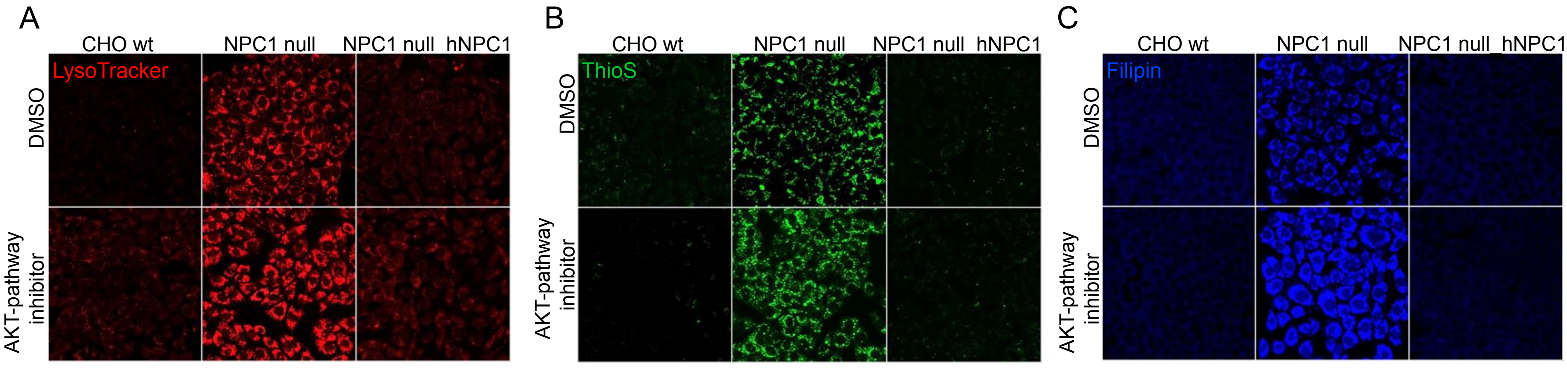


Mondal et al, SFig.17

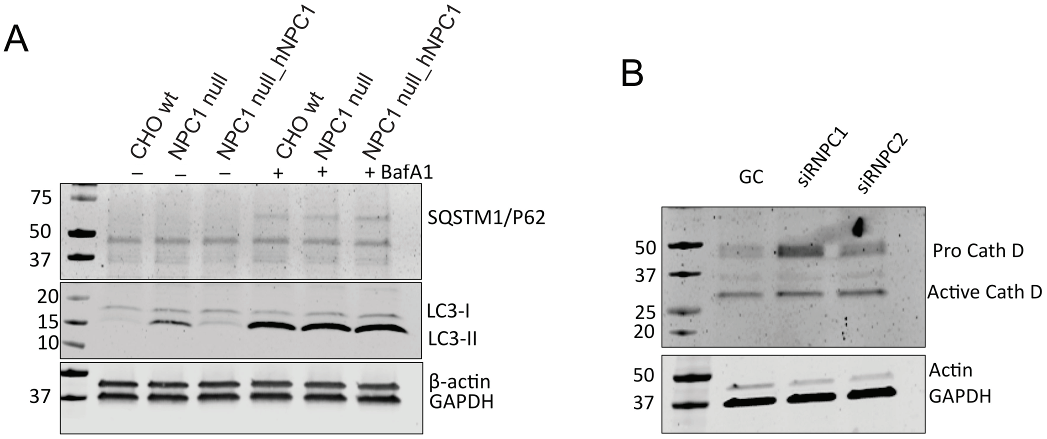


Mondal et al, SFig.18

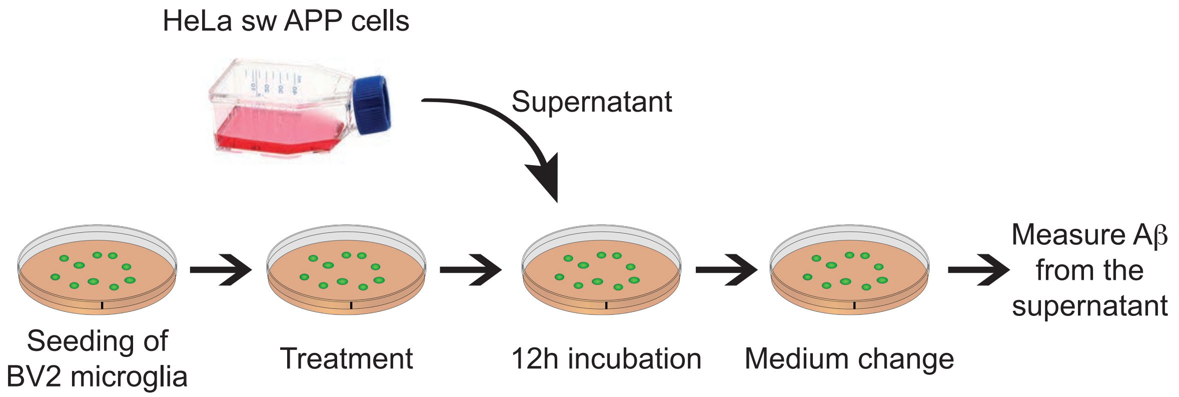


Mondal et al, SFig.19

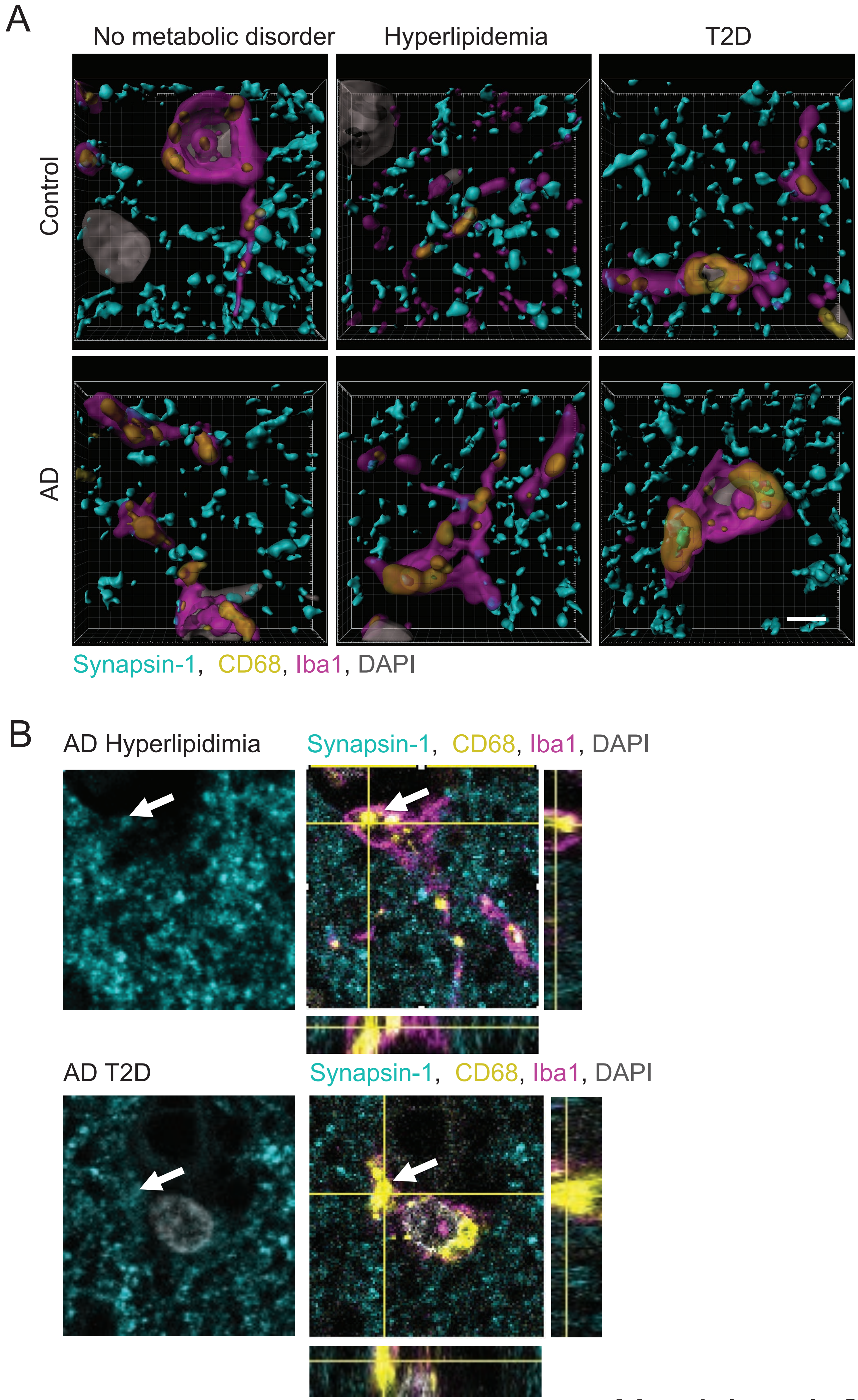


Mondal et al, SFig.20

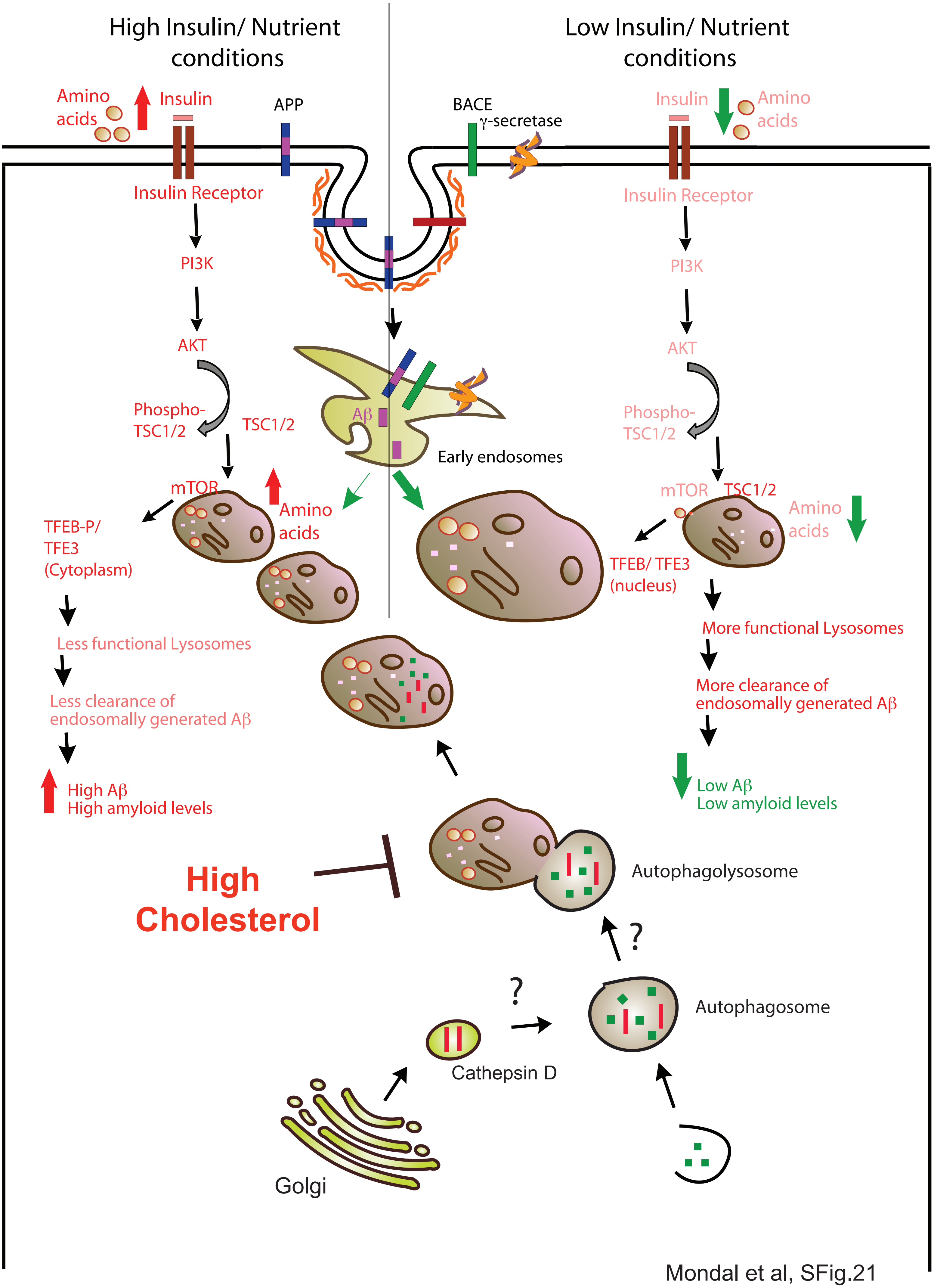


Mondal et al, SFig.21

**Reactome Pathway Enrichment Analysis**

| Pathway name | FDR |
|---|---|
| Signaling by Interleukins | 3.75E-12 |
| Diseases of signal transduction | 1.72E-10 |
| Signaling by SCF-KIT | 2.59E-10 |
| Insulin receptor signalling cascade | 3.04E-10 |
| Signaling by VEGF | 3.04E-10 |
| Signaling by EGFR | 3.04E-10 |
| PIP3 activates AKT signaling | 3.04E-10 |
| IRS-mediated signalling | 3.04E-10 |
| Negative regulation of the PI3K/AKT network | 3.04E-10 |
| Signaling by Insulin receptor | 3.05E-10 |
| IRS-related events triggered by IGF1R | 3.05E-10 |
| PI3K/AKT activation | 3.05E-10 |
| GAB1 signalosome | 3.05E-10 |
| IGF1R signaling cascade | 3.05E-10 |
| Signaling by Type 1 Insulin-like Growth Factor 1 Receptor (IGF1R) | 3.19E-10 |
| Downstream signal transduction | 3.90E-10 |
| VEGFA-VEGFR2 Pathway | 4.34E-10 |
| Signaling by PDGF | 4.34E-10 |
| DAP12 signaling | 4.34E-10 |
| Downstream signaling events of B Cell Receptor (BCR) | 6.34E-10 |
| PI5P, PP2A and IER3 Regulate PI3K/AKT Signaling | 6.34E-10 |
| DAP12 interactions | 9.63E-10 |
| MAPK1/MAPK3 signaling | 1.22E-09 |
| PI3K/AKT Signaling in Cancer | 1.22E-09 |

Mondal et al, STable 1

### Legends to Supplementary Figures

**Supplementary Figure 1: Replication and validation of Aβ clearance by lysosomes**
**A.** Cells treated with two different concentrations of Chloroquine and probed for Aβ (black bars) or sAPPβ levels (grey bars). *p<0.05. n.s indicates not significant. **B.** Aβ and sAPPβ levels after silencing of Vamp-7, Syntaxin-7 and Vamp-5 using specific siRNAs along with controls (MedGC scrambled oligos, APP, BACE1 and Pen-2 specific siRNAs). **C.** HeLa-sweAPP cells transfected with TFEB and TFEB dNLS and assayed for Aβ levels. PCDNA used as a negative control *p<.05. **D.** HeLa-sweAPP cells transfected with TFEB and TFEB S211A/S142A and assayed for Aβ levels. PCDNA used as a negative control *p<.05. **E.** HeLa-sweAPP cells transfected with TFE3 and assayed for Aβ levels. PCDNA used as a negative control *p<.05. **F.** Cortex of WT mice transduced with an AAV-TFEB (red, top row) and AAV-GFP vector (green, bottom row, used as negative control). Nuclei are stained with DAPI (blue). **G.** Aβ extracted from cortex of WT mice injected with AAV-TFEB and AAV-GFP and assayed for Aβ levels. **p<.005, ***<.0005. **H.** Cortical tissue from control, AAV-GFP-injected and AAV-TFEB-injected mice was analysed by western blot to confirm the expression of the transgenes as well as the increased lysosomal biogenesis (assessed by LAMP-1 expression) induced by TFEB overexpression. CathepsinD and LC3-II levels were also determined. GAPDH was used as a loading control. **I.** Recombinant human Cathepsin D (0.625ng/μl) was incubated with synthetic Aβ. Following the incubation electrochemiluminescence assay was performed to determine the amount of degraded Aβ. Synthetic Aβ without Cathepsin D treatment was used as a negative control. Heat inactivated Cathepsin D was also incubated with synthetic Aβ. Heat inactivated Cathepsin D did not degrade Aβ. Error bars indicate S.D.

**Supplementary Figure 2:** Silencing of EIF2AK1 reduces both sAPPβ and Aβ levels. HeLa-sweAPP cells were transfected with siRNA pools against EIF2AK1, APP or BACE1 (positive controls) or with a control-siRNA (MEDGC) and the conditioned media were assayed for Aβ (black) and sAPPβ (grey) by ECL-multiplex assay. Error bars indicate SEM.

**Supplementary Figure 3: A.** HeLa-sweAPP cells transfected with siRNA pools against AKT1, AKT2 and APP (positive control) along with a scrambled MEDGC oligo as negative control and assayed for Aβ (black) and sAPPβ (grey) levels. **B.** HeLa-sweAPP cells transfected with siRNA pool against AKT1, AKT2 and APP (positive control) along with scrambled MEDGC oligo as negative control and assayed for Aβ42 *p<.05,** <.005, ***<.0005. Error bars indicate S.D.

**Supplementary Figure 4: A, B.** Bioinformatics analysis of siRNA screen hits identifies various signalling networks. Insulin/nutrient signalling pathway are seen as one of the top clusters. **C.** Reactome pathway enrichment analysis of the aging-similar genes showed a prominent contribution from the IIN-AKT pathways.

**Supplementary Figure 5:** HeLa-sweAPP cells transfected with siRNA pool against APP, BACE1, PEN2, AKT2 or individual siRNA against AKT2 or MEDGC as negative control and probed for Aβ levels. AKT2 levels were analyzed using AKT2 specific antibodies. GAPDH was used as a protein loading control. Error bars indicate S:D.

**Supplementary Figure 6:** Silencing of AKT1 and AKT2 reduces Aβ levels in cell lysates. HeLa-sweAPP cells were transfected with siRNA pool against AKT1, AKT2 or APP (positive control) or with a control-siRNA (MEDGC) and lysates of transfected cells were assayed for Aβ by ECL-assay (Error bars indicate SEM).

**Supplementary Figure 7:** Silencing of AKT2 does not influence APP or BACE1 levels. HeLa-sweAPP cells were transfected with siRNA pool against AKT2 or APP (positive control) or with a control-siRNA (MEDGC) and lysates of transfected cells were immunoblotted against APP and BACE1. GAPDH was used as a protein loading control.

**Supplementary Figure 8:** **A**. HeLa-sweAPP cells treated with β-secretase inhibitor (C3), γ-secretase inhibitor (DAPT) and AKT inhibitors (AKT inh I, AKT inh II. Lo-1μM and Hi-10 μM) and probed for APP cleavage products. DAPT treatment leads to a specific accumulation of C-terminal fragments of APP, which is not seen when the cells are treated with AKT inhibitors, C3 or DMSO (negative control). **B.** RNAi silencing of AKT1, AKT2 and as controls scrambled (MedGC), APP, BACE1 or PEN2 were performed in HeLa-sweAPP cells and probed for APP cleavage products using the C-terminal antibody of APP. The γ-secretase subunit, PEN2 silencing leads to a specific accumulation of C-terminal fragments of APP, which is not seen when the cells are treated with AKT1 or AKT2 siRNAs. GAPDH is used as a loading control. **C.** HeLa-sweAPP cells transfected with C99-GFP plasmid were treated for 12 h with either AKT inhibitor VIII, DMSO (negative control) or DAPT, the γ-secretase inhibitor as positive control. Cell lysates were analysed by Western blot probed with anti-GFP antibody. AICD-GFP levels are not reduced upon AKT-inhibitor treatment. Alternatively, DAPT treatment reduced the formation of AICD-GFP but instead produces C83-GFP due to the μ-cleavage of C99-GFP. Error bars are S.D.

**Supplementary Figure 9:** **A.** HeLa-sweAPP cells treated with different AKT pathway inhibitors and assayed for Aβ levels. DMSO treatment was used as negative control. γ-Secretase inhibitor DAPT and β-secretase inhibitor C3 treatment were used as positive controls.*p<.05,** <.005, ***<.0005. **B.** HeLa-sweAPP cells treated with FDA approved AKT inhibitors (Miltefosine and Perifosine) using different concentrations (lo-10μM, and Hi- 25μM for Miltefosine and 50μM for Perifosine) and assayed for Aβ levels. **C.** HeLa-sweAPP cells treated with different concentrations of AKT pathway inhibitor and probed for Aβ (black bars) or sAPPβ (grey bars). DMSO was used as a negative control. **D.** SH-SY5Y cells treated with AKT inhibitor and assayed for Aβ levels. ***p<.0005. DMSO and DAPT treatment were used as negative and positive controls respectively. **E.** HeLa-sweAPP cells treated with AKT inhibitor and assayed for Aβ42 levels. DMSO and DAPT treatment were used as negative and positive controls respectively. ***p<.0001. Error bars are S.D.

**Supplementary Figure 10:** **A.** Skin fibroblasts were derived from a healthy donor and iPS cells were established using the reprogramming factors OCT4, SOX2, KLF4 and c-MYC. From iPS cells, neuro epithelial stem cells, a long-term, self-renewing neural stem cell population, were established. Terminal differentiation into neuronal cultures was initiated by growth factor withdrawal and neurons terminally differentiated for 4 weeks before applying compounds for Aβ measurements. The cartoon was produced using Servier Medical Art (www.servier.com). **B.** Human iPSC-derived neuro epithelial stem cells express the neural stem cell-associated transcription factors SOX2 and DACH1 as well as the intermediate filament NESTIN. IPSC-derived differentiated neuronal cultures mainly consist of neurons expressing beta III-tubulin and MAP2ab. A smaller fraction of GFAP-expressing astrocytes is also present in the cultures.

**Supplementary Figure 11:** Cortical extract of wild type (WT) mice or mice lacking AKT2 assayed for Aβ levels *p<.05. Westen blot image of cortical extracts shows the absence of AKT2 in AKT2-/- mice compared to WT mice. GAPDH was used as a loading control.

**Supplementary Figure 12: A.** Untransfected HeLa cells that do not overexpress APP were treated with AKT pathway inhibitor. Conditioned medium obtained from AKT pathway inhibited HeLa cells were mixed with conditioned medium from HeLa-sweAPP cells (cells that overexpress APP and have robust levels of Aβ and sAPPβ) for different time points and incubated at 37°C. The Aβ and sAPPβ levels that remained after incubation were measured using electrochemiluminescence detection and normalized to the values from DMSO treated conditions. **B.** Untransfected HeLa cells that do not overexpress APP and hence have no detectable Aβ and sAPPβ were treated with either AKT inh I or AKT inh II at two different concentrations (Lo-1μM and Hi-10μM) followed by replacement of medium with conditioned medium from HeLa-sweAPP cells (cells that over ever express APP and have robust levels of Aβ and sAPPβ) for 3 h. The Aβ/ sAPPβ levels that remain after the incubation were measured using electrochemiluminescence detection and normalized to the values from DMSO treated conditions. **C.** HeLa-sweAPP cells transfected with siRNA pool against AKT1, AKT2, MEDGC followed by transfection with SEAP and assayed for released SEAP.

**Supplementary Figure 13: A.** iPSC treated with two different concentrations of insulin and probed for APP levels. DMSO treatment was used as negative control. GAPDH was used as a loading control. **B**. Hela-sweAPP cells treated with insulin in the presence or absence of AKT pathway inhibitor and probed for intracellular Aβ levels. DMSO treatment was used as negative control. *p<.05,**<.005 **C.** Hela-sweAPP cells treated with insulin in the presence or absence of AKT pathway inhibitor and probed APP levels. DMSO treatment was used as negative control. GAPDH was used as a loading control.

**Supplementary Figure 14: A.** HEK293 cells were serum starved for 10 h followed by amino acid starvation for 1h. Cells were then treated with AKT pathway inhibitor (10 μM) for 2.5 h. For insulin treatment, cells were pretreated (2 h) with AKT pathway inhibitor, and stimulated with medium containing insulin (1 μM) and AKT pathway inhibitor for 30 min. DMSO treatment was used as negative control. Cells were co-labeled for Lamp2 (green) and mTOR (red). Scale bar is 10 μm. **B.** Immunoblot analysis of TFEB-S211 phosphorylation (HeLa TFEB-GFP stable cell line) following 2 hour treatment with vehicle (DMSO) or 10μM Akt inhibitor (n=3, mean+/-SEM, p=0.002, t-test). Quantification of TFEB S211-P in the presence or absence of AKT pathway inhibitor. **C.** HeLa TFEB-GFP cells serum deprived (starved) treated with insulin in the presence of AKT inhibitor or DMSO (negative control). Cells were imaged for GFP (TFEB), DAPI (Nucleus) and lysotracker (red). Scale bar is 20 μm.

**Supplementary Figure 15:** RT-PCR analysis of Lamp1, Lamp2, CatD and v-ATPase after treatment with AKT pathway inhibitor (black bars) DMSO treatment (grey bars) was used as a negative control.

**Supplementary Figure 16**: **A.** Measurement of proteasomal activity in the presence of AKT pathway inhibitor or insulin. DMSO treatment was used as negative control and epoxomicin treatment was used as positive control. NM-Normal medium. SM-Starvation medium (devoid of serum). **B.** Hypothetical scheme for protein aggregation in post-mitotic cells (neurons). **C.**

Simulated effects of increased nutrition and insulin (left column) and of starvation (right column) on the aggregation of WT Aβ. The treatment simulations started at the beginning of day 2. The simulations were performed using protein aggregation kinetics of Aβ42 (Meisl *et al.*, 2014).

**Supplementary Figure 17**: **A, B, C.** WT, NPC1 null and NPC1 null cells stably expressed human NPC1 were treated with AKT-pathway inhibitor and stained for (A) Lysotracker (red), (B) ThioS (green) and (C) Filipin (blue).

**Supplementary Figure 18: A.** Western blot analysis of SQSTM1/P62 and LC3-I/LC3-II were performed on cell lysates from CHO wt, NPC1 knockout CHO cells and NPC1 knockout CHO cells stably expressing human NPC1. β-actin and GAPDH were used as loading controls. **B.** siRNA based gene knock down of NPC1 and NPC2 were performed on HeLa cells and Cathepsin D levels were analyzed by western blot. β-actin and GAPDH were used as loading controls.

**Supplementary Figure 19:** Schematic representation of Aβ uptake assay in microglia.

**Supplementary Figure 20: A.** Imaris 3D reconstructions of confocal images. Microglia are represented by Iba1 (magenta) and CD68 (yellow), and pre-synaptic marker Synapsin-1 (cyan), counterstained with DAPI for nuclei (grey). Scale bar 3 microns. **B.** Magnified representative confocal images showing 20x20x4 micron regions of interest stained with Iba1 (magenta) and CD68 (yellow), and pre-synaptic marker Synapsin-1 (cyan), counterstained with DAPI for nuclei (grey). Synapsin-1 inclusions can be seen in microglial cells of AD individuals by orthogonal view.

**Supplementary Figure 21:** Schematic representation of regulation of amyloid levels depends on insulin signalling/Nutrient sensing pathway.

**STable 1:** Bioinformatics analysis of human tissue-specific transcriptome analysis of the ageing-similar genes showed a prominent contribution from the IIN-AKT pathways among the 25 most enriched pathways
