## Supplementary methods for "Nutrient signaling pathways regulate amyloid clearance and synaptic loss in Alzheimer’s disease"

### Cells

HeLa-sweAPP cells were cultured in DMEM (Gibco), supplemented with 10% FCS (Gibco), 1% penicillin, 1% streptomycin (from 10 mg/ml stock, Gibco), 0.1% G418 (from 50µg/µl stock, Carl Roth) and 0.1% Zeocin (from 100mg/ml stock, Invitrogen). HeLa cells were cultured in DMEM (Gibco), supplemented with 10% FCS (Gibco), 1% penicillin, 1% streptomycin (from 10 mg/ml stock, Gibco). HEK293 cells were cultured in DMEM (Gibco), supplemented with 10% FCS (Gibco), 1% penicillin, 1% streptomycin (from 10mg/ml stock, Gibco). SHSY5Y cells were cultured in DMEM (Gibco), supplemented with 10% FCS (Gibco), 1% penicillin, 1% streptomycin (from 10mg/ml stock, Gibco). BV2 microglia cells and HEK293 cells were cultured in DMEM (Gibco), supplemented with 10% FCS (Gibco), 1% penicillin, 1% streptomycin (from 10mg/ml stock, Gibco). CHO cells were cultured in DMEM/F12 medium (ThermoFisher scientific), supplemented with 10% FCS (Gibco), 1% Glutamax (Gibco).

### siRNA

siRNAs were purchased from Invitrogen (stealth siRNA).

### RNAi mediated screen

RNAi mediated screen was performed in HeLa-sweAPP cells. siRNA against specific genes were used for effective knockdowns. Oligofectamine (Invitrogen) was used for transfection. 5 nM final siRNA were prepared in Optimem (Gibco) to the final volume of 5 µl (mix 1) and incubated for 5 min at room temperature. 0.3 µl of Oligofectamine (Invitrogen) was mixed with 4.7 µl of Optimem (mix 2) and incubated at room temperature for 5 min. Mix 1 was combined with mix 2 and incubated for 20 min at room temperature. 10 µl of the transfection mix was added to each well of a 96 well plate, followed by 3000 cells in 100 µl of culture medium. Each siRNA transfection was performed in triplicate. 24 h after transfection, medium was replaced with 100 µl of fresh medium. 69 h after transfection, medium was again replaced with 100 µl fresh medium containing 10 % Alamar blue (AbD Serotec) in order to check cell viability. 72 h after transfection, Alamar blue measurements were taken using SpectraMAX GeminiXS (Molecular Devices) at an excitation wavelength of 544 nm and emission at 590 nm. Supernatant was collected and frozen at -20°C. The cells in the transfected plate were lysed in 50 μl of lysis buffer, incubated for 20 min on ice and stored at -20°C.

**Kinases Inhibitor screen**

Kinases inhibitor screen was performed in HeLa-sweAPP cells. 20,000 cells were seeded in each well of a 96 well plate (Nunc) a day prior to inhibitor treatment. The cells were treated with 10 μM concentration of each inhibitors (Calbiochem) for 4 h along with 10% Alamar blue (AbD Serotec). 4 h after treatment, Alamar blue measurement was taken using SpectraMAX GeminiXS (Molecular Devices) at an excitation wavelength of 544 nm and emission at 590 nm. Supernatant was collected and frozen at -20°C. The cells were then lysed with 50 μl of lysis buffer, incubated for 20 min on ice and stored at -20°C.

**Electrochemiluminescence detection of sAPPβ and Aβ from supernatants/ cell lysates**

Electrochemiluminescence assay was performed to determine the amount of Aβ present in the supernatants or cells lysates. Each well of the pre-coated plate was blocked using 35 μl TBST (Tris Buffered Saline containing Tween), containing 3% blocker A, for 1 h at room temperature with shaking, followed by 4 times washing in TBST. After washing, 10 μl of the supernatants or cell lysates was added to each well along with 10 μl detection antibody followed by incubation for 2 h at room temperature with shaking. Wells were washed 3 times with TBST and detection was performed using 35 μl of 2x MSDT read buffer.

**Quantitative RT-PCR analysis**

72 h after siRNA transfection, total RNA was isolated using the RNeasy Plus Mini kit (Qiagen). Purity of RNA (A260/A280, A260/A230) and concentration was determined using a Nanodrop spectrophotometer. 2 μg of total RNA was used for reverse transcription with oligo-dT primer using Superscript first-strand synthesis system for RT-PCR (Invitrogen) according to the manufacturer's recommended protocol. PCR was performed using TaqMan custom array (Applied Biosystems) following manufacturer's instructions.

Expression levels of genes were normalized against GAPDH controls. Levels of MedGC cDNA as an internal control were normalized to GAPDH cDNA according to the ΔΔCt method using the equation 2-(Ctsamples-CtGAPDH) treatment-(CtMed-GC− CtGAPDH) control.

**Western blotting**

Cells either transfected with specific siRNA against APP, BACE1, AKT1, AKT2, TFEB and the γ-secretase components along with MEDGC scrambled oligos as control or treated with various inhibitors (C3, DAPT) were lysed in buffer containing 2% NP-40 and 0.2% SDS and protease inhibitors (Roche). Equal amounts of the lysate (according to the protein content quantified by BCA assay (Pierce)) were run on 4-12% BIS-TRIS gels (Invitrogen). The gel was blotted onto a nitrocellulose membrane (BioRad) and probed with the respective antibodies.

Anti-APP, C-Terminal antibody: SIGMA (F3165-1MG) or the Aβ-region recognizing 6E10 Antibody; Anti-BACE1 antibody: Prosci incorporation (2235); GAPDH antibody: Meridian Science, Inc; AKT1, AKT2 antibodies: Cell signaling; Nicastrin antibody: 9C3; Anti GFP antibody: Roche; β-Actin: Sigma, TFEB, TSC2, p-TSC2, mTOR, p-S6K, S6K, P-4E-BP1, 4E-BP1, SQSTM1/P62: Cell signalling. LAMP2 antibody: Santacruz. NPC1, Cathepsin D: Abcam. Lamp2 for mouse: DSHB.

For the *in vivo* experiment, cortical tissue from control, AAV-GFP-injected and AAV-TFEB-injected mice was dissected around the area of injection and snap frozen on dry ice. Samples were homogenized in buffer containing 0.2% SDS, protease and phosphatase inhibitors (Sigma), ran on a 4-12% SDS-PAGE gel (Biorad), electrotransfered on a PVDF (Millipore) and probed with the following antibodies: anti-LAMP-1: DSHB; anti-TFEB: Bethyl Laboratories; anti-GFP: Abcam; anti-GAPDH: Abcam.

**Western blotting for AKT2 in human samples**

Human brain samples were lysed in PBS containing 2% SDS. They were homozenized in a Heidolph homogenizer. Equal amounts of the lysate (according to the protein content quantified by BCA assay (Pierce)) were run on 4-12% BIS-TRIS gels (Invitrogen). The gel was blotted onto a nitrocellulose membrane (BioRad) and probed with the AKT2 specific antibody (Cell signalling). GAPDH antibody was purchased from meridian Science.

**Treatment of Arc/Swe APP primary neurons with AKT specific inhibitor**

Arc/Swe APP primary neurons were prepared from the cortex of E16 embryos and cultivated for 5-7 days in Neurobasal medium (Gibco) containing 1x B-27 (Invitrogen) and 1x L-Glutamin (Gibco). On the day of inhibitor treatment medium was replaced with fresh medium containing 10% Alamar blue (AbD Serotec) along with AKT Inhibitor IV (Calbiochem). The plates were incubated for 4 h, followed by Alamar blue measurements using SpecrtaMAX GeminiXS at an excitation wavelength of 544 nm and emission at 590 nm. Supernatant was collected in another plate and frozen at -20°C. Electrochemiluminescence assay was performed to determine the amount of Aβ and sAPPβ secreted by these cells. DMSO was used as control.

**Treatment of SHSY5Y cells with AKT specific inhibitor**

30,000 cells were seeded in each well of a 96 well plate (Nunc) a day prior to inhibitor treatment. On the day of inhibitor treatment medium was replaced with fresh medium along with AKT Inhibitor VIII (Calbiochem) or DAPT (Sigma) (positive control). 8 h after treatment, 10% Alamar blue (AbD Serotec) was added to the inhibitor treated samples. Alamar blue measurement were taken after 12 h using SpectraMAX GeminiXS (Molecular Devices) at an excitation wavelength of 544 nm and emission at 590 nm. Supernatant was collected and frozen at -20°C. Electrochemiluminescence assay was performed to determine the amount of Aβ secreted by these cells.

**Treatment of HeLa-sweAPP cells with AKT pathway specific inhibitors**

10,000 cells were seeded in each well of a 96 well plate (Nunc) a day prior to inhibitor treatment. On the day of inhibitor treatment medium was replaced with fresh medium containing either AKT inhibitor VIII (Calbiochem), AKT pathway inhibitor (PI-103) (Calbiochem), Perofosine (Selleckchem), Miltefosine (Merck), Everolimus (Selleckchem), AZD2014 (Selleckchem), Torin 1 (Axon Medchem) or KU-0063794 (Selleckchem). Cells were incubated with the inhibitors for 4 h. Alamar blue measurements were taken to determine cell viability using SpectraMAX GeminiXS (Molecular Devices) at an excitation wavelength of 544 nm and emission at 590 nm. Supernatant was collected and frozen at -20°C. Electrochemiluminescence assay was performed to determine the amount of Aβ secreted by these cells. DMSO was used as a negative control.

**Treatment of Human iPSC-derived Neurons with AKT pathway specific inhibitors**

Human iPSC-derived long-term self-renewing neuroepithelial stem cells (lt-NES; iPSCs derived from a 34-year-old healthy male donor, control 1 in (Mertens et al., 2013) were maintained in Dulbecco’s modified Eagle’s medium (DMEM)/F12, 2 mM L-glutamine, 1.6 g/l glucose, 0.1 mg/ml penicillin/streptomycin, N2 supplement (Life Technologies), B27 (1 µl/ml; Life Technologies), and fibroblast growth factor 2 (FGF2) and epidermal growth factor (EGF; both 10 ng/ml; R&D Systems) on tissue culture plates coated with poly-L-ornithine/laminin (both Sigma), and passaged every 3-4 days. Neuronal differentiation was induced by withdrawal of FGF2 and EGF in differentiation media (Neurobasal medium supplemented with B27 (1:50; Life Technologies) and DMEM/ F12 supplemented with N2 mixed at a 1:1 ratio and containing 300 ng/ml cyclic AMP) that was exchanged every other day. Akt pathway inhibitors, DAPT and DMSO solvent control were added to neuronal cultures differentiated for 4 weeks. On the day of treatment inhibitor were pre-diluted in differentiation media and added to the cultures for 4 h. Medium was changed and cultures were incubated for 14 h with the same treatment before supernatant and protein lysates were collected. The inhibitors used were either AKT inhibitor VIII at 10 µM concentration (Calbiochem), AKT pathway inhibitor (PI-103) at 10 µM concentration (Calbiochem), Perofosine 50 µM concentration (Selleckchem) or Torin-1 10 µM concentration (Axon Medchem). DMSO and DAPT (10 µM concentration) treatment were used as negative and positive controls respectively. Electrochemiluminescence assay was performed to determine the amount of Aβ secreted by these cells after normalizing the protein contents of the cell lysates using Pierce BCA protein assay Kit (Thermo Scientific). Alzheimer's disease human patients derived iPSC were treated with AKT pathway inhibitor (Calbiochem) (Concentration 10 µM) for 14 h or Torin-1 (Axon Medchem) (Concentration 10µM) for 14 h. Supernatant was collected and frozen at -20°C. After normalization of protein concentration using BCA electrochemiluminescence assay was performed to determine the amount of Aβ secreted by these cells. DMSO treatment was used as negative control.

**Lysotracker staining on serum starved HeLa cells**

10000 HeLa cells were seeded in each well of a 16 well chamber slide (Thermo scientific). 24 h after seeding, cells were serum starved for 18 h using minimal essential medium (MEM) (Gibco). Then, fresh medium (MEM) containing AKT pathway inhibitor (PI-103 Calbiochem) was added to the cells for 4.5 h. After 3 h pretreatment with AKT pathway inhibitor, cells were incubated for 1.5 h with 1 µM insulin (Sigma). Medium was replaced with fresh medium containing 200 nM Lysotracker Red DND-99 (Invitrogen) and incubated for 75 min followed by 15min fixation using 4% PFA. Fixed cells were stained with DAPI and mounted for microscopy. DMSO containing MEM was used as control.

**AKT pathway specific inhibitor treatment on HEK293 cells**

20000 HEK293 cells were seeded in each well of a pre-coated 16 well chamber slide. Pre-coating was done with polyornithine followed by fibronectin. 36 h after seeding, cells were serum starved using MEM (Gibco) for 10 h followed by amino acid starvation for 1 hr with DMEM devoid of amino acids (DMEM-aa) medium. Medium was replaced with fresh DMEM-aa medium containing AKT pathway inhibitor (Calbiochem) for 2.5 h. DMSO containing medium were used as control. After 2 h pretreatment with AKT pathway inhibitor, cells were incubated for 30 min with 1 μM insulin (Sigma).

Cells were then washed twice with PBS and fixed with 4 % PFA for 15 min at room temperature, permeabilized with PBS containing 0.2 % Triton X-100 for 10 min, followed by 1 h incubation with blocking solution containing 2% Bovine serum albumin (BSA) in PBS 0.2 % Triton X-100. Cells were incubated with primary antibody diluted in blocking solution for 12h at 4 °C (mouse anti-Lamp2, Santa Cruz Biotechnology, sc-18822) in 1:100 dilution and rabbit anti-TSC2 (Cell Signaling Technology, 4308P) in 1:150 dilutions. Cells were washed 5 times 5 min with PBS containing 0.2 % Triton X-100 and incubated with secondary antibody for 1 h at room temperature, diluted at 1:400 in blocking solution, stained with DAPI and mounted for microscopy.

For western blot 100000 HEK293 cells were seeded in each well of a pre-coated 12 well plate and treated with AKT Pathway inhibitor and insulin as previously described. Cells were lysed with TBS containing protease inhibitor and phosphatase inhibitor. β-actin was used as loading control.

**Treatment of Alzheimer's disease patients iPSc-derived Neurons with AKT pathway specific inhibitors**

Alzheimer's disease human patients derived iPSC were treated with AKT pathway inhibitor (Calbiochem) (Concentration 10μM) for 14 h or Torin-1 (Axon Medchem) (Concentration 10μM) for 14 h. Supernatant was collected and frozen at -20°C. After normalization of protein concentration using BCA (Pierce), electrochemiluminescence assay was performed to determine the amount of Aβ secreted by these cells. DMSO treatment was used as negative control.

**Treatment of Alzheimer's disease patients iPSC-derived Neurons with insulin**

Alzheimer's disease human patients derived iPSC were treated with 500 nM or 1 μM insulin. Supernatant was collected and frozen at -20°C. After normalization of protein concentration using BCA (Pierce), electrochemiluminescence assay was performed to determine the amount of Aβ secreted by these cells. DMSO treatment was used as negative control.

**Electrochemiluminescence detection Aβ from cell lysates after AKT pathway inhibition and insulin treatment**

10,000 HeLa-sweAPP cells were seeded in each well of a 96 well plate (Nunc) a day prior to inhibitor treatment. On the day of inhibitor treatment medium was replaced with fresh medium containing AKT pathway inhibitor (Calbiochem) for 3 h, following the treatment with AKT pathway inhibitor cells were treated with 1μM insulin (sigma) for 6 h. After insulin treatment cells were lysed in 35 μl of lysis buffer (1%NP40 and 0.1%SDS in TBS). Electrochemiluminescence assay was performed to determine the amount of Aβ present in cell lysates after protein normalization using BCA.

**Over-nutrition, Periodic fasting and Chronic fasting Diet**

3000 HeLa cells were seeded in each well of a 16 well chamber slide (Thermo scientific). For over nutrition diet medium was replaced with fresh full medium every 24 h. For periodic fasting medium was replaced with fresh medium, alternating between full medium and serum free medium every 24 h. For chronic fasting medium was replaced with fresh medium, alternating between between full medium and serum free medium every 48 h for 4 days. Medium was replaced with fresh medium containing 200 nM Lysotracker Red DND-99 (Invitrogen) and incubated for 75 min followed by 15min fixation using 4% PFA. Fixed cells were stained with DAPI and mounted for microscopy. DMSO containing MEM was used as control.

**Effect of insulin and starvation on protein aggregation**

6000 HeLa cells were seeded in each well of a 16 well chamber slide (Thermo scientific). 36 h after seeding, medium was replaced by DMEM containing serum or with out serum for 12 h. Then, fresh medium containing AKT pathway inhibitor (PI-103 Calbiochem), 1 μM insulin (Sigma) or AKT pathway inhibitor and insulin was added to the cells for 12 h. Medium was replaced with fresh medium containing 200 nM Lysotracker Red DND-99

(Invitrogen) and incubated for 75 min followed by 15 min fixation using 4% PFA. Fixed cells were stained with DAPI and ThioflainS and mounted for microscopy. DMSO containing medium was used as control.

**Perodic starvation in mice for Aβ measurment**

WT mice were subjected to periodic starvation for a period of 1 week. In this period mice were starved for 18 h (no food was provided). The control group had food all the time. Following starvation mice cortex were collected. Cortex was homogenized in TBS containing a protease inhibitor cocktail (Roche) in a Heidolph homogenizer. Aβ was extracted by 100000g ultracentrifugation for 1 h at 8°C in TBS containing 2% SDS using a TLA120.1 rotor (Beckman-Coulter). After normalizing for the protein content the amount of Aβ was measured by electrochemiluminescence assay.

**Perodic starvation in Arc/SweAPP Tg mice for Aβ measurment**

12 months old APP Arc, Sw mice were subjected to periodic starvation for a period of 40 days. In this period mice were starved for 18 h (no food was provided). The control group had food all the time. Following starvation mice cortex were collected. Cortex was homogenized in TBS containing a protease inhibitor cocktail (Roche) in a Heidolph homogenizer. Aβ was extracted by 100000g ultracentrifugation for 1 h at 8°C in TBS containing 2% SDS using a TLA120.1 rotor (Beckman-Coulter). After normalizing for the protein content the amount of Aβ was measured by electrochemiluminescence assay. In addition to Aβ, lysosomal markers and synaptic markers were analysed using western blot and following antibody were used: Cathepsin D (Abcam), Lamp2 (DSHB), PSD95 (Millipore), Synaptophysin (Synaptic systems), GAPDH (Meridian life sciences).

Brain slices from both starved and fed mice were stained with Congo-red (Sigma) used provided protocol. Briefly, tissues were rinse in water for 3x5 minutes and the placed in Alkaline Sodium Chloride Solution for 20 minutes. Further, they were stained in Alkaline Congo Red Solution for 20 minutes. Rinse in 3 changes of absolute ethanol, 50% ethanol, water and PBS. Then tissues were mounted using MOWIOL. For spine density measurement, Thy1 GFP marker was used.

**ThioflavinS staining**

Cells were washed twice with water for 5 min, followed by 4 min incubation with 1% ThioflavinS solution in 50% ethanol. Cells were then washed three times with 50% ethanol for 1 min. Then cells were washed with water twice 5 min followed by washed with PBS thrice 10 min. Then cells were mounted for microscopy. Same protocol was used to perform ThioS staining on mice brain slice.

**AKT inhibitor treatment on Primary neurons for Lysotracker and CLEAR gene**

Primary neurons were prepared from the cortex of E16 embryos and cultivated for 5-7 days in Neurobasal medium (Gibco) containing 1x B-27 (Invitrogen) and 1x L-Glutamin (Gibco). Neuron were treated with 10 μM AKT pathway inhibitor (Pi103-Calbiochem) for 2h. The medium was replaced with fresh medium containing 200 nM Lysotracker DND-99 (invitrogen) and incubate for 75 min. Following the neuron was washed with PBS and fixed by 4% PFA solution. Nucleus was stained by DAPI and mounted for microscopy. DMSO treatment as used as control.

For analysis of CLEAR gene neuron were treated with 10 μM AKT pathway inhibitor for 3h. Following cells were lysed trizol solution, RNA was isolated and cDNA was prepared using same method mentioned before. RT-PCR was performed for Lamp1, Lamp2, CatD and vATPase. GAPDH was used as control.

**Immunocytochemical characterization of iPSC-derived neuronal cultures**

For immunocytochemical characterization of iPSC-derived neuronal cultures, cells were washed with PBS and fixed with 4% paraformaldehyde (PFA, 10 min, RT), blocked in 0.1% Triton X-100 (Sigma) and 10% FCS in PBS, incubated with the primary antibodies (16 h, 4°C), washed with PBS, counterstained with secondary antibodies (1 h, RT) and DAPI and mounted with Mowiol. Primary antibodies and concentrations were as follows: SOX2 (1:500, R&D Systems), NESTIN (1:600, R&D Systems), DACH1 (1:100, Proteintech), beta III-tubulin (1:2000, Covance), MAP2ab (1:250, Chemicon), GFAP (1:1000, DakoCytomation), APP (1:1000, 4G8, Chemicon), PS1 (1:300, APS18, GeneTex), PHF1 (1:1000, gift from Peter Davies). Secondary antibodies were Alexa488 anti-ms, Alexa555 anti-ms, Alexa488 anti-rb and Alexa555 anti-rb (all 1:1000, Life Technologies).

**Treatment of neurons isolated from Wt mice with AKT specific pathway inhibitor**

Primary neurons isolated from the cortex of E16 embryos of Wt mice and cultivated for 5-7 days in Neurobasal medium (Gibco) containing 1x B-27 (Invitrogen) and 1x L-Glutamin (Gibco). 10,000 cells were cultivated in each well of a 96 well plate (Nunc). On the day of inhibitor treatment medium was replaced with fresh medium containing 10% Alamar blue (AbD Serotec) along with, Akt pathway Inhibitor (Calbiochem). The plates were incubated for 4 h, followed by Alamar blue measurements using SpecrtaMAX GeminiXS at an excitation wavelength of 544 nm and emission at 590 nm. Supernatant was collected in another plate and frozen at -20°C. Electrochemiluminescence assay was performed to determine the amount of Aβ and sAPPβ secreted by these cells. DMSO was used as a negative control.

**Influence of AKT inhibitor on gamma secretase processing of APP**

10,000 cells were seeded in each well of a 96 well plate (Nunc) a day prior to inhibitor treatment. On the day of inhibitor treatment medium was replaced with fresh medium containing either AKTinh I (AKT inhibitor VIII (Calbiochem) or AKTinh II (Pi-103 (Calbiochem) (lo-1μM and Hi-10μM) for 4h. DMSO was used as a negative control. Following the inhibitor treatment whole cell extracts were prepared using a lysis buffer (PBS pH 7.4 containing 1%NP40 and 0.1% SDS) supplemented with protease inhibitors. Extracts were subjected to SDS-PAGE using pre-cast gels (Invitrogen). Gel loading was normalized to total protein content in the cell extract (Using BCA assay). Proteins were transferred onto nitrocellulose membranes, which were then blocked with PBS containing 5% (w/v) dry skim milk for at least 1 h at room temperature. The membrane was then incubated with primary APP C terminal antibody (Epitomics), followed by the appropriate horseradish peroxidase-conjugated secondary antibody for at least 1 h at room temperature. Both antibodies were diluted in 5% milk/PBS 0.05% Tween-20. Immunoblotted proteins were detected using an enhanced chemiluminescence kit (Pierce). GAPDH (Meridian life sciences) was used as the loading control.

**Influence of AKT knock down on gamma secretase processing of APP**

Transfection complex containing the AKT siRNAs mix were prepared in Opti-mem medium (Invitrogen) by mixing 0.3 μL of Oligofectamine (Invitrogen) and 5 nM siRNAs. Hela swAPP cells at a density of 3500 cells/well were seeded in a 96 well plate after addition of transfection complexes. 72 h after transfection, Alamar blue measurements were taken using SpectraMAX GeminiXS (Molecular Devices) at an excitation wavelength of 544 nm and emission at 590 nm. Supernatant was collected and frozen at -20°C. The cells in the transfected plate were lysed with 50 μl of lysis buffer (PBS pH 7.4 containing 1%NP40 and 0.1% SDS) supplemented with protease inhibitors, incubated for 20 min on ice and stored at -20°C. Extracts were subjected to SDS-PAGE using pre-cast gels (Invitrogen). Gel loading was normalized to total protein content in the cell extract (Using BCA assay). Proteins were transferred onto nitrocellulose membranes, which were then blocked with PBS containing 5% (w/v) dry skim milk for at least 1 h at room temperature. The membrane was then incubated with primary APP C terminal antibody (Epitomics), followed by the appropriate horseradish peroxidase-conjugated secondary antibody for at least 1 h at room temperature. Both antibodies were diluted in 5% milk/PBS 0.05% Tween-20. Immunoblotted proteins were detected using an enhanced chemiluminescence kit (Pierce). GAPDH (Meridian life sciences) was used as the loading control.

**Electrochemiluminescence detection of Aβ42**

Electrochemiluminescence assay was performed to determine the amount of Aβ42 secreted by the siRNA transfected or inhibitor treated cells. Each well of the pre-coated plate was blocked using 150 μl TBST (Tris Buffered Saline containing Tween), containing 1% blocker A (MSD), for 1 hr at room temperature with shaking, followed by 3 times washing with 100μl TBST. After washing, 25 μl of the supernatant from the cells was added to each well along with 25 μl of detection antibody and incubated for 2 hr at room temperature with shaking. Wells were washed 3 times with 100μl TBST and detection was performed using 150 μl of 2x MSDT read buffer.

**Aβ extraction from WT and Arc/SweAPP Tg mice Tissues**

Cortex/Hippocampus was homogenized in TBS containing a protease inhibitor cocktail (Roche) in a Heidolph homogenizer. Soluble Aβ was separated from the insoluble fraction by 100000g ultracentrifugation for 1 h at 4°C using a TLA120.1 rotor (Beckman-Coulter). After normalizing for the protein content the amount of Aβ was measured by electrochemiluminescence assay.

**Assay for the secretion of Secreted Alkaline Phosphatase (SEAP)**

SEAP (Applied Biosystems) experiments were performed using manufacturer's guidelines. Briefly cells were transfected first with respective siRNA. 48 h after siRNA transfections they were again transfected with plasmid containing SEAP. 24 h after SEAP transfections the supernatant was collected and analysed for secreted alkaline phosphatase. Transfection with MEDGC was used as siRNA transfection control.

**Treatment with rapamycin**

20,000 HeLa-sweAPP cells were seeded in each well of a 96 well plate (Nunc) a day prior to inhibitor treatment. On the day of inhibitor treatment medium was replaced with fresh medium containing 10 μM Rapamycin (Sigma). Cells were incubated with Rapamycin for different time points. Alamar blue measurements were taken to determine cell viability using SpectraMAX GeminiXS (Molecular Devices) at an excitation wavelength of 544 nm and emission at 590 nm. Supernatant was collected and frozen at -20°C. Electrochemiluminescence assay was performed to determine the amount of Aβ secreted by these cells.

**AKT2 Plasmid Transfections for Electrochemiluminescence (ECL) Assay**

HeLa-sweAPP cells were transfected with either mock plasmid (pcDNA), plasmid expressing AKT2 using Lipofectamine 2000 reagent (Invitrogen), according to the manufacturer's protocol. 24 h after transfection the medium exchanged and conditioned medium was collected for 3 h and this was analyzed for Aβ40 using the Meso Scale Discovery Electrochemiluminescence (ECL) platform. Alamar blue measurements were taken to determine cell viability using SpectraMAX GeminiXS (Molecular Devices) at an excitation wavelength of 544 nm and emission at 590 nm.

**TFEB Plasmid Transfections for Electrochemiluminescence (ECL) Assay**

HeLa-sweAPP cells were transfected with either mock plasmid (pCDNA), plasmid expressing TFEB GFP or plasmid expressing TFEB ΔNLS GFP using Lipofectamine 2000 reagent (Invitrogen), according to the manufacturer's protocol. 24 h after transfection the medium exchanged and conditioned medium was collected for 3 h and this was analyzed for Aβ40 using the Meso Scale Discovery Electrochemiluminescence (ECL) platform. Alamar blue measurements were taken to determine cell viability using SpectraMAX GeminiXS (Molecular Devices) at an excitation wavelength of 544 nm and emission at 590 nm.

**TFE3 Plasmid Transfections for Electrochemiluminescence (ECL) Assay**

HeLa-sweAPP cells were transfected with either mock plasmid (pCDNA) or plasmid expressing TFE3 EGFP (Addgene:38120) using Lipofectamine 2000 reagent (Invitrogen), according to the manufacturer's protocol. 24 h after transfection the medium exchanged and conditioned medium was collected for 3 h and this was analyzed for Aβ40 using the Meso Scale Discovery Electrochemiluminescence (ECL) platform. Alamar blue measurements were taken to determine cell viability using SpectraMAX GeminiXS (Molecular Devices) at an excitation wavelength of 544 nm and emission at 590 nm.

**TFEB GFP Plasmid Transfections for Confocal microscopy**

5,000 Hela cells were seeded in each well of a 16 well microscopy slide (Nunc) a day prior to plasmid transfections. Cells were transfected with plasmid expressing TFE3 GFP using Lipofectamine 2000 reagent (Invitrogen), according to the manufacturer's protocol. 24 h after transfection the medium exchanged with fresh medium containing AKT pathway inhibitor (Pi103-Calbiochem) for 4 h. DMSO was used as a negative control. Following the inhibitor treatment cells were washed with PBS twice and fixed with 4% PFA for 15 mins. Fixed cells were stained with DAPI and mounted. The cells were visualized using the Confocal Laser Scanning Microscope Leica TCS SP8.

**Electrochemiluminescence detection Aβ from cell lysates**

Electrochemiluminescence assay was performed to determine the amount of Aβ present cells lysates. Each well of the pre-coated 96 plate was blocked using 150 μl TBST (Tris Buffered Saline containing Tween), containing 1% blocker A, for 1 h at room temperature with shaking, followed by 4 times washing in TBST. After washing, 25 μl of cell lysates was added to each well along with 25 μl detection antibody followed by incubation for 2 h at room temperature with shaking. Wells were washed 3 times with TBST and detection was performed using 150 μl of 2x MSDT read buffer.

**Lysotracker analysis**

HeLa-sweAPP were treated with 10μM of AKT pathway inhibitor compound or DMSO control for 6 h and subsequently incubated with 100nm LysoTracker-Red DND-99 dye (Invitrogen, Carlsbad, CA) for 1.5 h. Cells were then fixed with 4% PFA and stained with Alexa-Fluor633 Phalloidin. Images were acquired by using SP8 Confocal Microscopy (Leica) and processed by ImageJ Software.

**TFEB Plasmid Transfections for Confocal microscopy**

5,000 cells were seeded in each well of a 16 well microscopy slide (Nunc) a day prior to plasmid transfections. Cells were transfected with plasmid expressing TFEB GFP using Lipofectamine 2000 reagent (Invitrogen), according to the manufacturer's protocol. 24 h after transfection the medium exchanged with fresh medium containing AKT inhibitor VIII (Calbiochem) for 4 h. DMSO was used as a negative control. Following the inhibitor treatment cells were washed with PBS twice and fixed with 4% PFA for 15 mins. Fixed cells were stained with DAPI and mounted. The cells were visualized using the Confocal Laser Scanning Microscope Leica TCS SP8.

**Serum starvation and treatment of HeLa-sweAPP cells with insulin**

10,000 HeLa-sweAPP cells were seeded in each well of a 96 well plate (Nunc) a day prior to serum starvation. HeLa-sweAPP cells were serum deprived for 12 h followed by treatment with insulin in the presence or absence of AKT pathway inhibitor. Conditioned medium was collected for 3 h and this was analyzed for Aβ40 using the Meso Scale Discovery Electrochemiluminescence (ECL) platform. Alamar blue measurements were taken to determine cell viability using SpectraMAX GeminiXS (Molecular Devices) at an excitation wavelength of 544 nm and emission at 590 nm.

**Amino acid and Methionine starvation**

10,000 cells were seeded in each well of a 96 well plate (Nunc) a day prior to amino acid and methionine starvation. On the day of experiment, medium was replaced with fresh medium either devoid of amino acids or methionine. Cells were incubated for 3 hrs in the fresh medium. Alamar blue measurements were taken to determine cell viability using SpectraMAX GeminiXS (Molecular Devices) at an excitation wavelength of 544 nm and emission at 590 nm. Supernatant was collected and frozen at -20°C. Electrochemiluminescence assay was performed to determine the amount of Aβ secreted by these cells. DMEM (Invitrogen) containing 10%FBS (Gibco) and 1% Pen/strep (Invitrogen) was used as a negative control.

**Serum starvation and treatment of HeLa TFEB-GFP cells with insulin**

5,000 cells were seeded in each well of a 16 well microscopy slide (Nunc) a day prior to serum starvation. HeLa TFEB-GFP cells were serum deprived for 12 h followed by treatment with insulin in the presence or absence of AKT pathway inhibitor. DMSO was used as a negative control. Following the inhibitor treatment cells were washed with PBS twice and fixed with 4% PFA for 15 mins. Fixed cells were stained with DAPI and mounted. The cells were visualized using the Confocal Laser Scanning Microscope Leica TCS SP8. Cells were imaged for GFP (TFEB) and DAPI (Nucleus).

**Proteasomal activity assay**

200,000 HeLa cells were seeded in a 6 well plate (nunc). 24 h after seeding cells were subjected to serum starvation for 12 h or medium was replaced with fresh medium. The cells were then treated with AKT pathway inhibititor, epoxomicin (Calbiochem) or insulin for 6h. Cell were then lysed with lysis buffer containing 50mM HEPES, 5mM EDTA, 150mM NaCl and 1% TritonX-100. Proteasomal activity was measured using a commercially available kit (Millipore) following the product guidelines. DMSO was used as control.

**Treatment of primary neurons with astrocyte conditioned medium (acm)**

25000 astrocytes stably expressed different isoforms of human ApoE2, ApoE3 and ApoE4 were seeded in one well of 48 well plate. Next day medium were replaced with Neurobasal medium containing 1x B-27 and 1x L-Glutamin for 1 day. Following astrocyte conditioned medium was collected and centrifuged at 300g for 5 min to remove unwanted cells and cell debris. Then neuron medium was replaced with acm and incubated for 1 day. Neurons were incubated with medium containing 200 nM Lysotracker Red DND-99 (Invitrogen) for 75 min. After neurons were fixed with using 4% PFA at room temperature for 15 min followed by methanol for 5 min in -20 °C. Fixed neurons were stained with antibody against MAP2 and DAPI follwed by mounted for microscopy.

**Treatment of primary neurons with astrocyte conditioned medium (acm) and AKT-pathway inhibitor**

25000 astrocytes stably expressed different isoforms of human ApoE2, ApoE3 and ApoE4 were seeded in one well of 48 well plate. Next day medium were replaced with Neurobasal medium containing 1x B-27 and 1x L-Glutamin for 1 day. Following astrocyte conditioned medium was collected and centrifuged at 300g for 5 min to remove unwanted cells and cell debris. Then neuron medium was replaced with acm and incubated for 1 day. AKT-path way inhibitor was added for last 12 h. DMSO was used as control. Neurons were incubated with medium containing 200 nM Lysotracker Red DND-99 (Invitrogen) for 75 min. After neurons were fixed with using 4% PFA at room temperature for 15 min followed by methanol for 5 min in -20 °C. Fixed neurons were stained with DAPI followed by mounted for microscopy.

**Stereotactical delivery of AKT inhibitor, Rapamycin and AAV vectors**

3-4 months old transgenic Arc/SweAPP Tg mice were used in the *in vivo* inhibitor experiment. Mice were anesthetized with a ketamine and xylazine mix. 2 ml of DMSO (control), AKT inhibitor VIII, or Rapamycin was stereotactically injected into each hippocampus (coordinates from bregma were -2a/p, ±1.5 m/l, and -2.3 d/v from the skull). Animals were sacrificed 4 h after injection and the hippocampus sub dissected and snap frozen. Hippocampi were homogenized in TBS containing a protease inhibitor cocktail (Roche) in a Heidolph homogenizer. Soluble Aβ was separated from the insoluble fraction by 100000g ultracentrifugation for 1 h at 4°C using a TLA120.1 rotor (Beckman-Coulter). After normalizing for the protein content the amount of Aβ was measured by electrochemiluminescence assay.

3-4 month old WT mice received an intra-cortical injection of either AAV-GFP or AAV-TFEB (Decressac et al., PNAS 2013). Surgeries were performed under gaseous anaesthesia and analgesia (2% isoflurane in 2:1 oxygen/nitrous oxide), using a stereotaxic mouse frame (Stoelting Germany). Vector solutions were injected using a 5 µl Hamilton syringe fitted with a glass capillary (outer diameter of 250 µm). 1 ml of vector solution was injected in the cortex (coordinates from bregma were +0.6a/p, ±1.0 m/l, and -1.0 d/v from the dura). Animals were sacrificed 10 days after stereotaxic injection and tissue was processed for either Aβ measurements or western blot or histological analysis.

**Supernatant mixing experiment**

HeLa cells were treated with medium containing AKT pathway inhibitor (Calbiochem) (Concentration 10µM) for 4 h followed by collecting this conditioned medium and mixing it with conditioned medium obtained from from HeLa-sweAPP cells at a ratio of 1:1. Mixed medium was incubated for various time points at 37°C. Electrochemiluminescence assay was performed to determine the amount of Aβ and sAPPβ degraded in a time dependent manner. DMSO treatment was used as a negative control.

**Treatment with Torin 1**

10,000 HeLa-sweAPP cells were seeded in each well of a 96 well plate (Nunc) a day prior to inhibitor treatment. On the day of inhibitor treatment medium was replaced with fresh medium containing 5µM Torin1 (Axon Medchem). Cells were incubated with Torin 1 for different time points. Alamar blue measurements were taken to determine cell viability using SpectraMAX GeminiXS (Molecular Devices) at an excitation wavelength of 544nm and emission at 590nm. Supernatant was collected and frozen at -20°C. Electrochemiluminescence assay was performed to determine the amount of Aβ and sAPPβ secreted by these cells. DMSO treatment was used as negative control.

**Treatment of Cathepsin D with synthetic Aβ**

Recombinant human Cathepsin D (R&D systems) was reconstituted in water followed by dilution in assay buffer following the manufacturer's guidelines. Diluted Cathepsin D (0.625ng/µl) was incubated with synthetic Aβ (Meso Scale Discovery) at 37°C for 1 h. Following the incubation electrochemiluminescence assay was performed to determine the amount of degraded Aβ. Synthetic Aβ without Cathepsin D treatment was used as a negative control. Heat inactivated Cathepsin D (2.5ng/µl, Heat inactivation achieved by incubation at 95°C for 10 mins) was also incubated with synthetic Aβ (MSD) at 37°C for 3 h. Heat inactivated Cathepsin D did not degrade Aβ.

**Electron Microscopy.**

100,000 HeLa-sweAPP cells were seeded in a 3 cm dish (Nunc) a day prior to inhibitor treatment. On the day of inhibitor treatment medium was replaced with fresh medium containing AKT pathway inhibitor (Calbiochem) for 8 h. DMSO was used as a negative control. Cells were fixed in half-strength Karnovsky fixative (2% paraformaldehyde, 2.5% glutaraldehyde, 0.025% CaCl2.2H2O and 0.1 M sodium cacodylate buffer, pH 7.4), postfixed with 1 % $OsO_4$ and 1.5 % $K_3Fe(CN)_6$, dehydrated in ethanol and embedded in Epon. Ultrathin sections were stained with uranyl acetate and lead citrate.

**Supernatant transfer experiment**

HeLa cells were treated with medium containing either AKTinh I (AKT inhibitor VIII (Calbiochem) or AKTinh II (Pi-103 (Calbiochem) (lo-1µM and Hi-10µM) for 4 h followed by replacement of medium with conditioned medium from HeLa-sweAPP cells for 3 h. Alamar blue measurements were taken to determine cell viability using SpectraMAX GeminiXS (Molecular Devices) at an excitation wavelength of 544 nm and emission at 590 nm. Supernatant was collected and frozen at -20°C. Electrochemiluminescence assay was performed to determine the amount of Aβ and sAPPβ degraded by these cells. DMSO treatment was used as a negative control.

**Z-Score**

Z-Score for the screens is calculated using the formula: $Z = (x - \mu) / \sigma$

Z= Z-score, x=Average value of individual sample point, μ= mean of the population, σ= Standard deviation of the population

**Western blotting for S6K, TFEB GFP, TFEB S211-P, Phospho- S6k**

Cells were lysed in TBS + 1% Triton X-100 + protease (Complete, Roche) and phosphatase (PhosSTOP, Roche) inhibitor cocktails and insoluble material was removed by centrifugation for 10 min at 20,000 *g*. Immunoblotting was performed using standard methods (Bio-Rad Mini Protean system with transfer to nitrocellulose) and the antibodies defined above. Chemiluminescent detection (SuperSignal West Pico and Femto substrates, Pierce) of HRP signals was performed using a Versadoc imaging station (Bio-Rad).

The following primary antibodies were used in our experiments: anti–GFP-HRP (horseradish peroxidase; Miltenyi and Rockland Immunochemicals); tubulin (Sigma-Aldrich). Ribosomal protein S6 (total and phospho-S256/236) were from Cell Signaling Technology). The rabbit anti–phospho-TFEB serine 211 antibody was previously described (Petit et al., 2013).

**Microscopy and Image analysis**

Spinning disk confocal microscopy analysis of TFEB localization was determined as previously described (Petit et al., 2013).

**Statistical analysis**

Data were analyzed using Prism (GraphPad Software) and the tests specified in the figure legends.

**Filipin staining**

5 mg/ml stock Filipin (Sigma F-9765) in DMSO was prepared freshly. Fixed cells were washed three times with PBS and incubated 1h with 250 μg/ml Filipin solution in 10% FBS in

PBS. Following, cells were washed twice with PBS and mounted with MOWIOL. The cells were imaged using confocal microscopy at 405 excitation laser.

**Cathepsin D activity assay:**

Cathepsin D activity was analyzed by Cathepsin D Activity Assay Kit (Abcam). It is a fluorescence-based assay that utilizes a cathepsin-D substrate sequence labeled with MCA (7-methoxycoumarin-4-acetic acid). Cell lysates were incubated with the reaction mix at 37°C for 1-2 h. Cathepsin D in the lysate will cleave the synthetic substrate to release fluorescence, which can then be quantified using a fluorescence plate reader at Ex/Em = 328/460 nm.

**Magic red Cathepsin B activity assay**

Cathepsin B activity was analyzed by Magic Red Cathepsin B assay kit from Immunochemistry Technologies. The Magic Red reagent MR-RR2 (cresyl violet coupled to two copies of the amino acid sequence, arginine-arginine (RR)) enters the cell in a non-fluorescent state. If cathepsin B is active, the Magic Red substrate is cleaved and the cresyl violet fluorophone becomes fluorescent upon excitation. Cells were incubated in culture medium containing Magic Red for 1 h. Cells are washed with PBS and fixed with 4% PFA for 20 minutes. Nuclei were stained with DAPI and slide was mounted with cover slip. Cells were imaged with confocal microscope.

**EGF degradation assay**

10,000 cells were seeded in each well of a 16 well microscopy slide (Nunc). After 48 h cells incubated with 200 ng/ml EGF.Tetra Methyl Rohdomine for 10 min to look at uptake. To look at degradation capacity, medium was replaced fresh medium and chased for 30 min. Cells were then fixed with 4% PFA, nucleus was stained by DAPI and mounted. Cells were imaged with confocal microscope.

**Cholesterol loading and removal**

5,000 cells were seeded in each well of a 16 well microscopy slide (Nunc). Next day CHO WT cell were treated with 1.5 µg/ml U18666A in DMEM/F12 supplemented with 10% FBS media for 48 h for cholesterol loading. For cholesterol loading with MBC:cholesterol, 44 h after seeding WT cells were treated with DMEM/F12 with 10 % LPDS for 24 h followed by 100 µM MBC:cholesterol complex in 10 % LPDS media (filter sterile) for 4 h. For cholesterol removal, NPC1 null cells were treated with DMEM/F12 medium containing 10% LPDS for last 48h or treaded with 1mM MBC in DMEM/F12 medium containing 10% LPDS for 24 h. After the treatment cells were incubated with medium containing 200 nM LysoTracker Red DND-99 (Invitrogen) for 75 min for LysoTracker staining. After cells were fixed with using 4% PFA at room temperature for 15 min followed by methanol for 5 min in -20 °C. Fixed cells were stained with ThioS and filipin followed by mounted for microscopy.

**AKT-pathway inhibitor treatment**

10,000 cells were seeded in each well of a 16 well microscopy slide (Nunc). After 48 h cells incubated 10 μM AKT-pathway inhibitor of 4 h. After the treatment cells were incubated with medium containing 200 nM LysoTracker Red DND-99 (Invitrogen) for 75 min for LysoTracker staining. After cells were fixed with using 4% PFA at room temperature for 15 min followed by methanol for 5 min in -20 °C. Fixed cells were stained with ThioS and filipin followed by mounted for microscopy.

**NPC1 mouse model**

A mouse model of NPC disease, the BALB/cNctr-*Npc1*$^{N}$/+ (stock number 003092), was purchased from the Jackson Laboratory, Bar Harbor, Maine, USA. The mice were housed in accordance with the EU Directive 2010/63/EU for animal experiments. Mice were maintained on a 12 hour light/dark cycle with access to water and a standard mouse diet *ad libitum*. Female and male NPC1+/- mice were mated to generate NPC1-/- and NPC1+/+ (wt, control) mice as NPC1-/- mice are not fertile. Mice were genotyped according to already published protocols (http://jaxmice.jax.org/strain/ 003092.html). Mouse genotype was further confirmed by using sodium dodecyl sulphate-polyacrylamide gel electrophoresis (SDS-PAGE)/Western blotting, and immunoblotting with an anti-NPC1 antibody (ab134113; Abcam).

All mice were anesthetized and perfused with 0.9% saline solution. Brains were removed and hemisected; left hemispheres were immediately cut into cerebellum, hippocampus, cortex and remaining brain and snap-frozen in liquid nitrogen and stored at -80°C, whereas right hemispheres were immersion fixed in fresh 4% paraformaldehyde in phosphate-buffered saline (PBS) for one hour and then transferred to a 15% sucrose solution for 24 hours to ensure cryoprotection. On the next day, hemispheres were frozen in liquid isopentane and stored at -80°C until cutting.

**Preparation of mouse brain tissue lysates and immunoblotting**

Preparation of mouse brain tissue lysates for analyzing BACE1 proteolysis of its substrates was performed as in Kuhn PH et al. (2012). To generate soluble fractions, tissues were homogenized in 0.25% DEA, 100mM NaCl buffer, containing protease inhibitor cocktail (Complete, Roche Applied Science), in a glass homogenizer until tissue lysate appeared uniform. The centrifugation was carried out at 100 000 x g at 4°C for 30 minutes. After the centrifugation, tissue supernatants were collected, placed in fresh tubes and stored at -80°C freezer until further use. To generate membrane-bound fractions, the obtained tissue pellets were further homogenized in 1% Triton buffer [1% Triton X-100, 150mM NaCl, 50mM Tris-HCl, pH 7.4 and 2mM EDTA] - containing protease inhibitor cocktail (Complete, Roche Applied Science) by using a glass homogenizer. The lysates were then pressed through a 23-gauge needle by using a 1 ml syringe until tissue lysates appeared uniform. These tissue lysates were left on ice for 30 minutes incubation. Following this step, the lysates were centrifuged at 100 000 x g at 4°C for 30 minutes. After the centrifugation, tissue supernatants were collected, placed in fresh tubes and stored at -80°C freezer until further use.

**Immunohistochemistry of mouse brain cryosections**

The immersion-fixed and cryprotected frozen right hemispheres from NPC1+/+ and NPC1-/- mice were used to prepare 10 μm thick sagittal cryosections for histological analysis on a Leica CM 3050S cryotome. Sections were stored at -20°C until used for immunohistochemistry.

For immunocytochemistry, cryosections were briefly washed in PBS containing 0.5% Triton X-100 (PBS-T) and blocked in 5% goat serum in PBS-T for 1 hour. Sections were subsequently incubated with primary antibodies diluted in 5% goat serum in PBS-T overnight. After 3-5 hours incubation with secondary antibody (conjugated to Alexa Fluor 488, 594 or 647 from ThermoFisher Scientific) and Hoechst (Immunocytochemistry Technologies) was used to counterstain nuclei. Confocal images were acquired on an inverted laser scanning confocal microscope Leica TCS SCP8.

**Aβ uptake assay using HeLa swAPP conditioned medium in BV2 microglia**

3500 BV2 microglia cells were seeded in single well of a PDL-coated 96 well plate. After 56 h in parallel, supernatant was collected from HeLa swAPP cells for 4h in DMEM medium (devoid FBS). The condition medium was then centrifuged at 300g for 5 min to collect the supernatant. The supernatant was then divided in to two parts where one part was adjusted to 10% FBS medium.  After 60 h of seeding, BV2 microglia cells were incubated with HeLa swAPP conditioned medium for 12h along with Control (DMSO), 1μM AKT-pathway inhibitor, 1μM Torin-1  in 10% FBS medium and  serum starved condition. After 12h medium was collected for Aβ analysis. Cells were further incubated with in respective medium containing 10% Alamar blue to check cell viability. Aβ uptake by microglia was then analyzed based on remaining Aβ in the supernatant.

**Aβ40-alexa-647 uptake assay in BV2 microglia**

5000 BV2 microglia cells were seeded in single well of 96 well plate on a PDL-coated glass coverslip. After 48 h of seeding, BV2 microglia cells were incubated with Control (DMSO), 1 μM AKT-pathway inhibitor, 1 μM Torin-1  in 10% FBS medium and  serum starved condition for 12h. Following, cell were incubated with 500mM  alexa-647 tagged-Aβ40 (AnaSpec) for 2h. After 2h incubation cells were washed twice with PBS and fixed with 4% PFA for 20 min in RT. Nucleus were stained with DAPI and glass slide was mounted using MOWIOL. The up taken Aβ per cell was then imaged and quantified using confocal microscopy.

**Synaptosomes preparation**

First B6.Cg-Tg(Camk2a-cre)T29-1Stl mouse was crossed with B6.Cg-Gt(ROSA)26Sortm14(CAG-tdTomato)Hze mouse. 2 month old pups were then sacrificed using intracardiac perfusion, after deep anesthesia. The brains tissue was the homogenized in Syn-PER Reagent (Thermo Fisher) containing protease inhibitor on ice and synaptosomes were isolated according to the provided protocol. Briefly, first homogenate was at 1200 × g for 10 minutes at 4°C and pellet was discarded and supernatant was transferred to a new tube. A sample of the supernatant (homogenate) was saved for analysis. The supernatant was the further centrifuged at 15,000 × g for 20 minutes at 4°C. The supernatant was removed from the synaptosome pellet. The supernatant (cytosolic fraction) sample was stored for analysis. The synaptosomes pellet was suspended in Syn-PER reagent containing 5% DMSO (500μL for 400mg of brain tissue). The synaptosomes were aliquoted and stored -20°C for further use.

**Synaptosome uptake assay in BV2 microglia**

5000 BV2 microglia cells were seeded in single well of 96 well plate on a PDL-coated glass coverslip. After 48 h of seeding, BV2 microglia cells were incubated with Control (DMSO), 1 μM AKT-pathway inhibitor, 1 μM Torin-1 in 10% FBS medium and serum starved condition for 12h. Following, cells were incubated with tdTomato positive synaptosomes for 2h. After 2h incubation cells were washed twice with PBS and fixed with 4% PFA for 20 min in RT. Nucleus were stained with DAPI and glass slide was mounted using MOWIOL. The up taken Aβ per cell was then imaged and quantified using confocal microscopy.

**Mathematical modeling**

The mathematical model consists of two submodels: protein turnover and protein aggregation. The protein turnover model describes the following processes: (1) transport of extracellular amino acids ($AA_e$) into the cell, (2) transport of cytoplasmic amino acids ($AA_i$) out of the cell, (3) synthesis of protein (P), (4) degradation of protein by ubiquitin-proteosomal pathway (UPP), and (5) degradation of protein by lysosomal-autophagic pathway (LAP). Meanwhile, we used a published aggregation model for Aβ40/42 to simulate the formation of protein aggregates (Pa) from Aβ monomers (m) (Meisl et al., 2014). Figure A illustrates the cellular processes taken into account in the model. In the following, we describe in detail the derivation of model equations and the determination of kinetic parameters for each of the submodels.

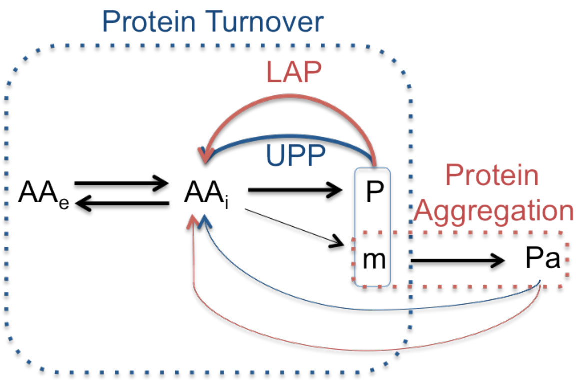


Figure A. Model of Protein Turnover and Aggregation.

Protein turnover submodel

The protein turnover submodel as shown in Figure A describes the mass balance of amino acids (extracellular $AA_e$ and cytoplasmic $AA_i$) and proteins (P and m). In the formulation of the model equations below, we do not differentiate between proteins (P) and monomers (m) of protein aggregates, and instead use only total proteins (also denoted by P). For parameter estimation, we used *in vitro* protein synthesis data from a previous study by Vabulas and Hartl (Vabulas and Hartl, 2005). In that study, the authors investigated the effects of nutrient restriction on protein turnover in HeLa cells. The amount of newly synthetized proteins was tracked by feeding the cells with radioactively labeled Methionin ($^{35}$S-Met) and detecting radioactivity of isolated intracellular proteins as a function of time (5, 10, 15 and 20 minutes). We employed data from the study, taken under four starvation conditions: (1) DMSO; (2) inhibition of UPP by epoxomicin; (3) DMSO after 6 hours of pre-starvation; (4) inhibition of UPP by epoxomicin after 6 hours of pre-starvation. Expoxomicin blocks the UPP degradation pathway (Vabulas and Hartl, 2005).

*Converting protein radioactivity to concentration:* At the beginning of these experiments (time point 0 minute), HeLa cells were put in deficient media and fed with the labeled methionine. At each time point, $5\times10^5$ cells were sampled from the culture for protein isolation and radioactivity measurements. As the data for protein amount were reported in the unit of $10^3$ cpm, we use the following formula for conversion into protein amount (nM):

$$1\,{}^{35}S-Met\;[nM] = \frac{10^{12}\left[\frac{fmol}{mmol}\right]}{SA_{{}^{35}S}-Met\;\left[\frac{Ci}{mmol}\right]\cdot 2.22\cdot 10^{12}\left[\frac{dpm}{Ci}\right]\cdot \varepsilon\left[\frac{cpm}{dpm}\right]\cdot V[\mu L]}\,1\,{}^{35}S-Met\;[10^3 cpm]$$

The specific activity $SA$ of $^{35}$S-Met is 1175 Ci/mmol (methionine.) and the efficiency of scintillator counter is reported to be 97% (Counter). In the study, the total volume V corresponds to the volume of $5\times10^5$ HeLa cells. Since the volume of a HeLa cell has been reported to be $2.6\times10^{-6}$μL (Milo, 2013), we used V = 1.3 μL. Figure B shows the converted protein data taken from the study of Vabulas and Hartl (Figure 1B in (Vabulas and Hartl, 2005)) which were used for the parameter estimation in this study.

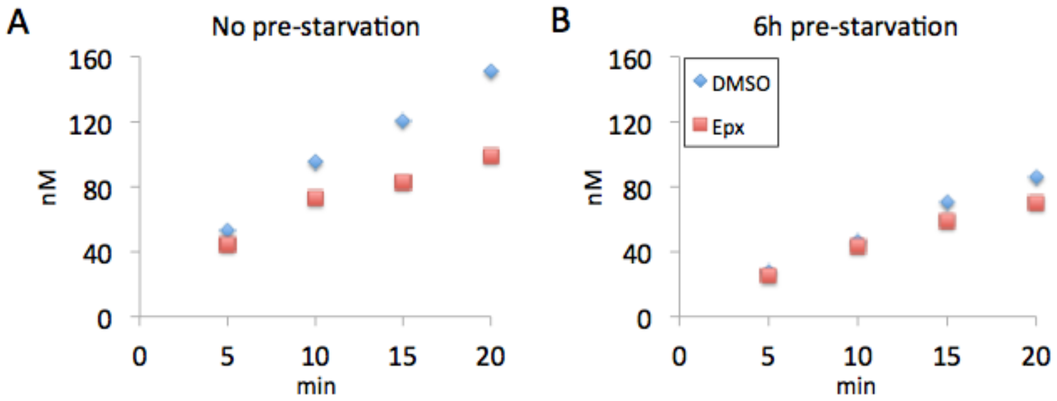


Figure B. Radioactively labeled methionine concentration (A) under no pre-starvation and (B) with 6 hours of pre-starvation. Epoxomicin (Epx) treatment blocks ubiquitin-proteosomal pathway of protein degradation. Adapted from (Vabulas and Hartl, 2005).

*Kinetics of Protein Turnover:* The exchange of amino acids across the cell membrane is modeled using mass action kinetics as follow:

Rate of transport of $AA_e$ into the cell: $r_i(AA_e) = k_i[AA_e]$

Rate of transport of $AA_i$ out of the cell: $r_e(AA_i) = k_e[AA_i]$

where $[AA_e]$ and $[AA_i]$ denote the concentrations of external and cytoplasmic amino acids, respectively, $k_i$ denotes the rate constant of amino acid transport into the cell, and denotes the rate constant of amino acid transport out of the cell. The protein degradation pathways are also modeled using mass action kinetics:

Rate of protein degradation by UPP: $r_{UPP}(AA_P) = k_{UPP}[AA_P]$

Rate of protein degradation by LAP: $r_{LAP}(AA_P) = k_{LAP}[AA_P]$

where $[AA_P]$ denotes the concentration of amino acids in proteins, and $k\_UPP$ and denote the rate constants of degradation by UPP and LAP, respectively.

In the study of Vabulas and Hartl (Vabulas and Hartl, 2005), the starvation condition involved 100-fold reduction of leucine (Leu) and methionine (Met) concentrations in the media at a ratio of 4:1 (Leu:Met). For this reason, we modeled the protein synthesis rate using a second-order mass action kinetics, assuming Leu and Met are the rate limiting amino acids, as follow:

Rate of protein synthesis: $r_s(Leu_i, Met_i) = k_s[Leu_i][Met_i]$

where $k_s$ denotes the rate constant of protein synthesis.

Here, we consider the synthesis of a typical protein of a size of 375aa, which is the median protein size in human cells (Brocchieri and Karlin, 2005). The average fractions of leucine and methionine in proteins are 0.0966 and 0.0242, respectively (ExPASy), which gives a Leu:Met ratio α in proteins of nearly 4 (3.99). This also means that there are 36.225 and 9.074 molecules of leucine and methionine in a typical protein, respectively. To convert from [$AA_P$] to the concentration of proteins [P], one needs to divide [$AA_P$] by the number of the corresponding amino acid in proteins (e.g., $[P] = \frac{[Leu_P]}{36.225}$).

*Model Formulation for Parameter Estimation*: We formulated the model based on the mass balance of Leu, Met and radioactively labeled Met, denoted by Met*. More specifically, we tracked the AA concentrations inside the cell ([$Leu_i$], [$Met_i$] and [$Met_i^*$]), as well as their concentrations in proteins ([$Leu_P$], [$Met_P$] and [$Met_P^*$]). We assumed that the external AA concentrations were not affected by the cell activity (i.e., the cells are supplied with excess external AA).

We investigated different possible regulations of protein transport and degradation according to the following considerations.

1. The transport of amino acids out of the cell membrane may be limited under starvation condition to maintain sufficient cytoplasmic amino acid level. In this case, the rate of transport of AA out of the cells is positively regulated by the external concentration of the corresponding AA. We implemented the rate of protein transport out of the cell as follows:

$$r_e(AA_i) = \begin{cases} k_e[AA_i] & \text{without regulation} \\ \dfrac{[AA_e]}{k_{U,e} + [AA_e]} k_e[AA_i] & \text{with regulation} \end{cases}$$

$$where\ AA = \{Leu, Met, Met^*\}$$

.

2. The degradation of proteins by UPP and LAP may be regulated by the amount of cytoplasmic amino acids (Vabulas and Hartl, 2005). We allowed different types of regulation on the rates of protein degradation by cytoplasmic AA. For each degradation pathway, we considered three options: no regulation, $AA_i$ upregulates degradation, or $AA_i$ inhibits degradation. Hence, we implemented the rate of protein degradation as follows:

$$r_{UPP}(AA_P) = \begin{cases} k_{UPP}[AA_P] & \text{without regulation} \\ \dfrac{[Leu_i] + [Met_i] + [Met_i^*]}{k_{U,UPP} + [Leu_i] + [Met_i] + [Met_i^*]} k_{UPP}[AA_P] & \text{with positive regulation} \\ \dfrac{k_{I,UPP}}{k_{I,UPP} + [Leu_i] + [Met_i] + [Met_i^*]} k_{UPP}[AA_P] & \text{with negative regulation} \end{cases}$$

$$r_{LAP}(AA_P) = \begin{cases} k_{LAP}[AA_P] & \text{without regulation} \\ \dfrac{[Leu_i] + [Met_i] + [Met_i^*]}{k_{U,LAP} + [Leu_i] + [Met_i] + [Met_i^*]} k_{LAP}[AA_P] & \text{with positive regulation} \\ \dfrac{k_{I,LAP}}{k_{I,LAP} + [Leu_i] + [Met_i] + [Met_i^*]} k_{LAP}[AA_P] & \text{with negative regulation} \end{cases}$$

where $AA = \{Leu, Met, Met^*\}$.

The protein turnover model consists of 6 ordinary differential equations as follows:

$$\frac{d[AA_i]}{dt} = -r_i(AA_e) + r_e(AA_i) - \frac{d[AA_P]}{dt} \qquad \text{where } AA = \{Leu, Met, Met^*\}$$

$$\frac{d[Leu_P]}{dt} = r_s(Leu_i, Met_i + Met_i^*) - r_{UPP}(Leu_P) - r_{LAP}(Leu_P)$$

$$\frac{d[Met_P]}{dt} = \frac{1}{\alpha} r_s(Leu_i, Met_i) - r_{UPP}(Met_P) - r_{LAP}(Met_P)$$

$$\frac{d[Met_P^*]}{dt} = \frac{1}{\alpha} r_s(Leu_i, Met_i^*) - r_{UPP}(Met_P^*) - r_{LAP}(Met_P^*)$$

By implementation of all possible combinations of regulatory interactions mentioned earlier, we formulated 18 possible ODE models (2 combinations of transport across membrane × 3 combinations of UPP degradation × 3 combinations for LAP degradation). The more regulatory interactions a model has, the larger is the number of parameters to be estimated. Table A gives the descriptions of the models used for the parameter estimation.

**Table A. Model Description and Identification**

| ***Model ID*** | ***Description*** |
|---|---|
| *1* | *no regulation (core protein turnover model)* |
| *2* | *UPP is negatively regulated by* $AA_i$ |
| *3* | *UPP is positively regulated by* $AA_i$ |
| *4* | *LAP is negatively regulated by* $AA_i$ |
| *5* | *LAP and UPP are negatively regulated by* $AA_i$ |
| *6* | *LAP is negatively regulated by* $AA_i$*; UPP is positively regulated by* $AA_i$ |
| *7* | *LAP is positively regulated by* $AA_i$ |
| *8* | *LAP is positively regulated by* $AA_i$*; UPP is negatively regulated by* $AA_i$ |
| *9* | *LAP and UPP are positively regulated by* $AA_i$ |
| *10* | *Protein export rate is positively regulated by* $AA_e$ |
| *11* | *Protein export rate is positively regulated by* $AA_e$*; UPP is negatively regulated by* $AA_i$ |
| *12* | *Protein export rate is positively regulated by* $AA_e$*; UPP is positively regulated by* $AA_i$ |
| *13* | *Protein export rate is positively regulated by* $AA_e$*; LAP is negatively regulated by* $AA_i$ |
| *14* | *Protein export rate is positively regulated by* $AA_e$*; LAP and UPP are negatively regulated by* $AA_i$ |
| *15* | *Protein export rate is positively regulated by* $AA_e$*; LAP is negatively regulated by* $AA_i$*; UPP is positively regulated by* $AA_i$ |
| *16* | *Protein export rate is positively regulated by* $AA_e$*; LAP is positively regulated by* $AA_i$ |

| | |
|---|---|
| *17* | *Protein export rate is positively regulated by $AA_e$; LAP is positively regulated by $AA_i$; UPP is negatively regulated by $AA_i$* |
| *18* | *Protein export rate is positively regulated by $AA_e$; LAP and UPP are positively regulated by $AA_i$* |

$$[AA_i] = [Leu_i] + [Met_i] + [Met_i^*];\ [AA_e] = [Leu_e] + [Met_e] + [Met_e^*]$$

*Parameter estimation:* For the parameter estimation of the protein turnover model, we employed radioactivity data of HeLa cells fed with $^{35}$S-Met at the onset of the measurements, as shown in Figure B. The parameter estimation was formulated as a constrained optimization over all rate constants, except  (see below for more detail). The optimal parameter values corresponded to the minimum of squares of the difference between model concentration predictions $[Met_P^*]$ and the experimental data in Figure B, summed over all time points. In generating model predictions of $[Met_P^*]$, we used the steady state concentrations of cytoplasmic AAs and proteins, calculated as follows:

1. Given the external concentrations of AAs and under a steady state condition, we can calculate the concentrations of cytoplasmic AAs by multiplying the external concentrations ($[Leu_e] = 8{\times}10^4$ nM and $[Met_e] = 2{\times}10^4$ nM) by the ratio $\frac{k_i}{k_e}$.
2. Meanwhile, the average protein concentration in human cells has been reported to be 0.2 g/ml (Milo, 2013). Assuming an average protein size of 50 kDa, 100 pmol of proteins is equal to 5 μg (GenScript). We used this conversion to obtain an average concentration of intracellular proteins of $4{\times}10^6$ nM, and therefore $[Leu_p] = 36.225{\times}4{\times}10^6 = 1.45{\times}10^8$ nM and $[Met_P] = 9.074{\times}4{\times}10^6 = 3.63{\times}10^7$ nM.
3. Based on the concentrations above and again under a steady state condition, we can compute the protein synthesis rate constant. For example, for the protein turnover model without any regulatory interaction, this is given by

$$k_s = \frac{[Leu_p]}{[Leu_i]\cdot[Met_i]}\left(k_{UPP} + k_{LAP}\right)$$

1. Since labeled methionine was fed only at the start of the experiment, the concentrations of labeled methionine as free cytoplasmic AAs and those in proteins are set to 0 (i.e. $[Met_i^*] = [Met_P^*] = 0$ nM).

We simulated the four starvation conditions performed in the study of Vabulas and Hartl (Vabulas and Hartl, 2005) according to the following procedure: (please refer to Figure B for the experimental data)

1. DMSO: We simulated the model using the steady state concentrations of cytoplasmic AAs and proteins from above, while setting the external concentrations of $[Leu_e] = 8\times10^2$ nM, $[Met_e] = 2\times10^2$ nM and $[Met_e^*]$ =42.6 nM (taken from experimental conditions reported in Supplementary Material of (Vabulas and Hartl, 2005)).
2. Epoxomicin: The simulation of starvation with epoxomicin treatment started with the same initial conditions as for DMSO in point 1. But, we set the rate constant of UPP set to 0 (i.e. $k_{UPP} = 0$) to describe the inhibition of UPP by epoxomicin.
3. Pre-starvation + DMSO: We implemented the pre-starvation by simulating using the steady state initial concentrations determined above with external concentrations $[Leu_e] = 8\times10^2$ nM, $[Met_e] = 2\times10^2$nM and $[Met_e^*] = 0$ nM for 6 hours. Subsequently, we simulated the labeling experiments starting from the final concentrations from the pre-starvation run but with $[Met_e^*] = 42.6$ nM.
4. Pre-starvation + epoxomicin: The simulation of pre-starvation was performed as in point 3 above. During the simulation of labeling experiments, we again started with the final concentrations from the pre-starvation run and setting $[Met_e^*] = 42.6$ nM. Here, we further set the rate constant of UPP to 0 (i.e. $k_{UPP} = 0$).

The following constraints were applied during parameter estimation:

1. The degradation rate constants have been reported to be between $3\times10^{-4}$ to $3\times10^{-3}$ $min^{-1}$ (Boisvert et al., 2012; Gerner et al., 2002). We therefore constrained the sum of the degradation rate constants, multiplied by any regulatory scaling factor evaluated at steady state condition, to be within the above limits.

2. We constrained the ratio $\frac{k_i}{k_o}$ to be between 9 to 50, such that given the concentrations of external leucine and methionine under normal conditions in the experiment $[Leu_e] = 8\times10^4$ nM and $[Met_e] = 2\times10^4$ nM (Vabulas and Hartl, 2005) $[Leu_i]$ would range between $7.2\times10^5$ and $4\times10^6$ nM and $[Met_i]$ would range between $1.8\times10^5$ and $1\times10^6$ nM. In the literature, the cytoplasmic leucine and methionine have been reported to be $7.3\times10^5$ nM and $1.9\times10^5$ nM, respectively (Piez and Eagle, 1958).

We solve the constrained optimization and obtain the optimal parameter values of each model in Table A using the global optimization toolbox SSmGO (SSmGO). The results of the parameter estimation are given in a supplementary Excel file. For avoiding data overfitting, we also calculated the Akaike criterion (AIC) (Akaike, 1974), as shown in Figure C below. AIC provides a measure of the trade-off between goodness-of-fit to experimental data and the number of parameters. A lower AIC value is preferable. Model 1 gave the lowest AIC and the model fitting is shown in Figure D.

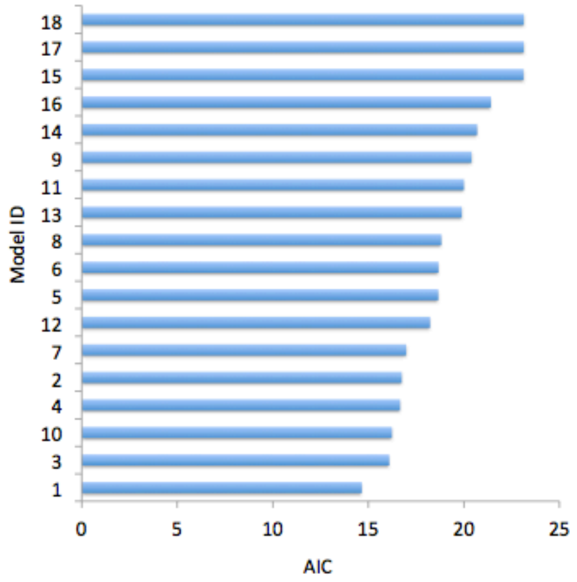


Figure C. Akaike index (AIC) in 18 protein turnover models.

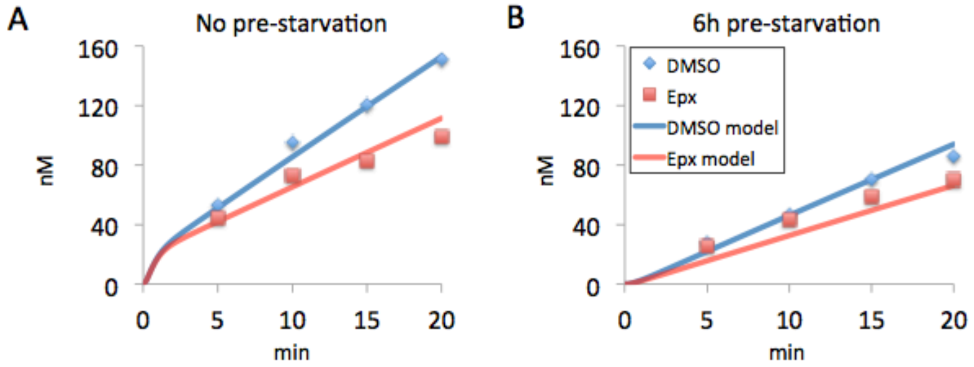

Figure D. Parameter estimation of Model 1 for experiments (A) without pre-starvation and (B) with 6 hours of pre-starvation. Data adapted from (Vabulas and Hartl, 2005).

Protein aggregation submodel

For the simulation of Aβ aggregation, we adopted the kinetic models of Aβ40 and Aβ42 aggregation, recently proposed by Meisl and colleagues (Meisl et al., 2014). The aggregation model took into account the following processes: (1) primary nucleation of protein aggregates, (2) secondary nucleation of aggregates and (3) elongation or growth of aggregates. The rates characterizing the aggregation kinetics are given below.

1. Primary nucleation: $n_1(m) = k_n[m]^2$

   where $k_n$ denotes the rate constant of primary nucleation and $[m]$ denotes the protein monomer concentrations.

2. Secondary nucleation: $n_2(m, P_a) = k_2 \frac{[m]^2}{1+[m]^2/K_M^2[P_a]}$

   where $k_2$ denotes the rate constant of secondary nucleation, $K_M$ denotes the concentration constant and $[P_a]$ denotes the concentration of proteins in aggregates.

3. Elongation: $\varpi(m, nP_a) = 2k_+[m][nP_a]$

   where $k_+$ denotes the aggregate elongation rate constant and $[nP_a]$ denotes the number concentration of aggregate particles.

The kinetic parameters for Aβ40 and Aβ42 aggregation were obtained from data fitting, and are reproduced in Table B.

**Table B. Kinetic parameters for the aggregation of Aβ40 and Aβ42 (Meisl et al., 2014).**

| | ***Aβ40*** | ***Aβ42*** |
|---|---|---|
| $k_n$ ***(nM$^{-2}$×min$^{-1}$)*** | *$1.2\times10^{-22}$* | *$1.8\times10^{-20}$* |
| $k_2$ ***(nM$^{-2}$×min$^{-1}$)*** | *$1.8\times10^{-13}$* | *$6\times10^{-13}$* |
| $K_M$ ***(nM)*** | *$6\times10^{3}$* | *$1.2\times10^{4}$* |
| $k_+$***(nM$^{-1}$×min$^{-1}$)*** | *$1.8\times10^{-2}$* | *0.18* |

In formulating the model combining protein turnover and aggregation, we assumed that protein aggregates degrade at a slower rate than free proteins by a constant factor of $\rho_d < 1$. We also assumed that Aβ monomers are produced at a constant fraction $\rho_{A\beta} < 1$ of the total protein synthesis. The full model equations describing the protein turnover and aggregation of wild-type (WT) Aβ monomer are given by:

$\frac{d[Leu_i]}{dt} = r_i(Leu_e) - r_e(Leu_i) - r_s(Leu_i, Met_i) + r_{UPP}(Leu_P + Leu_m) + r_{LAP}(Leu_P + Leu_m) + s_{Leu}\rho_d[r_{UPP}(Pa) + r_{LAP}(Pa)]$

$\frac{d[Leu_P]}{dt} = (1 - \rho_{A\beta})r_s(Leu_i, Met_i) - r_{UPP}(Leu_P) - r_{LAP}(Leu_P)$

$\frac{d[Leu_m]}{dt} = \rho_{A\beta}r_s(Leu_i, Met_i) - r_{UPP}(Leu_m) - r_{LAP}(Leu_m) - s_{Leu}\varpi(m, nP_a)$

$\frac{d[Pa]}{dt} = \varpi(m, nP_a) - \rho_d\big(r_{UPP}(Pa) - r_{LAP}(Pa)\big)$

$\frac{d[nPa]}{dt} = n_1(m) + n_2(m, P_a) - [\![\rho_d(r]\!]_{UPP}(nPa) + r_{LAP}(nPa)\big)$

where $s_{Leu} = 36.225$ (number of molecules of Leu in P), $[m] = \frac{[Leu_m]}{36.225}$ and $[Met_i] = \frac{1}{\alpha}[Leu_i]$. While the number of leucine in Aβ40 and Aβ42 is not equal to $s_{Leu}$, the formulation above maintains the ratio of protein and monomer concentrations at the constant fraction $\rho_{A\beta}$.

In simulating the aggregation of mutant Aβ40 and Aβ42, we multiplied the rates of nucleation and elongation of protein aggregates by a factor of $\rho_{mut} = 7.5$. Correspondingly, the model equations for mutant Aβ aggregation are given by:

$\frac{d[Leu_i]}{dt} = r_i(Leu_e) - r_e(Leu_i) - r_s(Leu_i, Met_i) + r_{UPP}(Leu_P + Leu_m) + r_{LAP}(Leu_P + Leu_m) + s_{Leu}\rho_d[r_{UPP}(Pa) + r_{LAP}(Pa)]$

$\frac{d[Leu_P]}{dt} = (1 - \rho_{A\beta})r_s(Leu_i, Met_i) - r_{UPP}(Leu_P) - r_{LAP}(Leu_P)$

$\frac{d[Leu_m]}{dt} = \rho_{A\beta}r_s(Leu_i, Met_i) - r_{UPP}(Leu_m) - r_{LAP}(Leu_m) - s_{Leu}\rho_{mut}\varpi(m, nP_a)$

$\frac{d[Pa]}{dt} = \rho_{mut}\varpi(m, nP_a) - \rho_d\big(r_{UPP}(Pa) - r_{LAP}(Pa)\big)$

$\frac{d[nPa]}{dt} = \rho_{mut}n_1(m) + \rho_{mut}n_2(m, P_a) - [\![\rho_d(r]\!]_{UPP}(nPa) + r_{LAP}(nPa)\big)$

*Simulations of the Effects of Nutrition and Insulin:* The simulations of nutritional and insulin effects started from steady state condition. Here, the steady state concentrations of the AA, protein and aggregates were obtained by simulating the aggregation model for a period of 365 days with external concentrations $[Leu_e] = 8\times10^4$ nM and $[Met_e] = 2\times10^4$ nM. The concentrations of the cytoplasmic AA, proteins and protein aggregates were initially set to 0 nM.

For studying the effects of increased nutrition and insulin, we simulated the protein turnover and aggregation model starting from the steady state condition, where we increased the external concentrations by 3 fold: $[Leu_e] = 2.4\times10^5$ nM and $[Met_e] = 6\times10^4$ nM, increased the protein synthesis rate constant $k_s$ by 5 fold, increased the rate constant of UPP degradation $k_{UPP}$ by 30%, and decreased the rate constant of LAP degradation $k_{LAP}$ by 80%. On the other hand, for simulating the effects of starvation, we only decreased the external concentrations by 3 fold: $[Leu_e] = 2.67\times10^4$ nM and $[Met_e] = 6.67\times10^3$ nM, keeping the kinetic parameters unchanged.

We performed model simulations of the effects of nutrition and insulin using each of the top 5 protein turnover models (models 1, 2, 3, 4, and 10 in Table A), corresponding to the 5 lowest AIC values (see Figure C). Furthermore, we simulated the models using different parameters $\rho_d$ = {0.05, 0.1, 0.2} and $\rho_{A\beta}$ = {$5\times10^{-7}$, $10^{-6}$, $2\times10^{-6}$} for WT and mutant Aβ40 and Aβ42. The complete results of the model simulations can be found in Supplementary Files: Simulations of Ab40 Aggregation.pdf and Simulations of Ab42 Aggregation.pdf. The results using the best protein turnover model (model 1) with $\rho_d = 0.1$ and $\rho_{A\beta} = 10^{-6}$ were given in the paper. The rapid accumulation of protein aggregates due to nutrition and insulin increase was more pronounced for mutant Aβ42 than mutant Aβ40, which was not surprising as Aβ42 could aggregate more readily than Aβ40. The simulations of the five models are in general agreement in regard to the effects of nutrition and insulin on WT and mutant Aβ aggregation. Also, the same general observations can be made for different parameters $\rho_d$ and $\rho_{A\beta}$. Therefore, model-based findings in our study are insensitive to the assumption on regulatory structure and model parameters.

**Bioinformatics analysis of tissue-specific human transcriptome**

Figure E illustrates the workflow of the analysis of the Genotype-Tissue Expression (GTEx) transcriptomics data. More details of each of the steps in this workflow are provided below.

Differential gene expression analysis with age (Step 1 in Figure E)

The expressions of 52'576 transcripts in 29 human post-mortem tissue samples were obtained from the GTEx project (GTEx v3 dataset). Genes that were not expressed in any of the samples were excluded from the analysis. We also eliminated genes with no expression in more than 95% of the tissue samples, reducing the number of gene transcripts to 32'217. We determined the genes whose expression varied significantly with ageing across different tissues using a linear mixed effect model (LMM) (Mele et al., 2015). In the LMM analysis, the gene expression was modeled with age as a covariate, tissues as a fixed effect, and individuals as a random effect. We found 2'108 genes with age-dependent expression across all tissues in the dataset (FDR<0.05, Benjamini and Hochberg procedure), referred to as the ageing genes.

Trait – Gene dataset (Step 2 in Figure E)

We curated human traits - gene associations from GWAS studies and Mendelian mutations. GWAS data were downloaded from the NHGRI GWAS Catalog (https://www.ebi.ac.uk/gwas/) and the PheGenI database (http://www.ncbi.nlm.nih.gov/gap/phegeni). Both databases provided associations between single nucleotide polymorphisms (SNPs) and complex traits and diseases in human. Only intragenic SNPs associated with human traits and diseases were included in this study. The disease-gene Mendelian associations were obtained from OMIM, a database of human phenotypes with known associations with gene mutations. Subsequently, we curated the combined human traits-genes associations manually to remove redundant human trait and disease names. The gene names were finally cross-checked and corrected against the human genome organization (HUGO) gene symbols. Our human traits-gene association database comprised a total of 3'358 traits and 7'257 genes.

Gene Ontology Semantic Similarity (GOSS) score measurement (Step 3 in Figure E)

Gene Ontology semantic similarity (GOSS) scores for GO biological process terms were calculated between ageing – represented by the ageing genes above – and human traits in the curated database above, using the Lin scoring function *mgosim* in the R-package *GOSemSim (Yu et al., 2010)*. The GOSS score provides a measure of similarity between genes or gene groups based on the biological knowledge embedded in the GO structure. In order to establish the statistical significance of the computed GOSS scores, we performed a randomization analysis by generating 1'000 random sets of genes with the same cardinality (number) as the ageing genes, using a uniform random sampling from the list of 32'217 genes above. We computed the GOSS scores between each random ageing gene set and a reshuffled list of human traits-gene associations, where the genes were randomly permuted. The randomization analysis produced 1'000 random GOSS scores for each ageing - human trait pair, based on which we computed the statistical significance. The set of human traits and the corresponding genes (as ENTREZ Gene ID) that had significant similarity with ageing is given in Table M1 below ($p < 0.05$).

Pathway enrichment analysis (Step 4 in Figure E)

The pathway enrichment analysis of the union of the genes from human traits with statistically significant similarity with ageing, was performed using the Reactome pathway database (https://www.reactome.org).

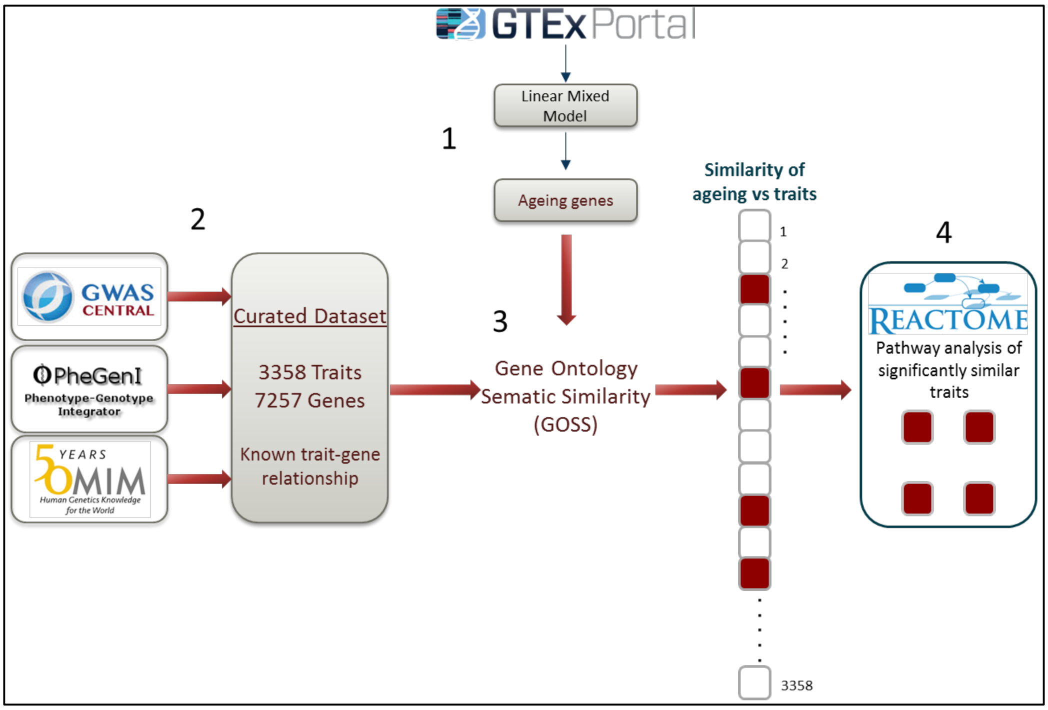


Figure E: A workflow of the bioinformatics analysis of ageing. *Step 1*: Identification of genes that are differentially expressed with age using a LMM analysis of transcriptomics data from the GTEx project. *Step 2:* Curation of a compendium of human traits – gene associations from GWAS, OMIM and PheGenI databases. *Step 3*: Gene Ontology Semantic Similarity (GOSS)

analysis for identifying human traints and genes that are similar in terms of GO biological processes with the ageing genes. Step 4: Reactome pathway enrichment analysis.

Table M1: Ageing-similar human traits and genes.

| ***Human traits*** | ***ENTREZ Gene ID*** |
|---|---|
| *Acampomelic campomelic dysplasia* | *10011* |
| | *6662* |
| *Adenocarcinoma of lung* | *7917* |
| | *7015* |
| | *5071* |
| | *4758* |
| | *2064* |
| | *1136* |
| | *673* |
| *Abdominal aortic aneurysm* | *153090* |
| | *4035* |
| *Aplastic anemia* | *4683* |
| | *3458* |
| | *2149* |
| *Type 5D distal arthrogryposis* | *9427* |
| *Lethal arthrogryposis with anterior horn cell disease* | *2733* |
| *Ataxia-telangiectasia* | *472* |
| | *185* |
| *Atrichia with papular lesions* | *55806* |
| *Basal cell carcinoma* | *8643* |
| | *7157* |
| | *6608* |
| | *5921* |
| | *5727* |
| *Basal ganglia calcification* | *6575* |
| | *5159* |
| | *5155* |
| *Budd-Chiari syndrome* | *3717* |
| | *2153* |
| *Campomelic dysplasia with autosomal sex reversal* | *10011* |
| | *6662* |
| *CARASIL syndrome* | *5654* |
| *Cardiovascular function* | *83478* |
| | *7481* |
| | *6336* |
| | *6331* |

| | |
|---|---|
| | *4211* |
| | *857* |
| *Colorectal cancer* | *201163* |
| | *79695* |
| | *27030* |
| | *26585* |
| | *25788* |
| | *23476* |
| | *10395* |
| | *8313* |
| | *7157* |
| | *7099* |
| | *7097* |
| | *6790* |
| | *6714* |
| | *5795* |
| | *5782* |
| | *5426* |
| | *5424* |
| | *5320* |
| | *5290* |
| | *5157* |
| | *4893* |
| | *4163* |
| | *4092* |
| | *4091* |
| | *3911* |
| | *2261* |
| | *2033* |
| | *1630* |
| | *1499* |
| | *999* |
| | *701* |
| | *673* |
| | *595* |
| | *581* |
| | *324* |
| | *207* |
| *Cowden syndrome* | *100144748* |
| | *23476* |
| | *6392* |
| | *6390* |
| | *6389* |
| | *5728* |
| | *5290* |

| | |
|---|---|
| | *207* |
| *Crohn disease-associated growth failure* | *3569* |
| | *3082* |
| *Autosomal recessive cutis laxa* | *30008* |
| *Transient neonatal cyanosis* | *3048* |
| *Digenic deafness* | *2707* |
| *Permanent neonatal diabetes mellitus* | *6833* |
| | *3630* |
| | *2645* |
| *Ehlers-Danlos syndrome, type VIIC* | *9509* |
| *Gastric cancer risk after H. pylori infection* | *3557* |
| | *3553* |
| *Glioma* | *51750* |
| | *7157* |
| | *7015* |
| | *5728* |
| | *3417* |
| *Hepatocellular cancer* | *23476* |
| | *23095* |
| | *8312* |
| | *7157* |
| | *5290* |
| | *5157* |
| | *3482* |
| | *1499* |
| | *841* |
| | *717* |
| *Histiocytosis-lymphadenopathy plus syndrome* | *55315* |
| *Resistance to HIV-1* | *7098* |
| | *6356* |
| | *6347* |
| *Susceptibility to intracranial hemorrhage in brain cerebrovascular malformations* | *3569* |
| | *3082* |
| *Susceptibility to Kaposi sarcoma* | *3569* |
| | *3082* |
| *Lymphangioleiomyomatosis* | *7249* |
| | *7248* |
| *Somatic B-cell non-Hodgkin Lymphoma* | *472* |
| | *185* |
| *Mantle cell Lymphoma* | *472* |
| | *185* |
| *Susceptibility to cerebral malaria* | *7124* |
| | *3383* |
| | *948* |

| | |
|---|---|
| *Martsolf syndrome* | *25782* |
| | *25* |
| *Mast syndrome* | *51324* |
| *Microvascular complications of diabetes* | *7422* |
| | *6648* |
| | *5444* |
| | *4072* |
| | *3557* |
| | *3077* |
| | *2056* |
| | *1636* |
| *Muscular dystrophy* | *9499* |
| *Somatic myelofibrosis* | *10019* |
| | *3717* |
| | *811* |
| *Myocardial infarction* | *375056* |
| | *221692* |
| | *55759* |
| | *7804* |
| | *7292* |
| | *6597* |
| | *5687* |
| | *4973* |
| | *4049* |
| | *3690* |
| | *2730* |
| | *2729* |
| | *2099* |
| | *1952* |
| | *1636* |
| | *348* |
| *Nanophthalmos* | *83552* |
| *Postmenopausal Osteoporosis* | *1278* |
| | *811* |
| | *799* |
| *Ovarian cancer* | *63908* |
| | *25976* |
| | *23476* |
| | *22800* |
| | *10801* |
| | *8631* |
| | *5290* |
| | *4978* |
| | *4758* |
| | *2064* |

| | |
|---|---|
| | *1499* |
| | *999* |
| | *207* |
| *Piebaldism* | *6591* |
| | *3815* |
| *Proliferative vasculopathy and hydraencephaly-hydrocephaly syndrome* | *55640* |
| *Primary pulmonary hypertension* | *4093* |
| | *4091* |
| | *3777* |
| | *857* |
| *Systemic juvenile rheumatoid arthritis* | *4282* |
| | *3569* |
| | *3082* |
| *Saethre-Chotzen syndrome* | *7291* |
| | *2263* |
| *Squamous cell carcinoma* | *8795* |
| | *5728* |
| | *3621* |
| *Somatic T-cell prolymphocytic leukemia* | *472* |
| | *185* |
| *Follicular thyroid carcinoma* | *9562* |
| | *5728* |
| | *4893* |
| | *3265* |
| *Autosomal dominant woolly hair* | *121391* |

**Immunohistochemistry for human PM tissue and confocal microscopy**

Formalin fixed and paraffin-embedded post-mortem superior temporal lobe of 4μm thickness was provided from the Edinburgh Brain Bank and the Bristol Brain Bank. Tissue was deparaffinised with xylene, followed by rehydration using serial ethanol dilutions of ascending water content (100% EtOH, 90% EtOH, 70% EtOH, 50% EtOH, and $H_2O$). For antigen retrieval, the tissue slides were pressure cooked for 3 minutes in citric acid (Vectalabs, H-3300). Subsequently, they were blocked with 10% Normal Donkey Serum and 0.3% TritonX-100 for 1 hour, before incubating with the primary antibodies overnight in block solution. The following primary antibodies were used: goat Iba1 (1:500; ab5076), mouse CD68 (1:100; M0876), rabbit Synapsin 1 (1:500; AB1543P). After the overnight incubation, primaries were washed off, followed by secondary antibody incubation for 1 hour (1:500; highly cross adsorbed H+L IgG; ThermoFisher Scientific) and DAPI counterstain (10 minutes, 1ug/ml).

Washes were done with diluted phosphate buffered saline (PBS pH 7.4) and PBS-0.3% TritonX-100.

Using a Leica TCS8 confocal microscope at the 63x oil immersion objective, 20 images were acquired per case from all areas of the grey matter using a random sampling approach. An ImageJ macro was generated for z-stacking, thresholding, masking, and multiplying images to provide the percent area where CD68 and Synapsin 1 co-localise. All slides were blinded for image acquisition and analysis by an independent researcher.

Six groups of participants were chosen based on varying disease status. Namely, non-demented controls, non-demented with hyperlipidaemia, non-demented controls with Type 2 Diabetes (T2D), Alzheimer's disease (AD) with no metabolic disease, AD with hyperlipidaemia, and lastly AD with T2D. All groups are age-matched.

**Ethical approval for human tissue:**

Use of human tissue for post-mortem studies has been reviewed and approved by the Edinburgh Brain Bank ethics committee and the ACCORD medical research ethics committee, AMREC (ACCORD is the Academic and Clinical Central Office for Research and Development, a joint office of the University of Edinburgh and NHS Lothian, approval number 15-HV-016). The Edinburgh Brain Bank is a Medical Research Council funded facility with research ethics committee (REC) approval (16/ES/0084).

### References


Akaike, H. (1974). A new look at the statistical model identification. Automatic Control, IEEE Transactions on *19*, 716-723.

Boisvert, F.-M., Ahmad, Y., Gierliński, M., Charrière, F., Lamont, D., Scott, M., Barton, G., and Lamond, A.I. (2012). A quantitative spatial proteomics analysis of proteome turnover in human cells. Mol Cell Proteomics *11*, M111.011429.

Brocchieri, L., and Karlin, S. (2005). Protein length in eukaryotic and prokaryotic proteomes. Nucleic acids research *33*, 3390-3400.

Counter, E.o.B.L.S.

ExPASy.

GenScript.

DePristo MA, Banks E, Poplin R, et al. A framework for variation discovery and genotyping using next-generation DNA sequencing data. Nat Genet 2011;43:491-8.

Gerner, C., Vejda, S., Gelbmann, D., Bayer, E., Gotzmann, J., Schulte-Hermann, R., and Mikulits, W. (2002). Concomitant determination of absolute values of cellular protein amounts, synthesis rates, and turnover rates by quantitative proteome profiling. Molecular & Cellular Proteomics *1*, 528-537.
Li H, Durbin R. Fast and accurate short read alignment with Burrows- Wheeler transform. Bioinformatics 2009; 25:1754-60.
McKenna A, Hanna M, Banks E, et al. The Genome Analysis Toolkit: a MapReduce framework for analyzing next-generation DNA sequencing data. Genome Res 2010; 20:1297-303.
Meisl, G., Yang, X., Hellstrand, E., Frohm, B., Kirkegaard, J.B., Cohen, S.I.A., Dobson, C.M., Linse, S., and Knowles, T.P.J. (2014). Differences in nucleation behavior underlie the contrasting aggregation kinetics of the Aβ40 and Aβ42 peptides. Proc Natl Acad Sci U S A *111*, 9384-9389.
Mele, M., Ferreira, P.G., Reverter, F., DeLuca, D.S., Monlong, J., Sammeth, M., Young, T.R., Goldmann, J.M., Pervouchine, D.D., Sullivan, T.J., *et al.* (2015). Human genomics. The human transcriptome across tissues and individuals. Science *348*, 660-665.
methionine., S.a.o.l.
Milo, R. (2013). What is the total number of protein molecules per cell volume? A call to rethink some published values. Bioessays *35*, 1050-1055.
Petit, C.S., Roczniak-Ferguson, A., and Ferguson, S.M. (2013). Recruitment of folliculin to lysosomes supports the amino acid-dependent activation of Rag GTPases. J Cell Biol *202*, 1107-1122.
Piez, K.A., and Eagle, H. (1958). The free amino acid pool of cultured human cells. The Journal of biological chemistry *231*, 533-545.
SSmGO.
Purcell S, Neale B, Todd-Brown K, Thomas L, Ferreira MAR, Bender D, Maller J, Sklar P, de Bakker PIW, Daly MJ & Sham PC (2007). PLINK: a toolset for whole-genome association and population-based linkage analysis. American Journal of Human Genetics, 81.
Vabulas, R.M., and Hartl, F.U. (2005). Protein synthesis upon acute nutrient restriction relies on proteasome function. Science *310*, 1960-1963.
Wang K, Li M, Hakonarson H. ANNOVAR: functional annotation of genetic variants from high-throughput sequencing data. Nucleic Acids Res 2010; 38(16):e164.
Yu, G., Li, F., Qin, Y., Bo, X., Wu, Y., and Wang, S. (2010). GOSemSim: an R package for measuring semantic similarity among GO terms and gene products. Bioinformatics *26*, 976-978.
